# Ecological axes of skull diversification in a massive vertebrate radiation

**DOI:** 10.64898/2026.06.19.733456

**Authors:** Elizabeth Christina Santos, Rose Faucher, Aintzane Santaquiteria, JoJo West, Jonathan W. Armbruster, Carole Baldwin, Thaddaeus J. Buser, Kent Carpenter, Juan Martín Diaz de Astarloa, Melissa Rincon-Sandoval, Samantha M. Gartner, Shih-Pin Huang, Jin-Koo Kim, Hernán López-Fernández, Nathan Lujan, Nicholas Mandrak, Masaki Miya, Mayara Pereira Neves, Melanie M. Paquin, John J. Pogonoski, Emily M. Troyer, Mark Westneat, William T. White, Edward O. Wiley, Giorgio Carnevale, Guillermo Ortí, Christopher M. Martinez, Lily C. Hughes, Ricardo Betancur-R, Kory Evans, Dahiana Arcila

## Abstract

Eupercarian spiny-rayed fishes are one of the largest vertebrate radiations, rivaling mammals and occupying nearly every aquatic habitat. We present a densely sampled, time-calibrated phylogenomic framework for Eupercaria, supporting a revised classification, combined with the largest cranial phenomics dataset for fishes. Habitat and trophic ecology make independent, complementary contributions to skull shape. Most species cluster around a conserved generalized architecture, the Percomorph Pile, from which one clade of pufferfishes, anglerfishes, butterflyfishes, and surgeonfishes repeatedly invaded novel morphospace; exceptionally high rates on its deep branches indicate that rapid skull evolution arose early in this clade. Freshwater lineages converge on the ancestral condition, reflecting late arrival into systems occupied by older otophysans, whereas durophages show the greatest disparity and converge on derived forms. Cranial diversity was partitioned among subclades during the Cretaceous and later within them across the Cenozoic, showing that clade-level differences in evolutionary rates and ecological opportunity jointly shaped skull diversification.

## Main text

Morphological diversification over deep time can be shaped by multiple ecological axes acting independently or in concert^1–5^. Ecological opportunity, in the form of the availability of unfilled niches through habitat colonization, the loss of competitors, or the evolution of key innovations, is thought to release lineages from stabilizing selection and accelerate morphological divergence^1,6^. Two axes through which such opportunity operates are habitat and trophic ecology. Habitat transitions expose lineages to new physical environments and competitive landscapes^7^, whereas dietary specialization directly impacts the ability to acquire energy resources which can shape the biomechanics of feeding structures, as prey size, hardness, and capture mode impose selection that can act independently of the environment a species inhabits^8–12^. Whether these two axes make independent or redundant contributions to phenotypic diversification has rarely been tested with sufficient phylogenetic, ecological, and morphological scope^13–16^.

Clades distributed across diverse environments allow for powerful comparative analyses of evolution in response to ecological variation, and these investigations become more compelling when independent transitions to the same habitat or trophic niche reveal recurring patterns. However, such insights depend on a robust phylogenetic framework. Spiny-rayed fishes (acanthomorphs) are one of the most diverse vertebrate groups on the planet, encompassing ∼10% of all extant vertebrate species^17^. Their evolutionary history has resisted resolution for decades^17–21^, and the group was once dubbed the “bush at the top” of the fish tree of life^22,23^. Much of this diversity, and much of the difficulty, is concentrated in the series Eupercaria (perch-like fishes), which contains >6,000 species and 160 families^20^—a level of diversity rivaling that of all living mammals^24^. Eupercarians have radiated into a diverse range of forms, from beaked wrasses and pufferfishes that dart around coral reefs^8,25,26^ to the unusual predatory anglerfishes and gelatinous snailfishes inhabiting the crushing depths of the ocean^14,27^. Eupercarians have colonized nearly every aquatic habitat on the planet^28^, ranging from freshwater streams^29–31^ to the freezing waters of the Antarctic^32^. This diversity has long confounded phylogenetic estimation because of seemingly rapid early divergences among lineages (Data S1), and multi-locus molecular phylogenies published within the past decade have not converged on a stable set of interrelationships within Eupercaria^20,35^ (Fig. 1), earning this clade the title of the “new bush at the top”^20^. The timeline of eupercarian diversification is likewise uncertain, with some studies arguing for explosive diversification near the Cretaceous–Paleogene (K–Pg) boundary^21^ and others for a more gradual radiation over the Late Cretaceous and Cenozoic^17,35^.

**Fig. 1.**
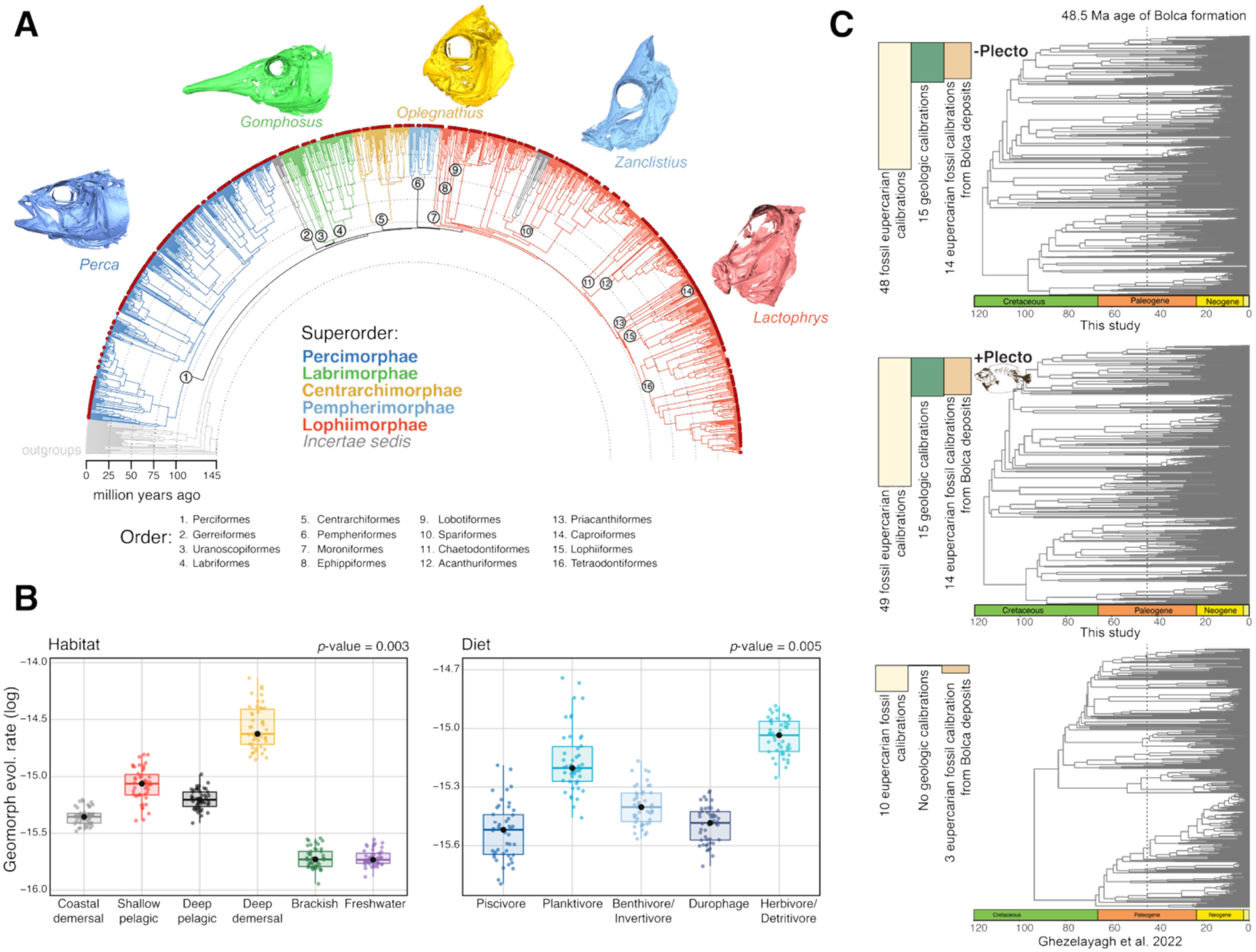
Phylogenomics, divergence times, and evolutionary rates across Eupercaria. (**A**) Phylogeny of Eupercaria based on 1,095 concatenated exon markers estimated with IQ-TREE (-Plecto variant shown). Clades are colored by superorder, with all 16 orders enumerated (Data S1). Red circles indicate the 571 species with a CT scan representative (table S6). (**B**) Evolutionary rates of cranial shape differ significantly among habitat categories (left) and diet categories (right); boxplots summarize rates across 46 tree topologies (*p* = 0.005 for both predictors). (**C**) Top: IQ-TREE chronogram excluding †Plectocretacicoidea from the calibration scheme (−Plecto). Middle: IQ-TREE chronogram including †Plectocretacicoidea as a calibration (+Plecto). Bottom: divergence estimates from ref.^17^. Numbers indicate the total fossil calibrations and the subset derived from Bolca for each tree. The dashed line marks the Monte Bolca Lagerstätte (∼48.5 Ma). Ingroup fossil calibrations are shown (Data S2). Time scales across geologic time were created using the deeptime R package^93^. Silhouettes are not to scale.

Here, we present a new phylogenomic framework for eupercarian fishes with extensive taxonomic sampling (1,050 species) (tables S1–S3), paired with time-calibrated phylogenies (tables S4–S5) and the largest phenomic dataset^2,36^ for fishes to date: 3D geometric morphometric data from micro-CT scans of 571 species, including all families in the phylogeny^37,38^. We focused on skulls because they integrate the mechanical demands of feeding with the ecological pressures of habitat^8,12,39^, perform a wide range of critical functions^15,40^, and evolve rapidly in response to selection^41^. Using this framework, we test whether habitat, trophic ecology, or their combination best explains patterns of skull shape variation, evolutionary rate, and morphological disparity across aquatic environments from freshwaters to the deep sea. Our results show that habitat and trophic ecology make independent, complementary contributions to cranial diversification and that access to morphological novelty is phylogenetically restricted, implicating clade-specific differences in evolutionary potential as a key modulator of diversification.

### Phylogenomics resolves eupercarian relationships and clarifies classification and diversification timing

We performed phylogenetic inference on a dataset comprising 1,095 exonic loci^42^ (Fig. 1A; figs. S1–S4). Taxonomic sampling included 133 eupercarian families (82.6% of known families), 484 genera (39.4%) and 1,050 (15.1%) species (tables S1–S2). Relationships based on concatenated exons were congruent with analyses using ultra conserved elements (UCEs)^17^, with most higher-level nodes shared between trees (Fig. 1A; fig. S5), suggesting that phylogenomics has delivered unprecedented resolution to eupercarian relationships^20,22,23,43^. In contrast, multi-locus phylogenies^44,45^ show lower higher-level congruence^35,20^ (figs. S6–S7). Concatenation-coalescent disagreement was also common, consistent with incomplete lineage sorting during rapid radiations.

This improved resolution (Fig. 1A) prompts reclassification of Eupercaria^20^. Both exon and UCE topologies^17^ support five major clades (fig. S5). A classification recently proposed^17,46,47^ recognized these as orders, triggering cascading nomenclatural changes. We instead recognize them as superorders, retaining traditional order-level names where monophyly is supported (Fig. 1A). Our classification recognizes five superorders, 16 orders, and 42 suborders (table S3). The phylogenetic bush that once characterized Eupercaria^20^ is now limited to 12 families within Lophiimorphae (=Acanthuriformes *sensu* ref.^17^). Discussion of this new scheme, its justification, and remaining areas of phylogenetic uncertainty are provided in Data S1.

The timeline of eupercarian diversification has remained contentious^16,17,20,38,50^, with debate over whether Late Cretaceous assemblages represent standing lineage diversity or sampling gaps^51^. We estimated divergence times using 49 fossil and 15 geologic calibrations (Data S2, Tables S4–S5); all major lineages originated during the Cretaceous (Fig. 1C). Crown ages for Eupercaria were consistent whether †Plectocretacicoidea was included (∼117 Ma) or excluded (∼116.7 Ma), unlike previous studies where it inflated estimates^25^. Recent work supports its placement as a stem tetraodontiform^52,53^, bridging a Late Cretaceous ghost lineage (Data S2). Even without †Plectocretacicoidea, substantial support for a revised timeline of diversification comes from 14 ingroup fossil calibrations from the Eocene Monte Bolca deposits (∼48.5 Ma^54^). Many of these calibrations are placed within deeply nested positions across several Eupercarian clades, such as Lophiiformes^55^ (Fig. 1C), providing a geochronologically precise anchor that is helping to reshape divergence estimates for many lineages, particularly fossiliferous reef-associated groups^54^. For example, crown ages for Labridae and Lutjanidae are herein revised upward by ∼10–20 Ma compared to previous estimates^34,56^. To accommodate dating uncertainty, we generated 46 alternative chronograms varying in molecular datasets, calibration scheme, and tree estimation approach (four based on all-genes topologies and 42 on gene-subset topologies; see Methods).

### Skull shapes cluster around a generalized perch-like shape

Eupercarian skull shape diversity (table S6, fig. S8) was characterized by a densely occupied cluster around the mean shape and inferred ancestral form, corresponding to a generalized perch-like skull (Fig. 2). Specifically, this perch-like shape consists of a terminal mouth, a large laterally placed orbit, skull of intermediate depth and width, premaxilla with elongated ascending process, maxilla that deepens posteriorly, and preopercle widely separated from the orbit. All eupercarian superorders overlap at least partially with this region of morphospace (fig. S9), which is best typified by the species closest to the centroid, the snapper *Lutjanus aratus* (also the species closest in shape to the inferred ancestor of Eupercaria). We hereafter refer to this region as the ‘Percomorph Pile’ because of the high density of taxa previously assigned to the order Perciformes^43,55^ (table S3). The tendency for many fish families to share this generalized shape has long contributed to confusion over their phylogenetic relationships.

**Fig. 2.**
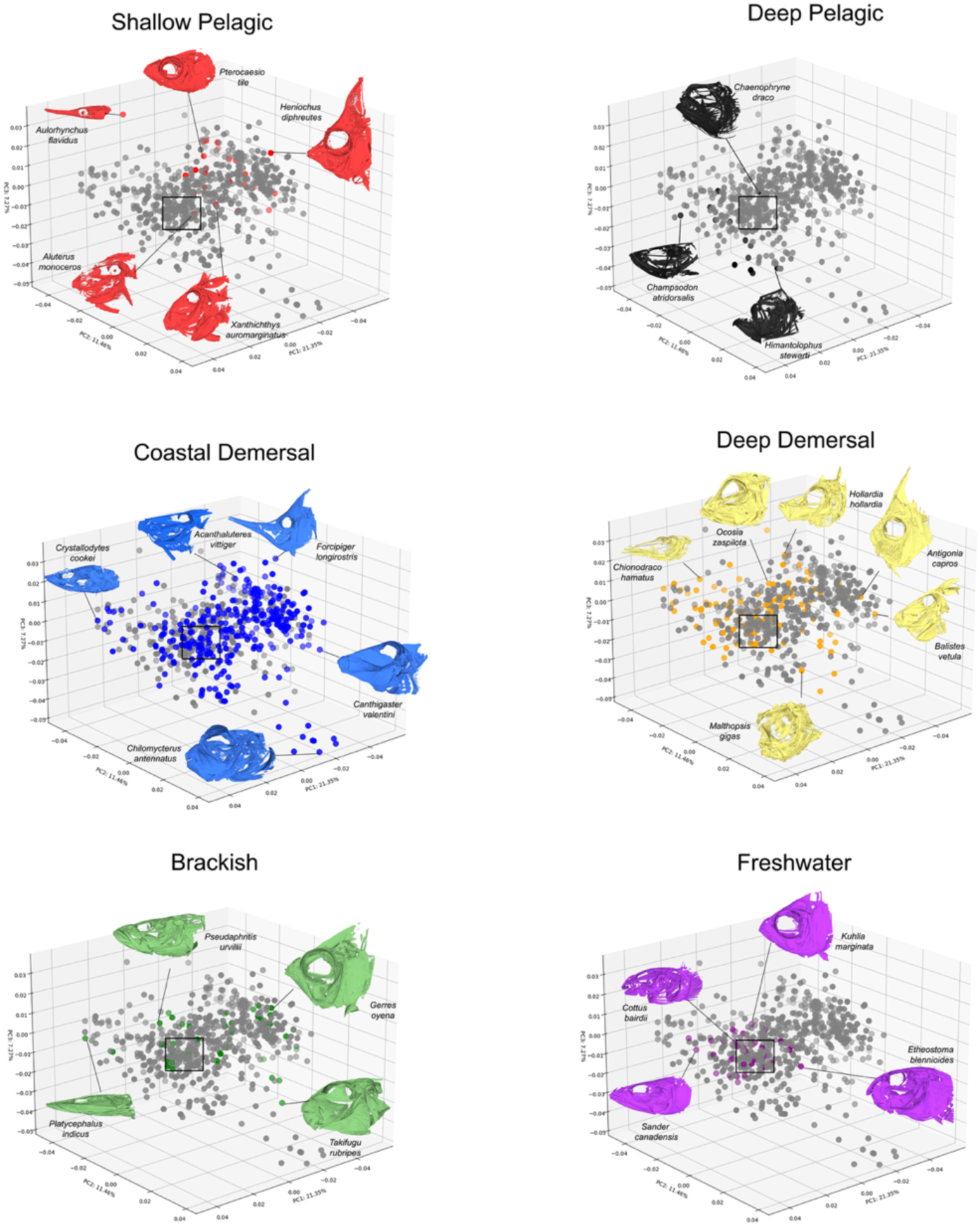
Cranial morphospace of eupercarian fishes partitioned by habitat. Morphospace plots partitioned by habitat for 571 species along three principal component axes of skull shape variation. Species belonging to each habitat category are highlighted in color; all remaining species are shown in gray for reference. Insets show representative skulls from different regions of morphospace. The square indicates the region of highest species density (the “Percomorph Pile”). Habitat explained 4.1% of cranial shape variance (phylogenetic Procrustes ANOVA; *p* = 0.003).

Departures from the Percomorph Pile reveal the full range of cranial forms in Eupercaria (Fig. 2). The main axis of morphospace (PC1, 21.3%) reflects a gradient from elongate, narrow-gaped skulls with posteriorly placed orbits to short, deep skulls with wide gapes and enlarged jaws. PC2 (11.5%) captures changes in skull elongation and mouth position, ranging from elongate skulls with slightly subterminal mouths to short-snouted forms with terminal mouths. PC3 (7.2%) reflects variation in skull width and depth, separating broader, deeper skulls (for example, many beaked tetraodontiforms^25^) from narrower, shallower forms such as the butterflyfish *Forcipiger longirostris*.

Certain eupercarian clades occupy especially broad regions of skull morphospace. Tetraodontiformes (puffers, molas, filefishes, and triggerfishes) span both extremes of PCs 2 and 3 (Fig. 2, fig. S10), ranging from narrow tube-snouted skulls (*Halimochirurgus*) to wide, robust forms with crushing tooth plates (e.g., *Diodon*). The living species furthest from the inferred ancestral eupercarian shape is a filefish (*Acanthaluteres vittiger*). Labrids also occupy a broad expanse of morphospace, including bird wrasses (*Gomphosus*), beaked parrotfishes (*Sparisoma*), and burrowing wrasses (*Iniistius*) (fig. S9). Although many Perciformes remain concentrated in the Percomorph Pile, the order as a whole extends widely through morphospace, including the distinctive Antarctic icefish radiation^32^ (*Bathydraco*; fig. S9).

### Habitat shapes cranial diversification but rarely defines a single skull phenotype

Habitat was significantly associated with skull shape in Eupercaria across all 46 phylogenies (R² = 0.022–0.190, all *p* < 0.005; mean R² = 0.066). Ancestral-state reconstructions did not identify a single unambiguous root habitat across the four all-genes trees. Root reconstructions alternated between brackish and coastal demersal, with brackish favored in two trees and coastal demersal favored in the other two. Across trees, mean root posterior was highest for brackish (0.553), followed by coastal demersal (0.350) and shallow pelagic (0.098), whereas deep demersal, deep pelagic, and freshwater received no support at the root. These habitats differed strongly in skull disparity, but most did not correspond to a single diagnostic cranial form. Species from nearly all habitats overlapped at least partly with the Percomorph Pile, indicating that each habitat contains some species with generalized, perch-like skulls (Figs. 2, 3). For example, the snapper genus *Lutjanus* occurs in several habitats with limited cranial divergence among species, whereas the highly derived bird wrasse (*Gomphosus varius*) and the more generalized red snapper (*Lutjanus campechanus*) are both coastal demersal fishes but occupy very different regions of morphospace. Thus, habitat captures broad ecological setting, but most habitats contain a mixture of generalized and highly divergent skull forms.

Freshwater eupercarians were the clearest case of restricted cranial diversification. They exhibited the lowest skull shape disparity among habitats (Fig. 4C; table S8), and unlike marine eupercarians, freshwater species (including members of Percidae, Centrarchidae, Sciaenidae, and Terapontidae)^30,31,57,58^ did not extend beyond the Percomorph Pile in morphospace (Fig. 2). This pattern was not explained by sampling (figs. S12–S14). When disparity was standardized by the evolutionary time lineages spent in each habitat, freshwater lineages generated moderate disparity relative to opportunity (Fig. 4G), exceeding coastal and deep-demersal clades on a per-time basis. However, when standardized by the number of in-situ speciation events, freshwaters remained the least disparate habitat (Fig. 4I). Cranial diversification has therefore not kept pace with species diversification in freshwater environments.

Ancestral habitat reconstructions further suggest that restricted freshwater morphospace occupation may reflect limited colonization pathways rather than evolutionary stasis^62^. Brackish environments received colonists from both freshwater and shallow marine clades, whereas freshwater habitats received colonists almost exclusively from brackish lineages (figs. S16–S19), consistent with brackish habitats acting as the primary gateway into fresh waters. For all metrics considered (evolutionary rate, total disparity, disparity per unit time, and disparity per *in-situ* speciation; Fig. 4), freshwater eupercarians ranked at or near the bottom, supporting a pattern of constrained morphospace occupation.

Deep-sea habitats showed the opposite tendency. Deep-demersal clades had the highest rates of cranial evolution (Fig. 4E), whereas deep-pelagic lineages showed the highest disparity per unit time (Fig. 4G). Deep-pelagic species also showed no overlap with the Percomorph Pile and were characterized by large, wide jaws, elongate teeth, and reduced orbits. These results suggest that deep-demersal and deep-pelagic settings promoted rapid cranial change through different components of the evolutionary process. By contrast, coastal demersal clades had the greatest total disparity (Fig. 4C) but much lower standardized values, consistent with slower or plateauing accumulation of skull variation. Brackish lineages showed low evolutionary rates and only moderate raw and standardized disparity (Fig. 4C, E, G, I), reflecting their role as a transitional habitat receiving colonists from multiple parts of the tree rather than a setting of unusually rapid in-situ cranial diversification.

### Diet structures the functional pathways of skull diversification

Diet was also strongly associated with skull shape across all 46 phylogenies (R² = 0.040–0.236, all *p* < 0.005) and evolutionary rate, but its signal was more directly tied to functional cranial variation than the habitat signal. Models combining both predictors (habitat and diet) explained more cranial shape variance than either predictor alone on every tree (combined R² = 0.085–0.128; table S7; the wider single-predictor ranges, habitat 0.022–0.190 and diet 0.040–0.236, reflect variation across the 46 trees rather than a larger effect on any one tree), indicating that cranial diversification reflects both the ecological setting in which species live and the trophic functions their skulls perform. Ancestral-state reconstructions recovered benthivore/invertivore as the most probable root diet in the four all-genes trees (mean posterior = 0.506), followed by durophagy (0.231), piscivory (0.142), herbivory/detritivory (0.109), and planktivory (0.012; Fig. 4B). Across diet guilds, herbivore/detritivores and planktivores showed the fastest rates of skull shape evolution, whereas piscivores and durophages evolved more slowly (Fig. 4F; table S8). These rate differences indicate that trophic ecology affected not only where lineages fall in morphospace, but also the tempo of cranial change.

Diet guilds also occupied distinct, although partly overlapping, regions of morphospace. Benthic invertivores were broadly distributed and were primarily characterized by sub-terminal mouths. Durophages were also highly disparate, with short powerful jaws, short teeth, and slightly expanded orbits. Herbivore/detritivores were more restricted and were characterized by elongate skulls, short jaws, and posteriorly displaced orbits. Piscivores clustered mostly near the center of morphospace, close to the Percomorph Pile, and were characterized by wider neurocrania, large wide jaws, and enlarged orbits. Planktivores were widely distributed but generally had truncated skulls, small jaws, and anteriorly displaced orbits.

These trophic patterns differed depending on whether disparity was measured as total occupation of morphospace or standardized by evolutionary opportunity. Durophages showed the greatest total skull disparity, consistent with the varied mechanical demands of processing hard-shelled prey (Fig. 4D). In contrast, planktivorous lineages stood out when disparity was standardized by time, indicating rapid cranial exploration relative to the amount of evolutionary time spent in this trophic niche (Fig. 4H). Collectively, these results suggest that habitat defines the broad ecological arena in which cranial diversification occurs, whereas diet more directly tracks the functional routes by which skull shape changes.

### Ecology shapes convergence toward ancestral and novel skull shapes

We applied complementary convergence tests across the four all-genes trees and classified results into six categories reflecting convergence on ancestral or derived skull shapes (Fig. 2; Table 1). Among habitats, coastal demersal lineages had a significant rate-based signal but no root proximity, consistent with retention of generalized skull forms. Freshwater lineages provided the clearest evidence for root-proximal convergence, with significant RRphylo and Wheatsheaf signals, the lowest disparity, and the shortest root distance (Table 1). Shallow-pelagic and deep-pelagic lineages had weak or no convergence signal. Deep-demersal lineages had significant rate-based and Wheatsheaf convergence, strong phylogenetic signal, and root distance differing from the null (*RRphylo* < 0.001; Wheatsheaf = 1.170, *p* ≤ 0.001; Blomberg’s *K* = 1.38; root distance = 22.809, *p* = 0.003). Brackish lineages produced only a rate-based signal (*RRphylo* = 0.019), consistent with a mixed assemblage.

**Table 1.** Multi-metric evaluation of phenotypic convergence across ecological groups. We assessed convergence using three complementary approaches (*RRphylo*, *Ct*1, and *Wheatsheaf index*) and contextualized these results with metrics describing phylogenetic signal (*Blomberg’s K*), trait disparity (*CV*), and divergence from the reconstructed ancestral state (*Root distance* and *Root p*).

| *Ecological group* | *Convergence metrics* |  |  | *Phylogenetic signal and skull disparity* |  | *Root reconstruction* |  |
| --- | --- | --- | --- | --- | --- | --- | --- |
|  | *RRphylo* | *Ct* | *Wheatsheaf index* | *Blomberg's K* (higher = stronger signal) | *CV* (lower = + uniform) | *Root distance* (lower = nearer root) | *Root p* |
| ***Habitat*** |  |  |  |  |  |  |  |
| Coastal demersal | **0.001\*\*\*** | — | 0.92 | 1.07 | 0.44 | 30.5 | 1.000 |
| Shallow pelagic | 0.026* | 0.145 | 0.86 | 1.00 | 0.71 | 28.9 | 0.633 |
| Deep pelagic | 0.249 | — | **1.2\*\*\*** | 1.09 | — | 30.9 | 0.676 |
| Deep demersal | **0.001\*\*\*** | 0.185 | **1.2\*\*\*** | **1.38** | 0.54 | 22.8 | **0.003\*\*** |
| Brackish | 0.019* | 0.335 | 1.0 | 0.38 | 0.54 | 24.9 | 0.117 |
| Freshwater | **0.001\*\*\*** | 0.485 | **1.7\*\*\*** | 0.41 | **0.36** | **19.7** | **< 0.001\*\*\*** |
| ***Diet*** |  |  |  |  |  |  |  |
| Piscivore | **0.001\*\*\*** | **0.010\*\*** | **1.3\*\*\*** | 0.74 | 0.51 | **22.1** | **< 0.001\*\*\*** |
| Benthivore / invertivore | **0.001\*\*\*** | **0.010\*\*** | 1.0 | 0.98 | 0.56 | 25.7 | **0.009\*\*** |
| Durophage | 0.655 | 0.510 | 0.93 | **1.54** | 0.65 | 30.4 | 0.956 |
| Planktivore | 0.070 | 0.235 | 0.98 | — | 0.65 | 32.5 | 0.804 |
| Herbivore / detritivore | **0.001\*\*\*** | 0.060 | 0.95 | 0.82 | **0.29** | 39.5 | 1.000 |

*θ* (*RRphylo*), *Ct1*, and *Wheatsheaf index* are complementary metrics of phenotypic convergence. *Blomberg's K* measures phylogenetic signal, *CV* measures within-group trait disparity, *Root distance* measures divergence from the reconstructed ancestral state, and *Root p* tests whether that divergence differs from null expectation. For other *Ct* metrics see Tables S10 (habitat) and S11 (diet). Bold values with asterisks indicate significance (\**p* ≤ 0.05; \*\**p* ≤ 0.01; \*\*\**p* ≤ 0.001). Dashes indicate unavailable values.

Among diet guilds, piscivores provided the strongest evidence for root-proximal convergence, with all three metrics significant and the shortest root distance (*RRphylo* < 0.001; Ct1 *p* = 0.010; Wheatsheaf = 1.324, *p* ≤ 0.001; root distance = 22.133, *p* < 0.001). Benthivore/invertivores also had significant *RRphylo*, Ct1, and root proximity (*RRphylo* < 0.001; Ct1 *p* = 0.010; root *p* = 0.009), although the Wheatsheaf index was not significant. Durophages had no significant convergence despite the highest Blomberg’s *K* (1.54), suggesting phylogenetic conservatism rather than repeated independent convergence. Planktivores likewise lacked clear convergence, whereas herbivore/detritivores displayed a distinctive derived pattern with significant *RRphylo*, low disparity, and the greatest root distance among trophic guilds (*RRphylo* < 0.001; CV = 0.292; root distance = 39.470), although Ct1 and the Wheatsheaf index were not significant).

Skull shape convergence therefore took different forms depending on ecological context: freshwater lineages converged on the ancestral percomorph skull shape, coastal demersal lineages retained shapes near the ancestral condition, and guilds with specialized feeding demands, such as herbivore/detritivores, converged on novel, derived morphologies. Shared ecology drove phylogenetically distant lineages toward similar cranial forms, but the direction of convergence depended on the functional demands imposed on the skull. Because Ct1, Wheatsheaf, and *RRphylo* each capture different dimensions of convergence, non-significant results for individual metrics should not be taken as evidence against convergence; our framework instead evaluates consistency across complementary tests alongside phylogenetic signal and root proximity (Table 1; table S10).

### Elevated skull evolutionary rates accompanied a two-phase expansion of eupercarian morphospace

BayesTraits^63,64^ analyses run across the four all-genes trees suggested that the rate of eupercarian skull shape diversification was highly heterogeneous across the history of this radiation. The best-fitting model was a variable-rates Brownian motion process with a lambda tree transformation (fig. S20). High evolutionary rates characterized the Tetraodontiformes (pufferfishes), Lophiiformes (anglerfishes), Acanthuriformes (surgeonfishes), and Chaetodontiformes (butterflyfishes), all within the superorder Lophiimorphae (Fig. 5; figs. S21–S22). Members of these groups make up the extremes of the eupercarian morphospace (Fig. 2; fig. S10), suggesting that high evolutionary rates are associated with breaking away from the Percomorph Pile. Exceptionally high rates on the deep branches ancestral to these orders (Fig. 5) suggest that rate shifts occurred early and were retained across taxonomic orders.

Disparity-through-time analyses^71^ revealed that skull shape diversification unfolded over two temporal phases. Specifically, skull shape disparity was partitioned among subclades relative to the null expectation during the initial phase of diversification (Fig. 5; fig. S23). Ancestral habitat and diet reconstructions suggest that early eupercarian diversification took place mainly in coastal demersal or brackish settings (Fig. 4; figs. S12–S14) and was dominated by benthivorous/invertivorous and piscivorous trophic regimes, implying a connection between early skull shape diversification and interaction with the benthos in shallow waters^72^. Following the K–Pg mass extinction (66 Ma), a second phase unfolded throughout the Cenozoic, a broad period characterized by global climate fluctuations^44,73^ and rampant habitat transitions in Eupercaria (Fig. 4). During this latter phase, disparity was partitioned within subclades (Fig. 5; fig. S23), and diversification involved repeated shifts into more specialized guilds such as planktivory, durophagy, and herbivory/detritivory. This bi-phasic pattern, which was recovered consistently among the four all-genes tree variants (fig. S23), is consistent with an adaptive radiation giving early origins to the major clades within Eupercaria^3,17,71,74^, followed by steady diversification and evolution of novel phenotypes.

### Ecological opportunity and trophic specialization let some lineages escape the Percomorph Pile

Differences in ecological opportunity among habitats and trophic guilds likely explain why some lineages diverged from the Percomorph Pile while others did not. Freshwater habitats are renowned for adaptive radiations in mostly non-eupercarian groups such as cichlids and whitefishes (Cichlidae, Salmonidae)^70,75–79^. Freshwater eupercarians also show ecological and morphological divergence^30,31^, and unmeasured traits may reveal greater ecomorphological diversity. Yet, at least for skulls, freshwater eupercarians lack the extreme novelty seen in marine eupercarians, such as wolftrap anglerfishes^80^, porcupine fishes^25^, and surgeonfishes^81^, even though coastal demersal clades show the lowest relative disparity per unit time (Fig. 4G).

Several factors may explain this low disparity. Otophysi, the dominant freshwater fish clade, is much older than Eupercaria^29,84,85^ and contains high species and morphological diversity in tropical rivers^82,86^. Because freshwater is derived in Eupercaria (Fig. 4), otophysan incumbency likely limited ecological opportunity for eupercarian colonizers. Freshwater eupercarians are concentrated in North America and Australasia^30,31,57^ rather than species-rich tropical basins such as the Amazon and Congo, consistent with ecological saturation in those systems^76^. Where freshwater eupercarians diversified, they often occupied hyper-piscivorous niches, a dietary strategy rare in Otophysi but one that can constrain skull shape evolution. Their low disparity therefore likely reflects both limited ecological opportunity and functional constraints^16,87^ imposed by the niches available to late-arriving colonizers.

In marine habitats, Lophiimorphae stands out as the most diverse and rapidly evolving eupercarian lineage (Figs. 2, 5; figs. S9–S10). Although many lophiimorphs are reef-associated, the clade also colonized the deep sea (spikefishes and anglerfishes) and open ocean (louvar and mola). Reefs likely provided ecological opportunities for distinctive adaptations^8,12,81,88^, but repeated transitions into other marine habitats also contributed to morphological diversification^14,25^. Among the nine series of percomorphs^20^, Eupercaria is unusual in spanning marine habitats from polar regions to the bathypelagic zone. By contrast, Ovalentaria, the second-most diverse series, includes speciose groups such as cichlids and blennies but is largely restricted to shallow marine and freshwater habitats. This contrast suggests that repeated habitat transitions exposed eupercarians to novel ecological opportunities, with some lineages having greater evolutionary capacity to respond.

Trophic specialization shaped escape from the Percomorph Pile in ways that complemented habitat. Durophages occupied the broadest region of cranial morphospace and converged on a derived form with short, powerful jaws and reinforced tooth plates, consistent with the demands of crushing hard-shelled prey (Figs. 3, 4D). Herbivore/detritivores showed a different derived pattern, with elongate skulls, short jaws, posteriorly displaced orbits, and the greatest mean root distance among trophic guilds (Table 1). Planktivores were disparity-rich relative to opportunity (Fig. 4H), consistent with rapid exploration of skull modifications used to capture small prey in the water column. Piscivores and benthivore/invertivores, by contrast, retained skull shapes closer to the ancestral percomorph configuration. Thus, diet-driven escape from the Percomorph Pile was concentrated in guilds with strong functional demands on the feeding apparatus, whereas simpler ram-and-suction feeding allowed many lineages to retain the generalized perch-like skull.

**Fig. 3.**
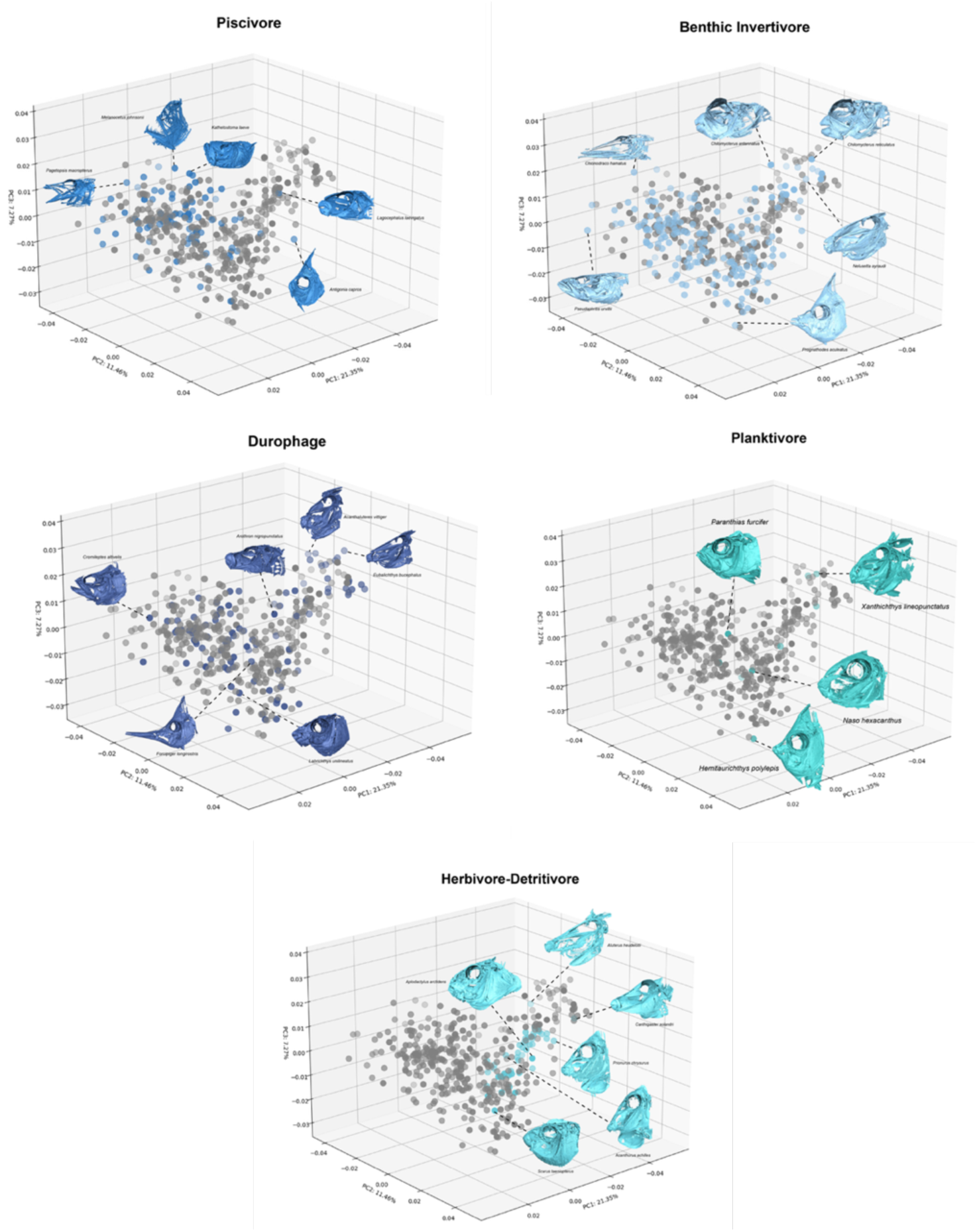
Cranial morphospace partitioned by diet. Three-dimensional morphospace plot for eupercarian species along three principal component axes of skull shape variation. Species belonging to each trophic guild are highlighted in color within the morphospace; all remaining species are shown in gray for reference. Insets show representative CT-reconstructed skulls from different regions of morphospace. Durophages occupied the widest distribution of cranial morphospace among trophic categories; piscivores and herbivores/detritivores were more restricted. Diet explained 4.5% of cranial shape variance (phylogenetic Procrustes ANOVA; *p* = 0.005).

We estimate that the ancestral eupercarian had a generalized perch-like skull associated primarily with benthivorous/invertivorous feeding, a combination that was strongly modified in some clades but retained with only minor changes in others (Figs. 2, 5). This skull shape likely evolved early within spiny-rayed fishes (acanthomorphs), as suggested by similar perch-like skulls in non-eupercarian clades such as cichlids, snooks (Centropomidae), and Nile perches (Latidae). Many of these groups were still classified in Perciformes as recently as the 2010s (Table S3). Our findings echo Gregory’s 1933 view^89^ that many teleost lineages retained a “primitive percomorph skull type,” whereas others “lost the primitive percomorph type” and assumed novel forms. Because perch-like species occur in every aquatic habitat (Fig. 2), and because benthivory/invertivory is a widespread trophic strategy, the persistence of this generalized skull shape suggests that it is highly versatile and functional across ecological settings.

## Conclusions

Differences in evolutionary rates among clades, combined with the ecological opportunities offered by different aquatic environments and selective pressures associated with different trophic ecologies, shaped skull diversification in one of the largest vertebrate radiations. Ecology sets the stage for how phenotypic diversification plays out, from restricting it to a handful of forms to enabling morphological innovation. Three major findings emerge from this work. First, our densely sampled phylogenomic framework provides the most robust backbone yet assembled for Eupercaria, with stronger resolution of higher-level relationships and a foundation for a revised, more stable classification (Fig. 1, fig. S5, Data S1). Divergence-time estimates were consistent regardless of whether the Late Cretaceous fossil †Plectocretacicoidea was used as a calibration (Fig. 1C), confirming that major eupercarian lineages were established by the Late Cretaceous.

Second, we present one of the largest three-dimensional cranial datasets for vertebrates yet assembled from micro-CT scans of museum specimens. This dataset complements recent work on fish body shape^45^ and enables comparisons between cranial and axial skeletal evolution^41,56^. Habitat and trophic ecology made independent, complementary contributions to skull diversification, but most variation remained organized around a conserved generalized perch-like architecture, the Percomorph Pile. Habitat captured the ecological setting, from freshwaters to the deep sea, whereas diet reflected functional pathways shaping skull form. Freshwater lineages remained closest to the ancestral condition and ranked near the bottom across disparity metrics despite high species richness, consistent with late arrival into systems occupied by older otophysan radiations and with constraints of the hyper-piscivorous niches available to eupercarian colonizers. Deep-sea lineages showed rapid skull evolution, with the highest rates in deep-demersal clades and the greatest disparity per unit time in deep-pelagic lineages. These habitats are rarely compared under the same framework, with past studies focusing on shallow marine versus deep-sea habitats^14,44,90^, or freshwater versus marine habitats^4,85,91,92^. Among trophic guilds, durophages showed the greatest total skull disparity, whereas planktivores showed high disparity relative to opportunity. Habitat and diet are therefore not redundant axes but explain complementary aspects of cranial diversification.

Third, ecological context shaped both the direction of convergence and the evolution of novelty. Freshwater lineages repeatedly converged on skull shapes close to the ancestral percomorph condition, whereas specialized trophic guilds converged on more derived forms. The most extreme departures from the Percomorph Pile were concentrated in Lophiimorphae, where high evolutionary rates on deep internal branches indicate that rapid skull shape evolution was established early and retained across the clade. Thus, ecological opportunity alone did not generate morphological novelty; lophiimorphs also had an unusual capacity to evolve away from the ancestral skull plan. More broadly, the timeline of eupercarian skull diversification followed a two-phase trajectory, with disparity first partitioned among subclades during the Late Cretaceous and later accumulating within subclades through the Cenozoic. In all, our results show that ecological setting, evolutionary opportunity, and colonization history interacted to determine not only how much skull shape diversity accumulated but also whether lineages were retained near the widespread ancestral perch-like morphology or evolved toward more specialized and derived cranial designs.

## Materials and methods

### Taxonomic sampling

Taxonomic sampling began with 1,135 ingroup and 37 outgroup individuals representing 1,131 species (Table S2). New genomic data was collected from 1,108 individuals as part of the NSF-funded FishLife project. Of these, 355 were recently published in studies on targeted subclades^14,26,33,54,96^, and 753 are newly published in this study. The remaining 64 individuals represent previously published exon capture data or exons mined from published genomes available on NCBI^48^.

DNA was extracted using the DNeasy Blood and Tissue Kit (Qiagen, Valencia, CA). We shipped DNA extractions to Arbor Biosciences (Ann Arbor, MI) for library preparation, target enrichment, and sequencing. Sequencing of paired-end 150-bp reads was completed on a HiSeq 4000 with a total of 192 samples multiplexed per lane. Target capture probes were based on a set of 1,105 single-copy nuclear exon markers designed for fish phylogenomics^42^ (Eupercaria bait set) plus 19 legacy markers^20^ and mitochondrial DNA.

After quality control steps (see below), final sampling of eupercarians included 1,083 individuals representing 1,050 species, 484 genera and 133 families. Twenty-eight families were included as outgroups, spanning the acanthomorph lineages Holocentriformes, Ophidiaria, Batrachoidiaria, Pelagiaria, Syngnatharia, Gobiaria, Anabantaria, Carangaria, and Ovalentaria^20^. To assess the relative degree of taxonomic sampling, we compared our sampling against the species list compiled by Rabosky et al. ^35^ and available at www.fishtreeoflife.org. We chose this reference because the family-level taxonomic scheme was more comparable to our list (as it is based on ref. ^20^) compared to FishBase^97^ or the Catalog of Fishes^98^ as of June 2024. Based on this reference list, our phylogenetic sampling comprises 15.1% of species, 39.4% of genera, and 82.6% of families in Eupercaria (Table S1).

Due to the cost of sequencing and computational demands, systematic studies typically prioritize sampling many taxa or many genes, but rarely both. Nonetheless, the ultimate goal of systematics is to build a complete Tree of Life sampling all taxa under a robust, comprehensive molecular framework. To assess progress towards this goal for Eupercaria, we also compared the sampling of eupercarians in the trees of Rabosky et al.^35^ (species with molecular data only) and Ghezelayagh et al.^17^ against the same reference species list (Table S1). Our study has three times higher species-level sampling and ∼60% higher genus-level sampling than Ghezelayagh et al., the best-sampled phylogenomic-scale study of spiny-rayed fishes at the time of writing.

### Assembly and quality control

Assembly, initial raw data quality control steps, and alignment were conducted using the pipeline^42^ available at https://github.com/lilychughes/FishLifeExonCapture. Low quality raw reads and adapter contamination were trimmed using Trimmomatic v.0.39^99^. Trimmed reads were mapped against the reference sequences used for probe design with BWA v.0.7.17^100^ and PCR duplicates were removed using SAMtools v.1.9^101^. An initial sequence for each marker was assembled with Velvet v.1.2.10^102^, and the longest contig was used as a reference sequence to extend contigs using aTRAM 2.2^103^ with the Trinity v.2.2 as the assembler^104^. Redundant contigs were excluded with CD-HIT-EST v.4.8.1^105,106^, and open reading frames for the remaining contigs were identified using Exonerate v.2.4.0^107^. Redundant contigs with reading frames exceeding 1% sequence divergence were discarded.

Nuclear loci that were assembled successfully (n=1,105) were aligned using MACSE v.2.03^108^ with the “trimNonHomologousFragments” option. Some legacy markers can retain paralogues when obtained using this target capture probe set and deserve additional scrutiny^42^. For these markers, we checked their gene trees by eye for pseudogenes. Ten markers had pseudogenes and were excluded from our dataset. After these steps, the final dataset contained 1,095 loci (1,086 FishLife exons and 9 nuclear legacy markers).

Further quality control steps are based on the pipeline described by Arcila et al.^96^. First, we visually inspected an initial phylogeny made with FastTree v.2.1.11^109^ and removed individuals that were obviously contaminated (i.e., found in the wrong order or series). We referenced CO1 sequences captured using the FishLife probe set^42^ against the Barcode of Life Data System (BOLD) database^110^ using scripts from the “fishlifeqc” package available at: https://github.com/Ulises-Rosas/fishlifeqc. BOLD results are reported in Table S2. Individuals that did not match to at least to genus level were re-inspected in the initial tree. Some taxa belonged to groups with poor sampling or with necessary systematic revision, where greater sampling is needed to be confident of a misidentification (noted in Table S2). Otherwise, mismatched individuals were interpreted as misidentified and dropped from the dataset. Next, for those species with multiple individuals sampled, we visually inspected the preliminary tree to confirm that species were monophyletic and corroborated these relationships against the BOLD results. Individuals that were more closely related to other species were removed from the dataset for being potentially misidentified. In the initial tree, we also inspected individuals with very short branch lengths between them (“T-clades”). If sister taxa belonged to different species or genera yet had very short branches, these were interpreted to be misidentified and were removed from the dataset. An additional six taxa were removed for having very few exons assembled to avoid issues related to missing data. After these quality control steps, 52 taxa in total were removed from the dataset (Table S2).

Using the final set of taxa, we performed branch length correlation (BLC) tests^111^ to detect additional within-gene contamination that may not be easily detectable once genes are concatenated. The logic of this test is that contaminated sequences will show very long branches once constrained to a reference topology. We generated a reference phylogeny using the program IQ-TREE v2.1.2 “COVID edition”^112^ based on the concatenated alignment of 1,095 genes and using mixture models^113^. We then generated gene trees for each marker with topology constrained to match the reference phylogeny. We generated a branch-length ratio for every taxon in every gene tree, which was the length of the branch in the gene tree over the length of the corresponding branch in the reference tree (after pruning the reference tree to the same individuals contained by the gene tree). Branches with a ratio >5 were flagged (totaling 1.37% of sequences in the dataset) and discarded due to potential contamination. After removing these sequences, genes were un-aligned using the “unalign.md” script within the Goalign toolkit^114^, then re-aligned following the steps above. The final alignment used for phylogenetic inference was 485,136 base pairs long, and alignments for individual markers varied in length from 108–2,718 bp (mean 443 bp).

### Phylogenetic inference and classification

After quality control, the 1,095 markers were concatenated using utility scripts in the AMAS package^115^. A concatenation-based phylogeny was evaluated with maximum likelihood using the program IQ-TREE v2.1.2 “COVID edition”^112^ implementing mixture models^7^ (option -m set to “MIX{JC,K2-,HKY,GTR}”). Support was measured using 1,000 Ultrafast bootstrap replicates ^116^ with the “-bnni” option to reduce the risk of overestimating support due to severe model violations. To account for potential incomplete lineage sorting, we also performed a multi-species coalescent analysis using ASTRAL-II v.5.7.1^117^ based on gene trees estimated using IQ-TREE with the same settings as above. Prior to use with ASTRAL, nodes within gene trees with bootstrap values <33% were collapsed into polytomies to reduce noise^118^. Support was evaluated using local posterior probabilities^119^ (option “-t 3”).

To account for uncertainty in topology and divergence times in downstream analyses, we evaluated additional trees based on subsets of our molecular dataset^33,120^. We divided the dataset into 11 subsets with 105 genes each. Six of these genes (“anchor genes”) contained all taxa when concatenated together, and were common to all subsets to ensure all taxa were sampled in all subsequent trees. The remaining genes were semi-randomly assigned to a subset (99 genes per subset) using a custom R script that minimized the standard deviation of the number of parsimony-informative sites among subsets. For each subset, we inferred concatenation-based and coalescent-based species tree using IQ-TREE and ASTRAL, respectively, using the settings listed above. This gene subset approach differs from the more common practice of using trees from a Bayesian posterior distribution estimated from a single dataset, because it ensures that variation among the trees is due to differences in the underlying genomic data^33,120^.

We visualized disparity in topology and branch lengths with a tree space plot (fig. S1) made using multidimensional scaling (MDS) visualization implemented with the R package phytools v.1.5–1 ^121^. One IQ-TREE gene-subset tree appeared to be an outlier in tree space and so we excluded it from downstream analyses. The final set of 23 trees included two “all-genes” trees (IQ-TREE and ASTRAL), 11 ASTRAL “gene-subset” trees, and 10 IQ-TREE gene subset trees. Figure S2 compares the topologies of the two all-genes trees, fig. S3 compares the topologies among the eleven concatenated trees, and fig. S4 compares the topologies among the twelve coalescent species trees; figs. S5–S7 compare the topology of our all-genes IQ-TREE to the topologies of three relevant, widely referenced phylogenies: Ghezelayagh et al.^17^ (fig. S5), Rabosky et al.^35^ (fig. S6), and Betancur-R et al.^20^ (fig. S7).

We updated the classification scheme from Betancur-R et al.^20^ for Eupercaria in accordance with our results and other recent studies, naming clades that are stable across multiple sources. Our classification scheme has important differences from that recently proposed by refs.^17,46,47^. Most notably, the five major clades of Eupercaria are recognized as superorders, allowing us to retain traditional order-level names such as Tetraodontiformes. Our scheme recognizes 16 orders, 42 suborders, and 163 families across Eupercaria. Details and justification of our new classification are given in Data S1 and table S3.

### Divergence time estimation

We assembled a list of 64 node calibrations from the literature, including 8 outgroup and 41 ingroup fossil calibrations based on well-preserved articulated skeletal remains, as well as geologic calibrations based on the Isthmus of Panama to constrain the divergence time of 15 sister-species pairs (Data S2, table S5). Following the recommendations by Parham et al.^122^, we determine hard lower bounds for each calibration as the minimum age of the oldest fossil assignable to a given clade. We then estimated soft upper bounds of calibrations based on the algorithm proposed by Hedman^123^. This approach uses a list of fossil outgroup age records based on the oldest minima to produce an age distribution for the time of origin of a clade. We extracted the 95% credible interval for each calibration point.

We used two calibration schemes including or excluding the controversial fossil †*Plectocretacicus clarae*, which we placed on the most recent common ancestor (MRCA) of Tetraodontiformes and Lophiiformes^26^. The extinct superfamily †Plectocretacicoidea is purportedly a stem tetraodontiform, and phylogenetic analyses using morphological characters place it as the sister to all remaining Tetraodontiformes^49,50,124^. The earliest plectocretacicoid fossils are 94 Ma^124^. Therefore, due to the apical position of Tetraodontiformes within Eupercaria, this fossil has potential to greatly increase the age of early nodes in the phylogeny. However, some authors do not believe †Plectocretacicoidea are related to Tetraodontiformes, or at least that the evidence for such a relationship is uncompelling^21,125–127^, so we also used a calibration scheme without this fossil.

To generate time-calibrated phylogenies, we first selected a subset of 22 genes (13,335 bp) chosen to retain the same proportion of parsimony-invariable sites as in the full dataset with low amounts of missing data. The best substitution model for each codon position was estimated in IQ-TREE using ModelFinder^128^. This subset was used as input for divergence time estimation in BEAST v2.7.7^129^. Using the two constrained topologies inferred from the full dataset (IQ-TREE and ASTRAL), we ran four BEAST analyses, each using one of the two topologies and either including (+Plecto) or excluding (−Plecto) †Plectocretacicoidea from the calibration scheme. Each analysis was run twice for 300 million generations under the Optimized Relaxed Clock^130^. Calibration densities were specified as log-normal distributions by setting the standard deviation to 1.0 for all priors and adjusting the mean in BEAUti to match the 95% credible interval estimated for each calibration point. The root of the tree was calibrated using a normal distribution with a mean of 133 Ma, an average based on previous studies (see table S4), and a standard deviation of 5, which encompasses all values reported in the same table within three standard deviations of the mean.

To calibrate the 21 gene-subset trees, we used congruification^131^ on each tree using a randomly and evenly sampled post–burn-in tree from the corresponding BEAST posterior distribution from either +Plecto or -Plecto analyses, resulting in an additional 42 time-calibrated trees. Congruification was based on shared nodes between the full-dataset tree and each subset tree (e.g., ASTRAL subsets with the ASTRAL full tree, concatenation subsets with the concatenation full tree). We then used the shared ages as secondary calibrations to date the subsets in treePL after estimating the priors with the command *prime* and the smoothing parameter^132^. This approach preserved topological consistency while introducing variation in node ages across subsets, thereby propagating posterior uncertainty into downstream comparative analyses.

In total, we produced 46 time-calibrated trees for use with comparative analyses: four based on the all-genes topologies, and 42 based on gene-subset topologies. The variation in topology and divergence time estimates across these trees is derived from three sources: differences in underlying molecular data, tree estimation approach (concatenated versus summary coalescent), and fossil calibration scheme (with or without †Plectocretacicoidea). Computationally intensive analyses (ancestral-state reconstruction, convergence testing, variable-rate modeling, and disparity through time) used the four all-genes trees; PGLS and pairwise rate comparisons were repeated across all 46. For use with comparative methods, branch lengths of all trees were further adjusted by using the “minBranchLength” function in *paleotree* v.3.4.7 ^133^ to enforce a minimum branch length of 0.1 Ma. This was done because comparative analyses can be skewed by a small number of short branches^45^. We also randomly resolved polytomies present in the ASTRAL trees, which are incompatible with some comparative analyses, using the “fix.poly” function in the package RRphylo v.2.8.1 ^134^.

### Phenotypic dataset

Skull shape was measured using 3-D geometric morphometrics collected from micro-computed tomography (micro-CT) scans^38^ of alcohol-preserved museum specimens (Table S6). New scans were collected using a Bruker Skyscan 1273 scanner housed at Rice University. Skulls were segmented from scales and the rest of the body using Amira v.2020.3^135^ and exported as mesh files. Mesh files were digitized with 106 landmarks (34 fixed and 72 semi-sliding) in the software Stratovan Checkpoint^136^ (fig. S8). Landmarks were treated as bilaterally symmetrical and thus only placed on the left side of the skull^137^. The final CT-scan dataset contained 571 species of Eupercaria (*n*=1 scan per species). Of these, 280 were newly scanned and have not previously appeared in any published study. The remaining 291 were previously used in clade-specific studies by the authors^4,14,25,32,80^ or downloaded from the online repositories MorphoSource (https://www.morphosource.org/) or Virtual Natural History Museum (http://vnhm.de/VNHM/index.php) or shared with the authors by collection managers. Of the taxa sampled in our phylogeny, our 571-species CT scan dataset has representation for all families, 91.1% of the genera, and 54.3% of the species sampled in the tree. To ameliorate the influence of preservation artifacts for artificially inflating shape variation, we performed a local superimposition to standardize the position of individual skull elements^138^ before any downstream comparative analyses.

### Phylogenetic comparative analyses

The primary axes of shape variation were visualized using principal component analysis. To estimate the density of shapes, we projected the first three PC axes into a morphospace and estimated 3-D Gaussian kernel density using the *SciPy* package in Python^139^. We inferred the ancestral eupercarian shape, as well as living species closest or furthest from this shape, with the functions ‘mshape’, ‘findMeanSpec’, and ‘gm.prcomp’ from the R package *geomorph* v. 4.0.5 ^94,140^.

The first 38 PC axes, accounting for 85% of the shape variation, were extracted as input for downstream comparative analyses. Based on these shape axes, disparity through time was visualized using the “dtt” function^71^ in *geiger* v.2.0.11 ^141^. The expected temporal trend in disparity under a Brownian motion null model was estimated based on 10,000 simulations, with the four all-genes trees used as alternative inputs (fig. S23).

We assigned species to habitat categories based on previous compilations^44,91,97,142^ (Table S6). The six categories were: coastal demersal, shallow pelagic (includes coastal and offshore habitats), deep-sea demersal, deep-sea pelagic, freshwater, and a combined brackish, estuarine and diadromous state (e.g. species found primarily in brackish waters). Division into shallow versus deep-sea categories was based on where the species is predominately found, not its maximum depth of record^97^. Our use of ‘demersal’ is general and includes fishes found on or near the benthos, as opposed to being predominantly found within the water column (pelagic). These habitats represent a summary of the commonly hypothesized axes of ecomorphological diversification in fishes^9,31,33,44,90,92,143^. While each habitat category contains possible ecological subdivisions, our inclusive binning strategy maximizes phylogenetic replication needed to gain power for comparative methods^61^.

We assigned species to five trophic guilds (piscivore, benthivore/invertivore, durophage, planktivore, and herbivore/detritivore) using two complementary data sources: the comprehensive gut-content dataset of Parravicini et al.^151^ and the FishBase database^97^. The Parravicini dataset provided gut-content–based trophic assignments for reef-associated species, while FishBase contributed curated trophic-guild annotations for the broader pool of teleostean fishes (including pelagic, deep-sea, and freshwater taxa) used to fill gaps and cross-validate guild membership. This dataset compiles visual gut-content data from adult individuals collected at six locations: Marshall Islands^152^, Puerto Rico and the Virgin Islands^153^, Hawaii^154^, Madagascar^155^, Okinawa^156^, and New Caledonia^151^. The full dataset comprises 13,961 fish guts containing ingested material from 615 fish species. We verified all fish species and family names with the rfishbase R package^157^ and cross-checked them against the species sampled in our phylogeny^151^. The original Parravicini dataset categorized 1,200 individual prey items into 38 ecologically informative groups defined by higher-level prey taxonomy, with the exception of benthic autotrophs (e.g. algae and seagrass), detritus, inorganic material (e.g. sand), and zooplankton (gelatinous and non-gelatinous zooplankton, plus eggs and larvae across all taxa)^151^. We retained only species with non-empty guts from at least three individuals. From these prey-group profiles we assigned each species to one of five focal trophic guilds for downstream analyses (piscivore, benthivore/invertivore, durophage, planktivore, and herbivore/detritivore); the intersection of curated trophic-guild assignments with the 571-species CT-scan dataset comprised 408 species; after excluding two undersampled omnivores, 406 species were retained for the principal diet comparisons.

We estimated ancestral habitats using the “make.simmap” function in *phytools* v.2.2.3 ^121,144^ based on a continuous-time reversible Markov model. We designed an ordered model that prevented biologically unrealistic habitat transitions (table S9). This model enforced two rules. First, transitions between freshwater/euryhaline and the deep sea must include the shallow ocean as an intermediate step. Second, simultaneous transitions between water column use and depth categories were banned (e.g. shallow pelagic to deep-sea demersal); these were treated as separate events. All other transitions were allowed to vary and take on different rate values. SIMMAP analyses were repeated using the four all-genes trees to account for variation in topology and divergence times (figs. S12–S14). We converted the count of each transition type (output by SIMMAP) into a transition rate by dividing it by the total evolutionary time spent in each source habitat (also output by SIMMAP) following ref ^91^ (figs. S16–S19).

To test for a general effect of habitat on skull morphology, we fit a PGLS model relating habitat to landmark-based shape data using the “procD.pgls” function in *geomorph* v. 4.0.5 ^94,140^, assessing significance based on 10,000 iterations (table S7). All 46 trees were used as alternative inputs. Pairwise significance of differences among habitats was assessed using a post-hoc test implemented with the “pairwise” function in the *RRPP* package v.1.3.1 ^146^ (fig. S11). To visualize the shape differences between habitat and trophic groups we used the fitted values of the PGLS analysis to perform partial warps of the mean reference specimen *L. aratus* to the average shape of each diet and habitat group using the *plotRefToTarget* function in geomorph.

Shape disparity, as measured by the Procrustes variance of groups, was statistically compared across habitat categories using the “morphol.disparity” function in *geomorph* (Table S8). Significance was based on a permutation test with 10,000 iterations^145^. We further asked whether low habitat disparity could be explained by convergence among lineages in that habitat. To test this, we used the *convevol* v.2.0.0 package^60,61^ to estimate “Ct” convergence metrics for each habitat. These metrics are based on the commonly-used “C” metrics^60^ but are adjusted to lower the incidence of false positives^61^. We specified the lineages that colonized each habitat independently as the units of comparison within a given habitat (“groups” option in the “convSigCt” function). These lineages were identified based on ancestral state probabilities estimated from SIMMAP (see above). For each terminal taxon in each habitat, we “walked” rootward to identify the node associated with colonization of that habitat (nodes with a probability >70% in the focal habitat and an immediately ancestral node <70%); species that were derived from the same node were assigned the same group. Singleton colonization events, or colonization nodes associated with terminal branches, were excluded because these make up the majority of events and can skew results (figs. S12–S14). Coastal demersal and deep pelagic habitats were excluded from the convevol Ct convergence analysis (the former is likely the ancestral habitat of all eupercarians; the latter had only one lineage with >1 species), although RRphylo, Wheatsheaf, and root-distance metrics were still computed for them (Table 1; Fig. 4). Convergence analyses were run using the first four PC axes only due to computational limits and were repeated using the four all-genes trees as alternative inputs with groups based on the corresponding SIMMAP analysis using that tree. Statistical significance of convergence was assessed from a null distribution based on 1,000 simulations of trait evolution under Brownian motion (table S10).

**Figure 4.**
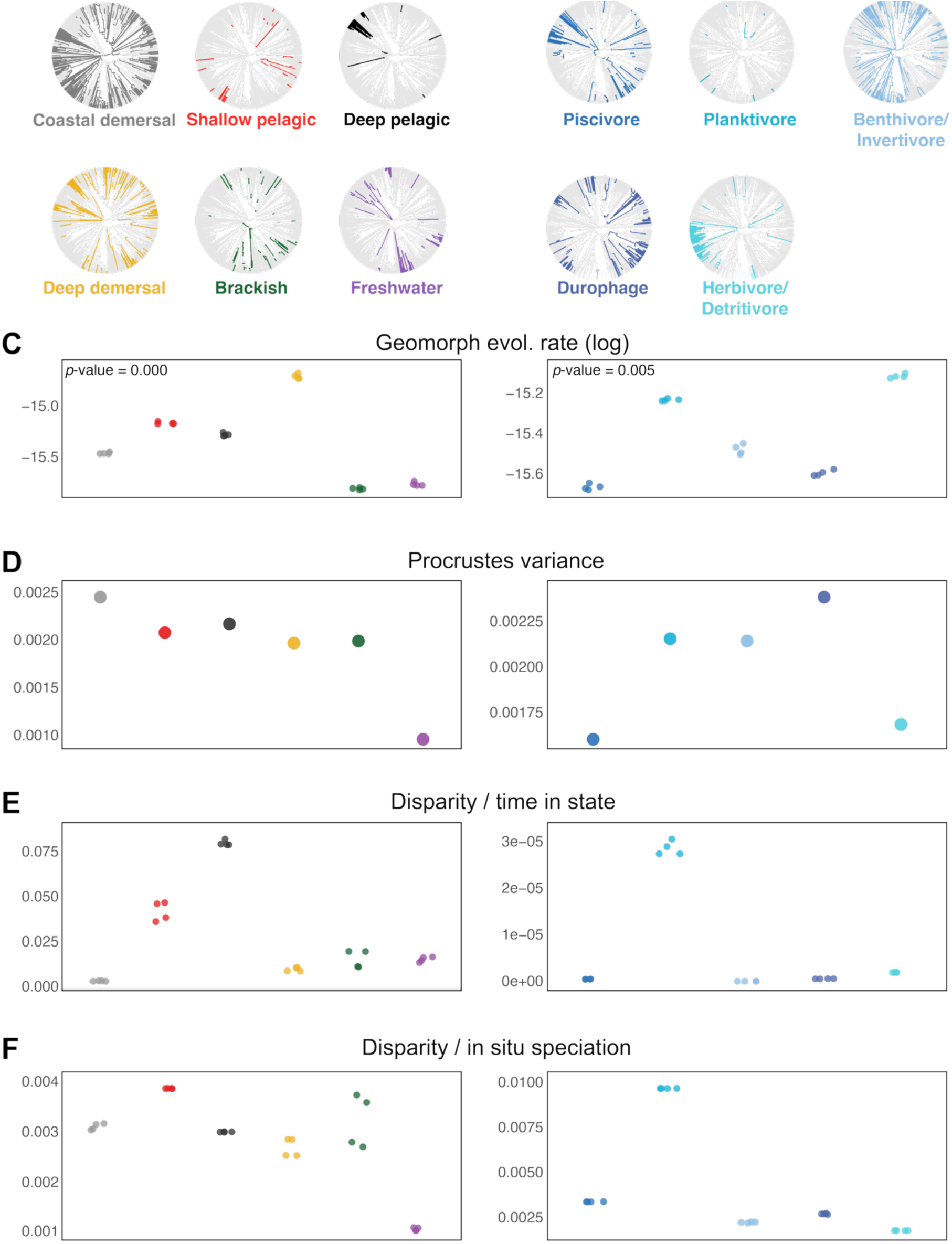
**Routes to accumulating phenotypic disparity across habitats and diet guilds**, shown using the IQ-TREE –Plectocretacicoidea variant (see figs. S12–S14 for the three alternative all-genes trees). (**A, B**) Ancestral reconstructions for habitat (A) and diet (B), with each category displayed separately for clarity. (**C, D**) Total (raw) skull shape disparity by habitat (C) and diet guild (D). (**E, F**) Evolutionary rates of skull shape estimated using *geomorph*^94,95^ by habitat (E) and diet guild (F). Both omnibus tests are significant on each main tree (*p* = 0.005). (**G, H**) Disparity standardized by time spent in each state (evolutionary opportunity) as inferred with SIMMAP, by habitat (G) and diet (H). (**I, J**) Disparity standardized by the number of *in-situ* speciation events, by habitat (I) and diet (J). Multiple dots per category in panels C–J represent values estimated separately for each of the four all-genes trees. Colors correspond to categories shown in panels A and B.

To synthesize the convergence signal across complementary axes, we integrated three convergence tests with measures of phylogenetic signal, within-group disparity, and divergence from the reconstructed ancestral skull shape (Table 1). Convergence was assessed jointly using RRphylo’s *search.conv* (a rate-aware angle-based test, reported as the θ (RRphylo) p-value), the Ct1–Ct4 framework of Grossnickle et al.^61^ (run with *convevol::calcConvCt* and *convSigCt*, with focal lineages defined from stochastic character maps and refined via *pwCheck*), and the Wheatsheaf index (*windex::test.windex*, 1,000 resamples). We contextualized these tests with Blomberg’s K (phylogenetic signal, *phytools::phylosig*), the within-group coefficient of variation of trait values (CV), and Euclidean distance from the species shape to the root state estimated with *phytools::fastAnc* on PC1–PC3, with significance evaluated by a permutation test of 4,999 random draws (root distance and root p). All metrics were computed on the four all-genes time-calibrated trees and combined across trees by taking means for continuous metrics and medians for the Wheatsheaf index. Full implementation details, including software versions, simulation depth, group-merging logic for *convevol*, the BM and OU theta–root calculations, and the multi-tree combining rules used to populate Table 1, are provided in the Supplementary Materials and Methods.

We estimated rates of morphological evolution using two approaches. First, we used the “compare.evol.rates” function in *geomorph* to statistically compare average rates (under Brownian motion) among habitat categories based on 10,000 simulations^95^. We repeated this analysis across all 46 alternative phylogenies (fig. S15). Second, we used BayesTraits V4^63^ to infer branch-specific shifts in evolutionary rates, following recent studies of complex multivariate datasets across vertebrate clades^2,147^. The BayesTraits software allows for fitting user-defined models by specifying transformations of the input tree according to predictions of different evolutionary scenarios^63^, which are then analyzed under the variable rates framework^64^. Since these analyses are computationally intensive, we first determined the best-fitting model of four options based on analyses using the IQ-TREE -Plecto variant as the input tree. These models were a single-rate Brownian motion model and three variable-rate models: Brownian motion, Brownian with a lambda transformation (morphological change related to phylogenetic relationships), and Brownian with a delta transformation (morphological change concentrated near the root of the tree). Bayes factors were used to weigh support for alternative models (fig. S20). We then ran analyses based on the best-fitting model across the four all-genes trees to account for variation in topology and divergence times, repeated three times per tree to determine convergence. Each analysis consisted of a reversible-jump MCMC process run for 400–500 million generations with a burnin of 30%. A stepping-stone sampler was used to estimate the marginal likelihood with 100 stones to run for 1,400,000 generations after convergence. Convergence of the runs was confirmed based on trace plots and Gelman diagnostics near 1 using the R package coda v.0.1.9-4^148^. The run with the best likelihood score of three was used for visualization (Fig. 5, figs. S21–S22). Following software recommendations^63^, we used the BayesTraits setting “TestCorrel” to constrain the correlation between PC axes to zero, because multivariate analyses require independence of traits. The output of variable-rate analyses is a posterior distribution of phylogenies where each branch is scaled by the degree of phenotypic change. BayesTraits output was processed using utility functions from the packages BTProcessR v.0.0.1^149^ and BTRTools 0.0.0.9^150^.

**Fig. 5.**
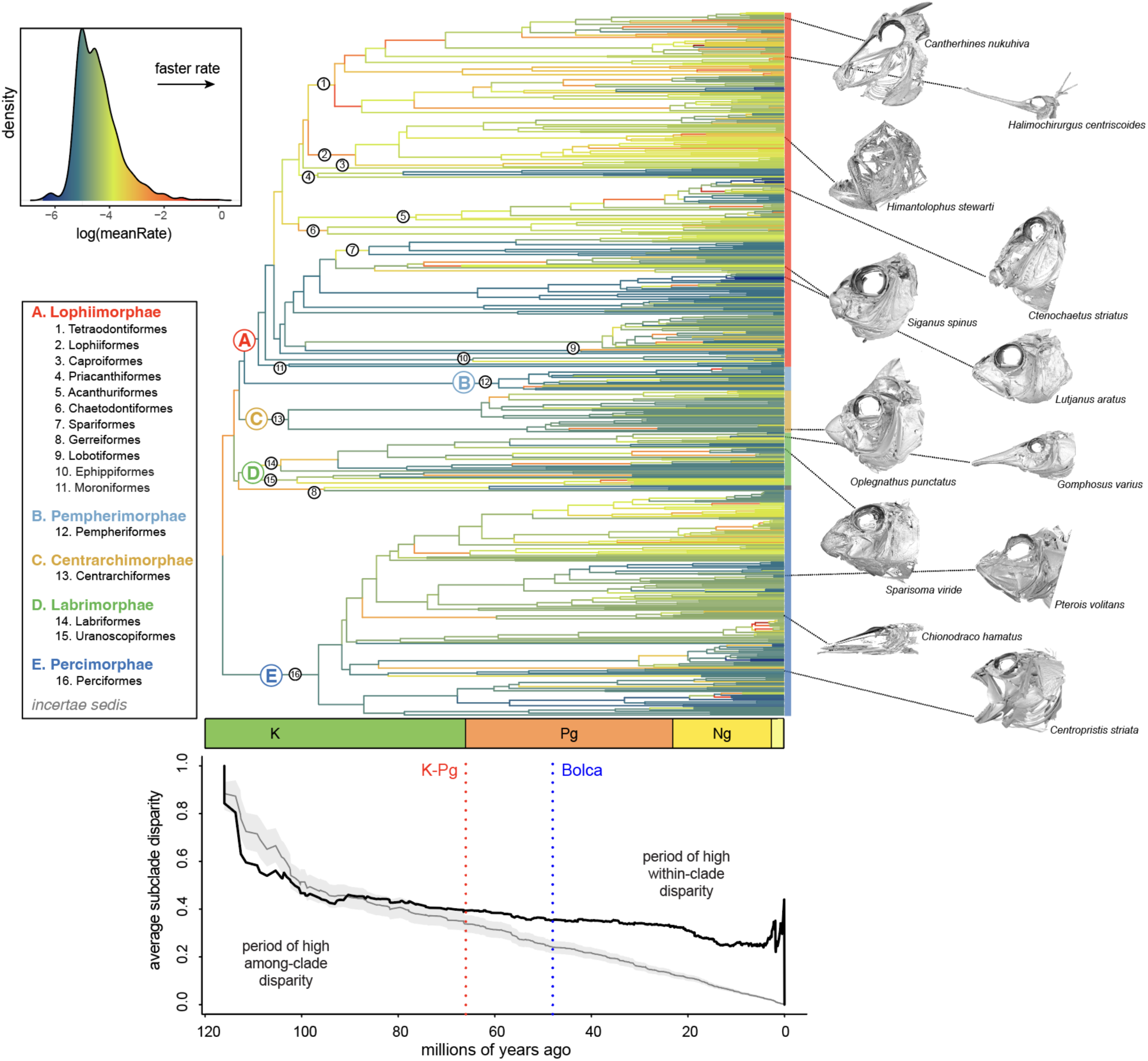
Tempo of skull shape evolution across Eupercaria. Branches of the phylogeny are colored according to the Brownian motion rate of shape evolution inferred by BayesTraits. Results shown here using the IQ-TREE -Plecto variant (see figs. S21–S22 for the three alternative all-genes trees). Taxonomic groups are labeled according to Data S1. Panel below tree shows the bi-phasic evolution of skull shape disparity, with relative disparity partitioned among subclades prior to the Bolca fossil deposits^53^ and within subclades after the Eocene (see fig. S23 for results across the four all-genes trees). The solid black line is the observed disparity, and the dashed black line and grey shading represent the mean and 95% confidence interval as predicted under a null model of Brownian evolution. Bottom right inset shows the distribution of BayesTraits rates associated with terminal branches according to habitats of living species.

## ACKNOWLEDGMENTS

We are grateful to the following museum collections and personnel for providing tissues for DNA extractions and/or voucher specimens for CT scanning: AMS (A. Hay), ANSP (M. Sabaj and M. Arce), ASIZ, AUFT, BGI, BMNH, BSKU, CAS (D. Catania), CBM, CPUM, CSIRO (A. Graham), FMNH (C. McMahan), HUMZ, JCU, JFBM (A. Simons), KU (A. Bentley and L. Smith), LACM (T. Clardy and W. Ludt), LSUMZ (P. Chakrabarty), MCZ (A. Williston and M. Sorce), NCSM (G. Hogue), NMMB, NOAA, NTM, ODU, OS, PKU, QM, RACE Groundfish and Shellfish Assessment Programs of the NOAA Alaska Fisheries Science Center (crews of the F/Vs Sea Storm, Ocean Explorer, Alaska Knight, and Arcturus), ROM (M. Burridge and M. Zur), SIO (B. Frable and P. Hastings), STRI, TCWC (K. Conway), UNMDP, UPR, USNM (C. Huddleston and D. Pitassy), UT, UW (K. Maslenikov and L. Tornabene), VIMS, and YPM (G. Watkins-Colwell and T. Near). Exon capture and sequencing was performed by Arbor Biosciences (Ann Arbor, MI). Computational resources were provided by the University of Oklahoma Supercomputing Center for Education and Research (OSCER) and the George Washington University (Pegasus). We thank Conrad Lee, Grant Parajuli, and Francesca Gastaldo for assistance with CT scanning. We thank members of the FishEvolutionLab and Evans Lab for their feedback on earlier versions of this manuscript. K.E. thanks Dr. Matt Friedman for a passing yet spirited discussion about the ancestral percomorph form in 2018.

## Author contributions

Conceptualization: E.C.S., R.B.R., K.E., D.A. Data curation: E.C.S., R.F., A.S., J.W., J.W.A., C.B., T.J.B., K.C., J.M.D.A., M.R.S., S.M.G., S.P.H., J.K.K., H.L.F., N.L., N.M., M.M., M.P.N., M.M.P., J.J.P., E.M.T., M.W., W.T.W., E.O.W., G.C., G.O., C.M.M., L.C.H., R.B.R., K.E., D.A. Formal analysis: E.C.S., R.F., A.S., J.W., M.R.S., L.C.H., R.B.R., K.E., D.A. Visualization: E.C.S., L.C.H., R.B.R., K.E., D.A. Writing – original draft: E.C.S., L.C.H., R.B.R., K.E., D.A. Writing – review & editing: E.C.S., R.F., A.S., J.W., J.W.A., C.B., T.J.B., K.C., J.M.D.A., M.R.S., S.M.G., S.P.H., J.K.K., H.L.F., N.L., N.M., M.M., M.P.N., M.M.P., J.J.P., E.M.T., M.W., W.T.W., E.O.W., G.C., G.O., C.M.M., L.C.H., R.B.R., K.E., D.A.

## Data and materials availability

All data are available in the main text or the supplementary materials. The phylogenomic datasets are publicly accessible under BioProject PRJNA1378242. All analysis code used in this study has been archived on Zenodo. Tables S2, S3, and S6 are also deposited on Zenodo (DOI: 10.5281/zenodo.17872389).

## SUPPLEMENTARY MATERIAL

Figs. S1 to S23; Tables S1, S4 to S5, S7 to S11; Data S1 and S2.

