## Supplementary Information for "Ecological axes of skull diversification in a massive vertebrate radiation"

**The PDF file includes:**

Materials and Methods
References
Figs. S1 to S23
**Table S1:** Comparing taxonomic sampling across studies.
**Table S2*:** List of genetic material and quality control (Available in Zenodo).
**Table S3*:** Comparing previous classifications of Eupercaria (Available in Zenodo).
**Table S4:** Age estimates for the root (total group Holocentridae) from previous studies (Also available in Data S2).
**Table S5:** List of geologic calibrations (Also available in Data S2).
**Table S6*:** List of voucher material used for CT scanning and habitat codes (Available in Zenodo).
**Table S7:** Results of PGLS comparing skull shape and habitat.
**Table S8:** Pairwise comparison of shape disparity among habitats.
**Table S9:** Ordered habitat transition model for ancestral state reconstructions in SIMMAP.
**Table S10:** Results of evolutionary convergence analyses (using the *convevol* package) using the four "all genes" trees as alternative inputs.

**Other Supplementary Material:**

**Data S1:** Annotated classification of Eupercaria
**Data S2:** Fossil calibrations for Eupercaria

*Given the substantial file size, supplementary material file is archived on Zenodo.

### Materials and Methods

#### *Habitat- and diet-based comparative datasets*

We analyzed ecological correlates of eupercarian skull shape using two partially overlapping datasets. Habitat-based analyses were conducted on the full 571-species skull dataset, for which habitat assignments and morphometric data were available. Diet-based analyses were conducted on a reduced subset of 408 species for which curated trophic-guild assignments were available. Habitat was coded into six categories (coastal demersal, shallow pelagic, deep pelagic, deep demersal, brackish, and freshwater). Diet was initially coded into six trophic guilds, but because only two species were classified as omnivores, that category was excluded from the final diet convergence summaries and visualizations, leaving five focal diet classes and 406 species for the principal diet comparisons. Across all phylogenetic analyses, we accounted for tree uncertainty by repeating each analysis on four alternative time-calibrated phylogenies representing two inference frameworks (ASTRAL and IQ-TREE) and two fossil-calibration schemes (with and without †Plectocretacicoidea). For each tree-specific analysis, we pruned the phylogeny, morphology matrix, and ecological metadata to their shared species set.

#### *Diet assignment from the Parravicini gut-content dataset*

Diet assignments came from two complementary data sources: the comprehensive gut-content dataset of Parravicini et al. (ref. 151 of the main text) and the FishBase database (ref. 97). The Parravicini dataset provided gut-content–based trophic assignments for reef-associated species, while FishBase contributed curated trophic-guild annotations for the broader pool of teleostean fishes (including pelagic, deep-sea, and freshwater taxa) used to fill gaps and cross-validate guild membership. The Parravicini dataset compiles visual gut-content data from adult individuals collated from six locations: Marshall Islands (ref. 152), Puerto Rico and the Virgin Islands (ref. 153), Hawaii (ref. 154), Madagascar (ref. 155), Okinawa (ref. 156) and New Caledonia (ref. 151). It contained 13,961 fish guts holding ingested material from 615 fish species. To ensure that the taxonomic nomenclature was up to date, we verified all fish species and family names with the rfishbase R package (ref. 157) and cross-checked them against the species sampled in our phylogeny (ref. 151). The original Parravicini dataset categorized 1,200 individual prey items into 38 ecologically informative groups defined by higher-level prey taxonomy, except for the following categories that were retained as broader bins: benthic autotrophs (e.g. algae and seagrass), detritus, inorganic material (e.g. sand) and zooplankton (gelatinous and non-gelatinous zooplankton, plus eggs and larvae across all taxa) (ref. 151). We retained only species with non-empty guts from at least three individuals. From these combined prey-group profiles and FishBase annotations we assigned each species to one of five focal trophic guilds for downstream analyses (piscivore, benthivore/invertivore, durophage, planktivore, and herbivore/detritivore); two omnivorous species were excluded from diet-specific convergence summaries because the category was undersampled.

#### *Skull-shape morphospace and ancestral-state reconstruction*

We quantified skull shape from the landmark-based morphometric dataset using principal component analysis of Procrustes-aligned coordinates. For the convergence and ancestral-state analyses, we used the first three principal component axes (PC1–PC3) as a multivariate representation of skull shape. For complementary analyses, including the Wheatsheaf index, we used the first four axes (PC1–PC4). For morphospace visualization, species were plotted in two-dimensional (PC1–PC2) or three-dimensional (PC1–PC3) skull-shape space, with focal ecological groups highlighted against the broader eupercarian background.

To quantify divergence from the reconstructed ancestral condition, we estimated the root state for each tree separately using *phytools::fastAnc* on PC1–PC3 and calculated Euclidean distance between each species and the reconstructed root in multivariate morphospace. For each habitat or diet class, we then calculated the mean distance to the root and evaluated whether that mean was smaller than expected by chance using a permutation test with 4,999 random draws of equal sample size from the same tree-specific dataset. We summarize this statistic in the table as Root distance and Root p. To characterize within-group disparity relative to the ancestral condition, we also calculated the coefficient of variation of species-level root distances within each ecological group. Lower CV values indicate that members of a group are more uniformly clustered around the reconstructed ancestral morphology.

***Phylogenetic signal and trait optima***

We estimated phylogenetic signal within ecological groups using Blomberg's K as implemented in *phytools::phylosig*. We calculated K separately for each of PC1–PC3 within each habitat or diet group when sufficient taxa were available after pruning, using 199 simulations for significance testing. For the summary table, K values were averaged across the available axes and trees. To evaluate whether group-specific trait optima remained close to the ancestral state, we also fit univariate Brownian motion (BM) and Ornstein–Uhlenbeck (OU) models to each PC axis using *geiger::fitContinuous* and extracted the difference between the fitted OU optimum (theta) and the reconstructed root value for that axis. We summarize this as theta–root, where lower absolute values indicate that the estimated optimum remains closer to the ancestral skull configuration.

***Three complementary tests of phenotypic convergence***

We assessed phenotypic convergence using three complementary approaches. First, we used the rate-aware RRphylo procedure *search.conv()*, which tests whether taxa sharing a focal ecological state show greater directional similarity in phenotypic evolution than expected given their phylogenetic history and inferred evolutionary rates. RRphylo models were fit separately for each tree using PC1–PC3, and each habitat or diet state was tested independently with 1,000 simulations. The table reports the mean state-specific *search.conv()* p-value across the four trees. In the manuscript table this column is labeled theta (RRphylo), but it corresponds to the RRphylo angle-based convergence test summarized as a p-value rather than to an OU optimum parameter.
Second, we used the Ct framework implemented in *convevol* to test for convergence among independently derived lineages occupying the same ecological state. Candidate independent focal lineages were identified from stochastic character maps when available. Each focal taxon was traced from the tip toward the root, and taxa sharing the same highest-supported origin node for the focal state were assigned to the same lineage group using a posterior cutoff of 0.70. When stochastic maps were unavailable, we defined provisional lineage groups from focal clades and singleton taxa. We then used *pwCheck()* to detect zero-information inter-group comparisons and iteratively merged problematic groups until invalid comparisons were resolved or the focal state became untestable. Observed Ct1–Ct4 statistics were estimated with *calcConvCt()*, and their significance was evaluated with *convSigCt()* using simulation-based null distributions. Because some tree–state combinations required targeted reruns, the compiled publication table retained the

best available completed result for each tree–state combination, prioritizing successful runs with valid p-values and greater simulation depth. Table 1 reports Ct1 p-values, whereas the full Ct1–Ct4 results are presented separately in Table S11.

Third, we quantified phenotypic clustering with the Wheatsheaf index using *windex::test.windex*. We conducted this analysis separately for each ecological group on each of the four trees using PC1–PC4 and 1,000 resampling replicates. For each state, we recorded the raw Wheatsheaf index and its associated p-value, then summarized results across trees using the median index and median p-value. In the integrated summary table, higher Wheatsheaf values indicate stronger clustering of focal taxa in morphospace, and significant p-values indicate stronger phenotypic similarity among focal taxa than expected under the phylogenetic background.

***Multi-tree synthesis and interpretation***

For all multi-tree summaries, tree-specific estimates were combined across the four main phylogenies by taking means for continuous metrics such as root distance, CV, and RRphylo p-values, and medians where appropriate for the Wheatsheaf index. Interpretive categories shown in the final synthesis table were assigned from the combined evidence rather than from any single metric alone. In this framework, proximity to the reconstructed root, low within-group disparity, high phenotypic clustering, and non-derived OU optima were interpreted as evidence of morphological conservatism, whereas greater displacement from the root and larger theta–root values were interpreted as evidence of derived divergence. Convergence metrics were evaluated jointly with these root- and disparity-based summaries to distinguish repeated evolution toward similar forms from simple retention of ancestral morphology.

For visualization of ecological structure in morphospace, species were plotted in skull-shape PC space with focal habitat or diet classes highlighted against the broader dataset. Two-dimensional panels used PC1 and PC2 with group-specific ellipses, whereas three-dimensional panels used PC1–PC3. Representative skull exemplars were overlaid in some diet morphospace panels to illustrate the range of cranial forms associated with each trophic guild; these exemplars were used for visualization only and were not part of the statistical tests.

### Data S1: Annotated classification of Eupercaria

#### Key
Bet2013 = sensu Betancur-R et al. 2013 [1]
Bet2017 = sensu Betancur-R et al. 2017 [2]
Ghez2022 = sensu Ghezelayagh et al. 2022 [3]
NT2024 = sensu Near and Thacker 2024 [4]
NS = not sampled
NM = not monophyletic

See also Table S3: Spreadsheet comparing historical classifications to our revised classification (Excel, separate document)

#### Philosophy and approach to classification
The classification we present herein is updated from Bet2017, which itself was an update of several prior sources[1,5]. This scheme incorporates advances in our understanding of the relationships of fishes based on phylogenomic-scale datasets, which have generally provided improved resolution and stability of these relationships since the multi-locus datasets of the 2010's. Our goal is a classification that reflects current phylogenetic evidence, while recognizing that phylogenetic classifications are dynamic hypotheses and must become updated as new data become available. As genome-scale studies continue to refine phylogenetic relationships, we expect future changes to become progressively fewer and more focused. Additionally, we acknowledge the importance of considering phylogenetic uncertainty by evaluating not only our own results but also robust, if sometimes discordant, phylogenomic inferences from other studies.

We used the following guidelines to evaluate and update classification. If the name was included in Bet2017 and its monophyly persists with phylogenomic-scale datasets, then we chose to keep that name at its previous rank (i.e. order or suborder); for example, keeping Tetraodontiformes and Lophiiformes as orders. One exception was the elevation of some infraorders to suborders (e.g. cottoids, zoarcoids), which was done to improve consistency with other taxonomic sources (see section "Perciformes" below). We chose to name new clades whenever a clade not named in Bet2017 has since stabilized, appearing across various molecular datasets (e.g. exons and UCE's) and concatenation-based and summary coalescent analyses (e.g. IQ-TREE and ASTRAL). Therefore, our own analyses were not the only source of information, as we prioritized naming clades that also appear in other studies (e.g. Ghez2022). These clades are more likely to remain stable in the future as phylogenetic approaches improve, as they are supported by markers distributed across the genome. While we could have left some of these clades nameless (e.g. unassigned to a suborder), naming these clades communicates their stability to other practitioners.

Where hierarchical relationships are not resolved, some groups are left "*incertae sedis*" (of uncertain placement) or "*incertae mutabilis*"[6] (of changeable placement) at the suborder, order or superorder level, communicating that these remain unstable even with phylogenomic-scale datasets or require improved taxonomic sampling to place. While this practice has been criticized[3,4], we believe that leaving taxa assigned to higher ranks is itself a form of communication, recognizing the state of our understanding of the interrelationships of these fishes and highlighting priorities for future investigation. For example, consider a monophyletic group A, with strong evidence of monophyly, and a smaller group B, closely related to A, but with unclear placement within it, though A and B together are monophyletic relative to sister group X.

Retaining B as *incertae sedis* reflects this uncertainty, providing more information than forcing B into A (lumping) or separating it entirely (splitting).

**Superorders instead of orders**

Both exons (this study) and UCEs (Ghez2022) unequivocally support the division of Eupercaria into five clades. Given that these clades are strongly supported and their validity is not in question, the taxonomic rank assigned to these clades depends on interpretative decisions regarding their placement within the hierarchy and the utility of this assignment for elucidating the group's evolutionary history. One might argue that this decision is subjective. However, taxonomic schemes are "real" in the sense that they are tools for communicating about organisms and their hypothesized relatedness, and the effectiveness of any one scheme is measurable by the number of practitioners that adopt it. Therefore, researchers should develop and use taxonomic schemes that will most effectively communicate biological information of interest.

Ghez2022/NT2024 recognized these five clades at the level of taxonomic orders, following a similar scheme proposed by Davis and Smith[7]. To maintain a ranked structure, clades nested within these orders could not themselves be recognized as orders. Consequently, several traditionally recognized orders within these clades needed to be relegated to suborders (e.g. Lophiiformes to "Lophioidei"; Tetraodontiformes to "Tetraodontoidei"), suborders to superfamilies (e.g. Ceratioidei to "Ceratioidea"), and so on. A broader source of confusion arises from the re-circumscription of traditionally recognized suborders to accommodate groups that earlier classifications, as well as our own, recognize as orders. For example, Ghez2022/NT2024 uses Tetraodontoidei to refer to Tetraodontiformes (encompassing nine families), whereas Tetraodontoidei has traditionally referred to a more restricted grouping of two families (Diodontidae and Tetraodontidae). This expansion of suborders to subsume orders adds to ambiguity and introduces unnecessary complexity to the classification. While this scheme names monophyletic groups and is therefore valid, it initiates a widespread cascade of taxonomic changes with unknown consequences for communication among scientists, natural resource managers, educators and the general public. As we argue herein, this cascade is unnecessary and avoidable.

In this study, we recognize these five clades as superorders, each with the suffix "-morphae". This allows us to keep traditional order-level names, provided they apply to groups that remain monophyletic and stable across different molecular markers and phylogeny estimation approaches. There is precedence for recognizing acanthopterygian clades of similar age and size as superorders. Bet2017 divided Ovalentaria, a series of Percomorpha second only to Eupercaria in diversity, into four superorders: Cichlomorphae, Atherinomorphae, Mugilomorphae, and Blenniimorphae. This communicated that Ovalentaria was comprised of four major clades, while also retaining traditional order-level names such as Atheriniformes, Beloniformes, Cyprinodontiformes, and Gobiesociformes. Unlike Ovalentaria, Bet2017 did not name superorders within Eupercaria due to limited resolution from multi-locus data. Phylogenomic-scale datasets have now provided this resolution. As Wiley and Lieberman[8] emphasized, taxonomic categories are genealogical constructs that express relative subordination within a clade rather than biological distinctiveness; assigning superorders to these monophyletic groups reflects their relative relationships and improves the logical consistency of the classification.

In summary, both Ghez2022 and our study recognize five major clades of Eupercaria, with Ghez2022/NT2024 considering these to be orders while we recognize them as superorders. We argue that this new approach of treating these clades as superorders communicates the newfound stability of relationships within Eupercaria while also retaining traditional order- and suborder-level names, provided they apply to groups that are monophyletic and stable. This solution therefore avoids the cascade of taxonomic revisions implicated by the Ghez2022/NT2024 scheme.

**Additional comparison to Near and Thacker (2024)**

As phylogenomics has given us unprecedented clarity on the interrelationships of fishes (Fig. S5), we now must reconcile historical classification with this new understanding[9]. This differs from other vertebrate groups for which order-level classification has remained stable[10]. While classifications reflect objective phylogenetic relationships, the choice to use categorical ranks or alternative methods to denote these relationships involves interpretive decisions. Given these interpretive choices, it is unsurprising that different groups of researchers generate different schemes based on the same underlying phylogenetic relationships. Nonetheless, the communicative utility of any scheme depends on the number of practitioners that adopt it, all else being equal. Below we clarify the differences between our scheme and that of Ghez2022/NT2024 so that researchers can choose to adopt the classification scheme that works best for their purposes.

Ghez2022 presented a new classification scheme for spiny-rayed fishes (Acanthomorpha), which was expanded to all fishes in NT2024. This scheme overlaps with Bet2017, including recognition of five major clades of Eupercaria (see above). The names "Centrarchiformes" and "Perciformes" describing two of these clades were shared with Bet2017. Similarly, a third clade was recognized at the order level, "Acropomatiformes" (equivalent to our "Pempheriformes"). Therefore, our classification schemes agree in recognizing three of these five major clades as taxonomic orders. In our scheme, these ordinal names are synonymous with their corresponding superordinal names (i.e. the superorders Percimorphae, Centrarchimorphae, and Pempherimorphae contain one order, Perciformes, Centrarchiformes, and Pempheriformes respectively).

The most obvious differences between our schemes relates to the remaining two major clades, termed "Labrimorphae" and "Lophiimorphae" (our study) or "Labriformes" and "Acanthuriformes" (Ghez2022/NT2024). In the latter case, 11 taxonomic orders recognized by Bet2017 were subsumed into a single order "Acanthuriformes". Three of the previous orders were relegated to suborders (Tetraodontoidei, Lophioidei, and Acanthuroidei), implying a cascade of taxonomic changes at lower levels (see above). The remaining orders were left unnamed, even though most of these dissolved names apply to stable monophyletic groups. "Labrimorphae" represents a similar case where the order "Uranoscopiformes" was sunk into Labriformes and left unrecognized despite being a stable clade. The propensity to "lump" is also reflected within Perciformes, as the Ghez2022/NT2024 scheme recognizes three suborders that constitute seven suborders in the Bet2017 scheme and 11 suborders in our updated scheme.

These observations demonstrate that the most significant difference between these two classification schemes is the tendency to "lump" versus "split", where "lumping" is the practice of applying names to fewer, more inclusive clades. Both classification schemes name stable monophyletic groups (Fig. S5); therefore, both schemes are valid[11]. Although it has been a focal topic in fish systematics[9], we do not discuss the merits of ranked versus rank-free classification approaches here (but see refs. [8,12]) because it is not the most consequential source of disagreement between these schemes (e.g., two schemes can be equally rank-free but disagree on the number of named clades and their size). Instead, we hereafter discuss the merits of lumping versus splitting and the resulting utility of these schemes.

Ghez2022 gave two justifications for their approach to classification. First, their scheme assigns every family to an order, and no families are left as *incertae sedis.* We have addressed this point above, specifically our belief that "*incertae sedis*" benefits communication by explicitly conveying phylogenetic uncertainty. Second, their taxonomic orders contain more families on average (18.2 families per order) than the Bet2017 scheme (7.4 families per order). They believed these low-diversity orders were less phylogenetically informative than inclusive orders. We disagree with this claim for two reasons: (1) low-diversity ranks reflect the underlying process of lineage diversification, and (2) ranks can become *less* phylogenetically informative as they become more inclusive.

Most clades contain few species[13]; high-diversity clades are celebrated because they are exceptional[14,15]. This pattern, termed the "hollow curve", is ubiquitous across the Tree of Life[16]. As long as ranked names are applied to monophyletic groups, then aggregate properties of ranks should approximate those of the clades they describe, such as recapitulating the hollow curve. In other words, low-diversity ranks should be more common than high-diversity ranks. For example, 10 of the 27 mammalian orders contain fewer than 10 species[17], such as Tubulidentata (aardvarks), Dermoptera (colugos), and Proboscidea (elephants). Eliminating low-diversity orders was an explicit goal of Ghez2022/NT2024, even though low-diversity clades are expected when diversification rates vary substantially across clades[18].

A feature of ranked classifications is communicative power about interrelationships; a clades's name implies information meant to be communicated with others. For example, families within an order are more closely related to each other than to families in a different order, assuming monophyly of orders. The order "Spariformes" communicates that the families Sparidae, Nemipteridae, and Lethrinidae are related. In the scheme of Ghez2022/NT2024, "Spariformes" is dissolved into a broader circumscription of Acanthuriformes. Under this scheme, practitioners can glean that Sparidae is more closely related to Nemipteridae and Haemulidae than to Centrarchidae (because they share an order), but cannot tell whether Sparidae is more closely related to Nemipteridae or to Haemulidae without referring to the phylogeny. Therefore, communicative power of the classification scheme is lost by dissolving Spariformes. The more families included within an order, the less information is conveyed about their interrelationships. Grouping units into lower ranks solves this problem (e.g. "Acanthuroidei" links Acanthuridae, Zanclidae and Luvaridae) but can introduce a new problem of prompting a cascade of taxonomic changes (discussed above). Note that orders that contain a single family (e.g. "Gerreiformes") may still be useful for communicating that a lineage is phylogenetically unique.

A "good" taxon name is predictive and stable[19]: predicting information about similarly and relatedness among lower units, and likely to persist in the future. A benefit of inclusive names is that they are inherently more stable, as they require less information about the interrelationships of constituent taxa[20]. For example, it is unclear whether Haemulidae and Lutjanidae are sister families, but we can at least be confident they both belong to the clade that also contains Tetraodontidae and Lophiidae. Still, we believe that our understanding of the systematics of fishes has advanced beyond this stage. Most interrelationships among higher taxa are concordant between exons (our study) and UCEs (Fig. S5). We have an unprecedented degree of insight into the phylogeny of fishes, and this can be communicated with classification.

In summary, we present an updated classification of Eupercaria that emphasizes concordance among phylogenomic-scale studies. Our scheme differs from Ghez2022/NT2024 by keeping traditional orders (provided they are monophyletic) and naming new clades to communicate phylogenetic resolution. We disagree with Ghez2022/NT2024 that less-inclusive taxon names are phylogenetically uninformative, and we believe that the cascade of taxonomic changes implied by their scheme are unnecessary.

**Superorder Percimorphae**
Equal to: Perciformes (Ghez2022/NT2024)

Percimorphae is consistently found to be sister to the clade formed by the other four superorders of Eupercaria (Figs. S2–S5).

### Order Perciformes

We use "Perciformes" consistently with Bet2013/Bet2017. A monophyletic definition of Perciformes was proposed by Bet2013 to eliminate the wastebasket circumscription still common in ichthyology textbooks[21,22], and is now widely accepted[3,4,23–25]. The order Perciformes and several suborder names are shared with the classification scheme of Ghez2022/NT2024 (e.g. Percoidei, Notothenoidei).

Many names at various ranks have been applied to clades within Perciformes (Table S3), and outdated names like "Scorpaeniformes" appear as recently as 2016[22]. This nomenclatural instability gives a false impression of phylogenetic uncertainty within Perciformes. In actuality, many relationships are consistent across molecular datasets (Fig. S2–S7) and stable clades can be recognized.

### Note on "Serranidae"

Our study has the densest taxonomic sampling of "serranids" of any phylogenomic study to-date. We found two nominal serranid clades, which are proposed herein to be suborders (Ephinepheloidei, Serranoidei). Ghez2022 found four clades but had much lower taxonomic sampling of nominal serranids. For comparison, a table of genera nominally considered to be serranids at the time of Smith and Craig [26] and their family-level assignments across recent sources is provided in Table S3.

Bet2017 opted not to break apart Serranidae (other than Niphonidae) since their multi-locus analysis recovered a monophyletic Serranidae exclusive of *Niphon.* However, our analyses and others strongly reject a monophyletic Serranidae (Figs. S2–S5). The non-monophyly of Serranidae has long been recognized using molecular datasets[26]. Smith and Craig[26] found that nominal serranids were polyphyletic and distributed across five clades: *Niphon*, *Acanthistius*, Epinephelinae, Anthiinae, and Serraninae. All but Anthiinae are recognized in our classification scheme.

**Changes since Bet2017:** Family and suborder recognized: Ephinepheloidei/Epinephelidae.

**Notes on *Acanthistius*:** The genus *Acanthistius* has long been resolved as a phylogenetically distinct lineage in molecular datasets[3,26,28]. Although Fowler[27] previously placed *Acanthistius* in its own subfamily, Acanthistiinae, we do not recommend elevating this group to the family level at this time. The genus has experienced considerable taxonomic instability, having been resolved as *incertae sedis* within Perciformes[26] and as sister to a clade comprising Anthididae and Epinephelidae[3]. Rabosky et al.[23] resolved *Acanthistius* as paraphyletic, with *A. cinctus* sister a large clade comprising Percophidae, Bovichthidae, Elignopidae and others, and *A. ocellatus* sister to the genus *Niphon*[23]. Our current sampling includes only a single species, *A. patachonicus*, which represents a highly divergent branch within the genus (Y. Menkara, pers. comm.). Given both the limited taxonomic sampling and the uncertainty surrounding the placement of *Acanthistius*, additional genomic data and broader species representation are required before the lineage can be confidently recognized at the family rank. Consistent with previous work, the family- and order-level placements of *Acanthistiu*s within Percomorpha remains uncertain (Figs. S2–S5).

### Suborder Epinepheloidei

Contains: Epinephelidae

The subfamily Epinephelinae was first established by Bleeker (1874a) during his revision of Indo-Archipelagic species of Epinephelini and related genera. Later, Smith and Craig[26] formally elevated the group and recognized Epinephelidae as a distinct family.

Note that our "Epinephelidae" contains genera called "Anthiadidae" by Ghez2022 and "Clade D: Anthiinae" by Smith and Craig[26] (see Table S3). Unlike these studies, we found that the "epinephelines" and "anthiines" form a monophyletic group. We therefore opted not to recognize the anthiines as a separate family.

### Suborder Serranoidei

Contains: Serranidae *sensu stricto*

Our circumscription of Serranidae *sensu stricto* is consistent with refs[3,24,26,28] (Table S3).

### Suborder Bembropoidei

Includes: Bembropidae (*Bembrops, Chrionema*)

This suborder was named by Bet2017 to recognize that these genera (formerly Percophidae) were unrelated to *Percophis*. Our IQ-TREES (but not ASTRAL trees) find Bembropidae to be sister to Serranidae consistent with Ghez2022 (Figs. S2–S5).

### Suborder Percoidei

Equal to: Percoidei of Ghez2022/NT2024

Includes: Percidae, Trachinidae, Niphonidae (NS)

This is a highly stable clade across our analyses and others (Fig. S2–S7). Molecular datasets have long found *Niphon* to be more closely related to percids instead of other serranids[3,23,26]. Smith and Craig[26] resurrected Niphonidae, originally proposed by Jordan (1923), which had already been considered a percoid in the classification scheme of Bet2017.

### Suborder Notothenoidei

Equal to: Notothenoidei of Ghez2022/NT2024

Includes: Percophidae (*Percophis*), Bovichtidae, Pseudaphritidae, Eleginopidae, Channichthyidae, Bathydraconidae, Harpagiferidae, Nototheniidae (NM). We follow Parker and Near [29] by sinking Artedidraconidae into Harpagiferidae.

Interrelationships of families within the notothenioid lineage are stable across our analyses and others (Fig. S2–S7). *Percophis* is now widely accepted to be a notothenioid[2,28]. We found a non-monophyletic Nototheniidae with *Notothenia* being more closely related to Channichthyidae, Bathydraconidae, and Harpagideridae than other nototheniids. This is consistent with studies using other molecular markers[3,30].

### Suborder Platycephaloidei

Includes: Platycephalidae

**Changes since Bet2017:** "Platycephaloidei" is now restricted to Platycephalidae. "Platycephaloidei" previously contained five families: Platycephalidae, Bembridae, Parabembridae, Hoplichthyidae, and Plectrogeniidae. In the phylogenomic era, neither exons (this study) nor UCEs (Ghez2022) support this clade. Platycephalidae, Bembridae, and Hoplichthyidae are each phylogenetically unique lineages with unclear affinities (Figs. S2–S4). Plectrogeniidae is unequivocally a scorpaenoid (see below).

Our analyses do not support Platycephalidae as a member of Scorpaenoidei sensu Smith et al.[31].

### Suborder Bembroidei

Includes: Bembridae, Parabembridae. Note that some sources do not recognize Parabembridae as a separate family[3,31]. These families are reciprocally monophyletic in our analyses.

**Changes since Bet2017:** "Platycephaloidei" previously contained Bembridae and Parabembridae. Similar to Platycephalidae, the clade formed by Bembridae+Parabembridae has been found to be a phylogenetically unique lineage using phylogenomic-scale datasets (exons and UCEs[3]).

Our analyses do not support Bembridae as a member of Scorpaenoidei sensu Smith et al. [31].

### Suborder Hoplichthyoidei

Includes: Hoplichthyidae

**Changes since Bet2017:** "Platycephaloidei" previously contained Hoplichthyidae. Similar to Platycephalidae, Hoplichthyidae has been found to be a phylogenetically unique lineage using phylogenomic-scale datasets (exons and UCEs[3]).
Hoplichthyidae is sometimes found to be sister to Normanichthyidae (our IQ-TREEs and Ghez2022), but this relationship is not supported when using ASTRAL (Figs. S2–S5).

Our analyses do not support Hoplichthyidae as a member of Scorpaenoidei sensu Smith et al. [31].

### Suborder Scorpaenoidei

**Changes since Bet2017:** Plectogeniidae is now assigned to this suborder rather than Platycephaloidei

Includes: Neosebastidae, Plectrogeniidae, Scorpaenidae, Synanceiidae. Our results support family-level taxonomic changes made by Smith et al. [31], specifically sinking Sebastidae and Setarchidae into Scorpaenidae, and sinking Aploactinidae, Apistidae and Tetrarogidae into Synanceiidae.

The following families were not sampled, but also assumed to be part of Scorpaenoidei following Bet2017: Congiopodidae, Eschmeyeridae, Gnathanacanthidae, Pataecidae, Perryenidae, Zanclorhynchidae. Other molecular studies support the inclusion of Congiopodidae[3,26,31] (but see ref. [23]), Pataecidae[23,26,31], and Zanclorhynchidae[23,26,31]. Smith et al. [31] sunk Eschmeyeridae, Gnathanacanthidae, Perryenidae and Pataecidae into Synanceiidae.

Smith et al. [31] proposed a suborder Scorpaenoidei consisting of: Platycephalidae, Hoplichthyidae, Triglidae, Bembridae, Synanceiidae, Neosebastidae, Plectrogeniidae, and Scorpaenidae *sensu lato*. Neither our analyses nor those of Ghez2022 using UCEs support this circumscription (Fig. S5). The families Bembridae, Hoplichthyidae, Platycephalidae, and Triglidae do not show a consistent association with the other scorpaenoid families. Rather, these four families are part of a larger monophyletic group with the Cottoidei, Zoarcoidei, Gasterosteoidei, Anoplopomatoidei, and Normanichthyoidei. While this clade is stable across exons and UCEs and using concatenation-based and coalescent analyses, the interrelationships within it are unstable (Figs. S2–S5).

Ghez2022 termed this large clade "Scorpaenoidei." This name has validity as it refers to a stable monophyletic group. However, we prefer to use a restricted circumscription of "Scorpaenoidei" in order to continue applying suborder-level names to other lineages within this large clade (e.g. "Cottoidei"[32,33] and "Zoarcoidei"[34]). The clade formed by Neosebastidae, Plectrogeniidae, Scorpaenidae, and Synanceiidae is largely stable (Figs. S2–S5); we recognize this stability by applying the name "Scorpaenoidei" following previous classifications[2]. We believe such "splitting" aids communication (see above).

### Suborder Trigloidei

Includes: Triglidae. Our results support sinking Peristediidae into Triglidae[31].

Our analyses do not support Triglidae as a member of Scorpaenoidei sensu Smith et al. [31]. Our usage of Trigloidei is identical to Bet2017.

### Suborder Normanichthyoidei

Includes: Normanichthyidae

Normanichthyidae is a phylogenetically unique lineage within the large clade called "Scorpaenoidei" sensu Ghez2022. Its sister group is unclear (Figs. S2–S7). Our usage of Normanichthyoidei is consistent with Bet2017.

### Suborder Cottoidei

Includes: Hexagrammidae, Liparidae, Cyclopteridae, Rhamphocottidae, Ereuniidae, Psychrolutidae, Cottidae, Agonidae, Jordaniidae (NS), Trichodontidae (NS), Zaniolepididae (NS). The inclusion of the latter four families is supported by other molecular studies[2,3,26,33].

**Changes since Bet2017:** "Cottoidei" had a larger circumscription, referring to the clade containing Anoplopomatidae, Gasterosteales, Zaniolepididae, Hexagrammidae, Cottales (Cottidae and allies), and Zoarcales (Zoarcidae and allies). This clade has proven stable in phylogenomic analyses of exons (our study) and UCEs (Ghez2022, who considered this clade part of Scorpaenoidei). Nonetheless, we chose to elevate infraorders of Bet2017 to suborders. This usage is more consistent with other taxonomic authorities (e.g. [5,22,24–26,35]; Table S3) and appears frequently across the literature[32–34].

"Cottoidei" is restricted to the former infraorders Cottales, Hexagrammales, and Zaniolepidoales of Bet2017 to acknowledge that Hexagrammidae and Zaniolepididae are consistently found to be affiliated with the other cottoid families (Figs. S2–S7 and refs.[3,33]). Interrelationships among families within Cottoidei are remarkably stable across molecular datasets and analyses (see also ref.[33]).

### Suborder Anoplopomatoidei

Includes: Anoplopomatidae

**Changes since Bet2017:** Anoplopomatidae was formerly an infraorder-level clade "Anoplopomatales" within the suborder Cottoidei. We elevated it to a suborder along with the Gasterosteoidei (="Gasterosteales"), Zoarcoidei (="Zoarcales"), and Cottoidei (="Cottales"+ "Hexagrammales"+"Zaniolepidoales").

Anoplopomatidae is a phylogenetically unique lineage within "Cottoidei" *sensu lato* (Bet2017), justifying its recognition as a suborder (Figs. S2–S7).

### Suborder Gasterosteoidei

Includes: Aulorhynchidae, Gasterosteidae, Hypoptychidae

**Changes since Bet2017:** Elevated infraorder "Gasterosteales" to a suborder

### Suborder Zoarcoidei

Includes: Bathymasteridae, Zoarcidae, Anarhichadidae, Cryptacanthodidae, Zaproridae, Stichaeidae, Ptilichthyidae, Pholidae, Scytalinidae (NS; part of Stichaeidae in Ghez2022) Eulophiidae (NS), Neozoarcidae (NS)

**Changes since Bet2017:** Elevated infraorder "Zoarcales" to a suborder

The monophyly of Stichaeidae is questionable[3,34]. Taxonomic revision of families within the zoarcoids is necessary. Our species-level sampling is not sufficient to recommend changes (but see refs.[3,34]).

**Superorder Centrarchimorphae**
Equal to: Centrarchiformes (Ghez2022/NT2024)

### Order Centrarchiformes

The name "Centrarchiformes" was first used by Bet2013 to group Centrarchidae+Elassomatidae (with Elassomatidae formerly called "Elassomatiformes" by Wiley and Johnson[5]). It has been used consistently since 2017 (e.g. refs.[2–4,23,36]).

In this study, all five suborder names are retained from Bet2017, although their circumscriptions have changed. A new suborder "Latroidei" is recognized herein. We follow Ghez2022 by splitting Percichthyidae into the monophyletic families Sinipercidae (*Siniperca, Coreoperca*) and Percichthyidae (*Maccullochella, Bostockia, Nannoperca, Percilia, Macquaria, Guyu*).

### Suborder Percalatoidei

Contains: *Percalates* (NS)

*Percalates* is a phylogenetically unique lineage. Analyses of UCEs suggest it is the sister lineage to all other centrarchiforms (Ghez2022/NT2024). *Percalates* was not given a family-level assignment in recent compilations (Bet2017/NT2024).

### Suborder Terapontoidei

Contains: Terapontidae, Kyphosidae, Kuhliidae, Oplegnathidae, Scorpididae (NM), Girellidae (NS), Distichiidae (NS), Microcanthidae (NS)

This clade is highly stable across our analyses and others (Figs. S2–S7). The inclusion of Girellidae, Distichidae, and Microcanthidae follows Bet2017 and is supported by other molecular analyses[3,36]. Scorpididae is not monophyletic, but the scorpidid genera *Atypichthys* and *Scorpis* are always found within this clade (Figs. S2–S5).

The name "Terapontiformes" has been used previously to describe this clade[1,2]. Given its stability and that it is consistently found to be sister to all remaining centrarchiforms, it may be justifiably recognized as an order-level clade. However, we keep "Centrarchiformes" here for consistency with other studies.

### Suborder Cirrhitoidei

**Changes since Bet2017:** Cirrhitoidei is now restricted to Cirrhitidae to accommodate this family's inconsistent placement within Centrarchiformes. Cirrhitidae is seemingly a phylogenetically unique lineage. "Cirrhitoidei" previously referred to a clade containing Aplodactylidae, Cheilodactylidae, Chironemidae, Cirrhitidae, and Latridae. This clade is not consistently supported by our analyses. It is found in our IQ-TREEs (Fig. S3) and several past studies[2,3,23,36,37]. However, all of our ASTRAL trees (Fig. S4) find Cirrhitidae to be a phylogenetically unique lineage sister to the clade formed by Centrarchoidei, Latroidei, and Percichthyoidei (see below).

Contains: Cirrhitidae

The assignment of Cirrhitidae to its own suborder Cirrhitoidei is taxonomically stable given the potential range of positions of Cirrhitidae seen across analyses (see above).

### Suborder Latroidei

**Changes since Bet2017:** New suborder erected to recognize the stability of this clade

Contains: Latridae (NM), Chironemidae, Cheilodactylidae, Aplodactylidae

These four families were previously assigned to Cirrhitoidei[2,37]. The assignment of Cirrhitidae to its own suborder accommodates its unstable position. In contrast, the clade containing these four families is highly stable (Figs. S2–S7).

Latridae (*Latris, Mendosoma*) is consistently non-monophyletic in our analyses with Cheilodactylidae (*Cheilodactylus, Nemadactylus*) nested within. The non-monophyly of Latridae is well known. Burridge and Smolenski [38] suggested Cheilodactylidae be subsumed within Latridae. Using denser taxonomic sampling than our study, Ludt et al. [37] considered *Nemadactylus* to be part of a monophyletic Latridae, and Cheilodactylidae monophyletic when restricted to *Cheilodactylus*.

### Suborder Centrarchoidei

**Changes since Bet2017:** Enoplosidae now in Percichthyoidei

Contains: Sinipercidae, Elassomatidae, Centrarchidae

This clade is highly stable across our analyses and others (Figs. S2–S7 and refs.[39–41]). Some authors consider Elassomatidae to be part of Centrarchidae[3].

### Suborder Percichthyoidei

**Changes since Bet2017:** Now includes Enoplosidae

Contains: Enoplosidae, Percichthyidae (Percichthyidae *sensu stricto* includes *Percilia* formerly placed in its own family, Perciliidae).

Although not consistently found in previous multilocus studies[2,23] (but see refs.[39,40]), this clade is highly stable across phylogenomic-scale analyses of exons and UCEs (Figs. S2–S5).

### *Incertae sedis* within Centrarchimorphae

Contains: Parascorpididae[2,4,35] (NS).

*Parascorpis* has not been sampled in recent molecular phylogenetic analyses.

### Superorder Pempherimorphae

#### Order Pempheriformes

Equal to: Acropomatiformes (Ghez2022/NT 2024)

**Changes from Bet2017:** Relationships within this group have stabilized across phylogenomic-scale studies using UCEs and exons (our study and refs. [3,42]). For example, the interrelationships among families are identical between our all-genes IQ-TREE and Ghez2022 (Fig. S5) after excluding rogue families Creediidae+Champsodontidae. Therefore, we name several suborder-level clades to recognize this stability. Smith et al. [42] named one suborder, Acropomatoidei, which is shared by our study. Bet2017 declined to name suborders within Pempheriformes.

Since 2017, the families Leptoscopidae and Percophidae are no longer believed to be in this order. Leptoscopidae has been re-assigned to Uranoscopiformes. The genus *Acanthaphritis* (formerly Percophidae) is now assigned to Hemerocoetidae, and Percophidae is restricted to *Percophis* which is in Perciformes (see references above).

Smith et al.[42] is presently the most densely sampled phylogenomic-scale analysis of Pempheriformes/Acropomatiformes. We follow Smith et al.[42] by splitting Acropomatidae into three families to resolve its monophyly: Acropomatidae (*Acropoma* and *Doederleinia*), Malakichthyidae (*Malakichthys* and *Hemilutjanus*), and Synagropidae (*Synagrops*). Polyprionidae now contains just *Polyprion* as Smith et al. placed *Stereolepis* into its own family Stereolepididae (not sampled here).

Creediidae and Champsodontidae are believed to belong to this group[2,3,42], but in our study (and others[42]) they are probably affected by long branch attraction, therefore we do not assign them to a suborder pending additional investigation.

Note that several families are not sampled in this study (see below). This group is a priority for future sampling. Sampling has been historically challenging since this group is primarily found in the deep sea.

#### Justification for "Pempheriformes" over "Acropomatiformes"

The order-level name "Acropomatiformes" has been frequently applied to this group[3,4,7,42,43]. This name seemingly dates to 2015 with the classification by Smith and Davis[7]. The name "Pempheriformes" has been in use slightly longer, since Bet2013[1], to refer to the clade containing Pempheridae and Glaucosomatidae which also has some morphological support[2]. The circumscription of Pempheriformes was updated in Bet2017 to include many families that are generally agreed to make up this clade[42].

Smith et al.[42] wrote in support of "Acropomatiformes" over "Pempheriformes". We did not find these arguments compelling enough to warrant changing this order's name. While Katayama initially recognized a grouping called the "*Acropoma*-stem clade" in 1959[44], this does not equate to naming an order-level clade. The first instance of this clade being recognized as an order was in 2013 under the name "Pempheriformes"[1]. Smith et al. further argued against use of "Pempheriformes" because the family Pempheridae was found outside of this clade in the 'megatree' analysis of Rabosky et al.[23], and suggested that the position of Acropomatidae was more stable. However, many relationships in Rabosky et al.'s 'megatree' are not held up by phylogenomic-scale studies (e.g. Fig. S6). NT2024 later defined Acropomatiformes as the least inclusive crown clade that contains *Acropoma japonicum* and *Pempheris schomburgkii,* indicating that the position of Pempheridae is indeed stable enough to define this clade. In summary, we do not see any compelling reason to use "Acropomatiformes" over "Pempheriformes" when the latter has been in use longer.

### Suborder Acropomatoidei

Equal to: Acropomatoidei of Smith et al.[42]

Includes: Acropomatidae, Epigonidae (NM), Symphysanodontidae, Howellidae (NS), Ostracoberycidae (NS), Scombropidae (NS)

Our analyses confirm a stable grouping of Acropomatidae, Epigonidae, and Symphysanodontidae (Figs. S2–S5), which had also been found by Bet2017 (Fig. S7). While Epigonidae (represented in our analyses by *Epigonus* and *Sphyraenops*) is not always monophyletic (Figs. S3, S4), all of its members were found to be within Acropomatoidei, which is confirmed by other studies[3,36] (including *Brinkmannella*[3]).

Inclusion of Howellidae, Ostracoberycidae, and Scombropidae, within this suborder is supported by other molecular studies[3,36,42].

### Suborder Bathyclupeoidei

Includes: Bathyclupeidae, Synagropidae

Bathyclupeidae and Synagropidae are consistently found to be sister families across our analyses (Fig. S2–S4). This sister-group relationship is corroborated by Smith et al.[42]. Ghez2022 did not sample *Synagrops*.

### Suborder Dinolestoidei

Includes: Dinolestidae (NS)

Dinolestidae is a phylogenetically unique lineage. Smith et al.[42] found it to be sister to a clade formed by Stereolepididae, Bathyclupeidae and Synagropidae, but membership of Stereolepididae within this clade had low bootstrap support. Ghez2022 did not sample *Dinolestes.*

### Suborder Malakichthyoidei

Includes: Malakichthyidae

*Malakichthys* is a phylogenetically unique lineage. Its sister group is unclear (Figs. S2–S4). It was not sampled by Ghez2022. Smith et al.[42] found it to be sister to *Hemilutjanus* (not sampled here) and therefore believed *Hemilutjanus* to be in the family Malakichthyidae.

### Suborder Pempheroidei

Includes: Glaucosomatidae, Lateolabracidae, Pempheridae

This is a highly stable clade (Figs. S2–S4) that has also been supported by other studies[3,42].

### Suborder Pentacerotoidei

Includes: Pentacerotidae, Banjosidae

The sister-group relationship of Pentacerotidae and Banjosidae is highly stable across our analyses (Fig. S2–S4) and other studies[2,3,36,42].

### Suborder Polyprionoidei

Includes: Polyprionidae (*Polyprion*)

*Polyprion* is a phylogenetically unique lineage that is often found to be sister to Pempheroidei (our IQ-TREEs and refs.[3,42]). However, this relationship was not found in our ASTRAL trees (Fig. S4). Note that Smith et al.[42] placed *Stereolepis* (not sampled here; see below) into its own family Stereolepididae because it was not closely related to *Polyprion.*

### Suborder Stereolepidoidei

Includes: Stereolepididae (*Stereolepis*) (NS)

*Stereolepis* is a phylogenetically unique lineage. Its sister group is unclear. Smith et al.[42] found it to be sister to Bathyclupeidae+Synagropidae with low bootstrap support. Ghez2022 found it to be sister to Banjosidae+Pentacerotidae.

### *Incertae sedis* within Pempherimorphae

Includes: Hemerocoetidae (NS), *Schuettea* (NS)

Smith et al.[42] found *Schuettea* to be related to Champsodontidae, and Hemerocoetidae to be related to Creediidae (also found by Ghez2022). Both Creediidae and Champsodontidae have very long branches in their study (UCE's) and ours (exons), and the positions of these two families is unstable at the superorder level (see below). Therefore, the positions of these families, and families related to them may be driven by systematic errors. *Schuettea* is nominally in Monodactylidae[23,24] but NT2024 declined to assign it to a family-level clade.

**Superorder Labrimorphae**
Equal to Labriformes (NT2024)

Two clades were found in all trees, corresponding to Uranoscopiformes (Bet2017) and the clade formed by Labridae and Centrogenyidae. We opted to continue recognizing two orders, Labriformes and Uranoscopiformes[1,2,7,23], rather than sinking Uranoscopiformes into Labriformes (Ghez2022/NT2024).

### Order Labriformes

Equal to: Labroidei of Ghez2022

**Changes since Bet2017:** Including Centrogenyidae in the order Labriformes

#### Suborder Labroidei

Contains: Labridae (includes taxa previously listed in Scaridae and Odacidae)

For details of the interrelationships within Labridae, we direct readers to ref. [45], which overlaps with the exon capture dataset in this study but has more extensive taxonomic sampling of labrids.

#### Suborder Centrogenyoidei

Contains: Centrogenyidae

We recognize Centrogenyidae (false scorpionfish; *incertae sedis* in Bet2017) as a member of Labriformes. Its position as the sister family to Labridae is uncontroversial in the phylogenomic era[3,4,45].

### Order Uranoscopiformes

Equal to: Uranoscopoidei of Ghez2022

**Changes since Bet2017:** Including Leptoscopidae in the order Uranoscopiformes

Contains: Ammodytidae, Leptoscopidae, Pinguipedidae (NM), Uranoscopidae, Cheimarrichthyidae (NS)

Family-level relationships within this order were consistent across trees. Uranoscopidae is the earliest diverging family (see also ref.[3]). We found that Pinguipedidae is not monophyletic, with *Pseudopercis* being sister to the clade formed by Leptoscopidae and the remaining genera of Pinguipedidae. The non-monophyly of Pinguipedidae has been previously recognized[46–48]; this family deserves further scrutiny with expanded taxonomic sampling.

The family Leptoscopidae has been historically difficult to place (ranging from Trachiniformes,

Sygnathiformes, and Pempheriformes)[49]. Our analyses confidently place it within Uranoscopiformes, as do other phylogenomic studies[3,42]. The affiliation of Leptoscopidae with this group has precedence based on both small molecular datasets[26] and morphology[4,50].

The inclusion of Cheimarrichthyidae is supported by other studies[3,23,26].

**Superorder Lophiimorphae**
Equal to: Acanthuriformes of Ghez2022/NT2024

### Justification for "Lophiimorphae" over "Acanthurimorphae" or similar

Smith and Davis[7,43] is credited with delimiting an arrangement of five clades within Eupercaria based on a multi-locus phylogeny of 11 loci. Their scheme was supported and elaborated on by Ghez2022 using UCE's. In both studies the clade discussed herein was termed "Acanthuriformes" (although "Euperciformes" appears in Supplementary Table 1 of Ghez2022 instead). Note that this order name had been used previously (and herein) with a much smaller circumscription[1,5] to include the families Acanthuridae, Luvaridae, Zanclidae.

In our study using exons, we recognize these five clades at the superorder level to retain traditional order-level names. Here we term this clade "Lophiimorphae". Note that our delimitation differs slightly from that of NT2024 because their definition would include Gerreidae, Leiognathidae, Creediidae and Champsodontidae under some phylogenetic hypotheses, whereas we keep these families as *incertae sedis* at the superorder level (i.e. not included within Lophiimorphae based on current information).

Instead of naming this clade "Acanthurimorphae" in reference to Smith and Davis[7] and Ghez2022, we chose to name it "Lophiimorphae", honoring Lophiiformes, for the following reasons. First, of all order names within this clade recognized recently, only two appear as far back as Nelson (2006): Tetraodontiformes and Lophiiformes. Specifically, all families mentioned within were members of just three orders 2006[35]: Perciformes, Tetraodontiformes, and Lophiiformes, as Perciformes was once a wastebasket taxon. The order name "Acanthuriformes" was not used until 2010[5]. Second, naming this clade "Lophiimorphae" over (e.g.) "Tetraodontimorphae" honors one of the most significant results to come out of molecular systematics of fishes: the movement of Lophiiformes from Paracanthopterygii into an apical position within the spiny-rayed fishes[9,51–53]. Ironically, the five-suborder arrangement within Lophiiformes has been quite stable, having been recognized since 1984 based on morphology[54] and supported ever since using molecules[55]. The same cannot be said of Tetraodontiformes or any other eupercarian order, for which the circumscription of suborders has been in flux for the past ten years[1,2]. Therefore, Lophiiformes are special by reminding us to be open-minded to new discoveries while still recognizing the efforts of past ichthyologists. They are arguably the group of fishes most representative of our philosophy towards classification in this study.

### Order Moroniformes

**Changes since Bet2017:** Newly recognized order

Contains: Moronidae, Sillaginidae

Bet2017 found a sister relationship between Sillaginidae and Moronidae, but with low bootstrap support (52/100) and therefore considered these families "*incertae sedis*". The relationship has since been found with higher support in other studies[3,23]: for example in Ghez2022 with a bootstrap support of 92. In our own analyses, we find this sister group relationship in the all-genes tree (bootstrap=100) and all gene-subset trees made using IQ-TREE (Fig. S3). However, we did not find this relationship using the ASTRAL trees (Fig. S4). Still, given the consistency of this relationship in our study and others', we recognize the order Moroniformes containing Sillaginidae and Moronidae herein.

The order name "Moroniformes" has been used previously although with very different circumscriptions. The latest edition of Fishes of the World[22] applied the name Moroniformes to a clade containing Moronidae, Ephippidae, and Drepaneidae, and placed Sillaginidae within Spariformes (note that we do not find support for these relationships). Smith and Craig[26] named a suborder "Moronoidei" as the MRCA between *Morone* and *Polyprion,* grouping 56 families together that we now understand to be spread across the acanthomorph phylogeny (e.g. Acropomatidae, Carangidae, Centrarchidae, Chiasmodontidae, Mullidae, Sparidae) in addition to Moronidae and Sillaginidae.

### Order Ephippiformes

Contains: Ephippidae, Drepaneidae

We continue to recognize the order Ephippiformes sensu Bet2013/Bet2017. This sister-group relationship has been consistently found across molecular studies[1–3,23]. Here we find it in both all-genes trees (IQ-TREE and ASTRAL) and all gene-subset trees.

### Order Spariformes

Contains: Sparidae, Lethrinidae, Nemipteridae

We continue to recognize Spariformes sensu Bet2013/Bet2017. This is a highly stable clade across molecular datasets. The relationships (Nemipteridae,(Lethrinidae,Sparidae)) were found by Bet2013/Bet2017 and Ghez2022, although Ghez2022 did not name this clade. These relationships were found in both all-genes trees (IQ-TREE and ASTRAL) and most gene-subset trees (in a few gene-subset trees, Nemipteridae is found outside of this clade). Note that Centracanthidae (*Centracanthus* and *Spicara,* not sampled here) is now considered to be a synonym of Sparidae[36].

### Order Lobotiformes

Contains: Lobotidae, Hapalogenyidae, Datnioididae (NS)

We continue to recognize Lobotiformes sensu Bet2017. A clade containing Lobotidae and Hapalogenyidae was found in all analyses, and in other studies[3,23]. We refer to other studies while considering Datnioididae (not sampled) to be within Lobotiformes[2,26,36]. Note that several sources consider Hapalogenyidae and Datnioididae to be synonymous with Lobotidae[3,24,25,56]. Ghez2022 found the monodactylid *Monodactylus* (not sampled) to be sister to Lobotiformes.

### Order Lophiiformes

Equal to: Lophioidei of Ghez2022/NT2024

We retain the traditional order-level name Lophiiformes over the suborder-level Lophioidei of Ghez2022. The five-suborder arrangement within Lophiiformes is among the most stable arrangements across Eupercaria, as the monophyly of all five clades has been resolved in all molecular studies to date (see ref.[55] for a recent overview of the systematics of Lophiiformes). Therefore, we believe that relegating Lophiiformes to a suborder and its suborders to superfamilies[3,4,57] represents an unnecessary taxonomic overhaul of one of the most stable eupercarian clades.

Lophiiformes is generally considered to be sister to Tetraodontiformes based on molecular analyses[2,3,51,58], a finding that also has morphological support[59]. This sister-group relationship was recovered by our all-genes IQ-TREE and most gene-subset trees (Fig. S3), but not our all-genes ASTRAL tree, for which Caproidae+Scatophagidae was sister to Lophiiformes (Fig. S4). Similarly, Rabosky et al.[23] found that Lophiiformes was sister to a clade formed by Cepolidae and Tetraodontiformes. It seems plausible that the interrelationships among Lophiiformes, Tetraodontiformes, Priacanthiformes, Caproiformes, and *incertae sedis* families like Siganidae are affected by (as-of-yet unidentified) systematic errors, yielding inconsistency among phylogenetic analyses.

### Suborder Lophioidei

Contains: Lophiidae

Lophiidae is typically found to be sister to all other lophiiforms[55] (Figs. S2–S7).

### Suborder Ogcocephaloidei

Contains: Ogcocephalidae

### Suborder Antennarioidei

Equal to: Antennariidae (Brownstein et al.[57])

Contains: Antennariidae, Rhycheridae, Tathicarpidae, Tetrabrachiidae, Brachionichthyidae, Lophichthyidae (NS), Histiophrynidae (NS)

The suborder Antennarioidei used to contain four nominal families: Antennariidae, Brachionichthyidae, Tetrabrachiidae, and Lophichthyidae. Lophichthyidae has not yet been included in any molecular phylogenetic analysis, but morphological analyses suggest it is sister to Tetrabrachiidae+Antennariidae[60]. The non-monophyly of Antennariidae was resolved by Hart et al. [61] using UCE's. These authors split Antennariidae into four monophyletic families:

Antennariidae (containing *Fowlerichthys, Antennarius, Antennatus, Abantennarius, Nudiantennarius, and Histrio*), Rhycheridae, Histiophrynidae, and Tathicarpidae. This arrangement has been supporting using exons (ref.[55] and this study) and so we follow their revised scheme. Note that other authors opted to synonymize Antennarioidei with Antennariidae instead (i.e. resolving the non-monophyly of Antennariidae by lumping instead of splitting)[57].

### Suborder Chaunacoidei

Contains: Chaunacidae

Chaunacidae is uncontroversially the sister group to Ceratioidei[55,57].

### Suborder Ceratioidei

Equal to: Ceratoidea (Ghez2022)

Contains: Oneirodidae, Melanocetidae, Himantolophidae, Diceratiidae, Linophrynidae, Gigantactinidae, Ceratiidae, Caulophrynidae, Centrophrynidae, Thaumatichthyidae, Neoceratiidae (NS)

That all eleven families listed here comprise a monophyletic group has been confirmed in other molecular[55,57] and morphological analyses[62]. The interrelationships within Ceratioidei are generally affected by issues such as incomplete lineage sorting[57], but some placements are highly stable in our analyses and others: specifically, Caulophrynidae as sister to all remaining ceratioids, and a clade containing Oneirodidae, Diceratiidae, Melanocetidae, and Himantolophidae (Figs. S2–S7).

### Order Tetraodontiformes

Equal to: Tetraodontoidei of Ghez2022/NT2024

Similar to Lophiiformes, we retain the traditional order-level Tetraodontiformes instead of the suborder-level Tetraodontoidei of Ghez2022. Assignments of tetraodontiform families to suborders has shifted over the past decade[1,2,5]. The interrelationships of Triacanthodoidei, Triacanthoidei, and *Triodon* (not sampled) are especially uncertain even across phylogenomic-scale analyses (Figs. S2–S4 and refs. [3,63]).

### Suborder Balistoidei

Contains: Balistidae, Monacanthidae

This sister-clade relationship is highly stable, appearing across all analyses (Figs. S2–S4).

### Suborder Ostracioidei

Contains: Ostraciidae, Aracanidae

This sister-clade relationship is highly stable, appearing across all analyses (Figs. S2–S4).

### Suborder Tetraodontoidei

**Changes from Bet2017:** Molidae is included in Tetraodontoidei to communicate its stable position as the sister to Diodontidae+Tetraodontidae. It was previously placed in its own suborder Moloidei (Bet2013/Bet2017). Placement of Molidae in Tetraodontoidei has precedence (e.g. refs.[5,22,35]).

Contains: Tetraodontidae, Diodontidae, Molidae

The clade (Molidae, (Diodontidae, Tetraodontidae)) is relatively stable, being recovered in most of our analyses except for some ASTRAL gene-subset trees (Figs. S2–S4) and in several past studies[3,23,63,64].

### Suborder Triacanthodoidei

Contains: Triacanthodidae

It remains unclear whether the Triacanthodidae (spikefishes) and Triacanthidae (triplespines) are reciprocally monophyletic (our coalescent analyses and refs.[2,3,23,63,64]) or not (our concatenation-based analyses). Wiley and Johnson[5] placed them in the shared suborder Triacanthoidei. Bet2017 found them to be sister families but placed them in their own suborders. This is the most stable arrangement given these uncertain interrelationships, and so we retain the two-suborder arrangement of Bet2013/Bet2017.

### Suborder Triacanthoidei

Contains: Triacanthidae

See comments under Triacanthodoidei.

### Suborder Triodontoidei

Contains: Triodontidae (NS)

### Order Caproiformes

Contains: Caproidae (including *Antigonia*)

We retain Caproiformes sensu Bet2017. Caproidae has been placed in its own order "Caproiformes" since Wiley and Johnson[5]. Caproidae is often found to be the sister to the clade containing Lophiiformes and Tetraodontiformes (e.g. Bet2017, Ghez2022). This position was also found in our all-genes IQ-TREE with bootstrap support=100 as well as most gene-subset IQ-TREES (Fig. S3). However, the placement of Caproidae was highly unstable across our ASTRAL trees (Fig. S4).

### Order Priacanthiformes

**Changes since Bet2017:** Scatophagidae is now included within the Priacanthiformes

Contains: Scatophagidae, Priacanthidae, Cepolidae

A clade containing (Scatophagidae, (Priacanthidae, Cepolidae)) was found by the all-genes IQ-TREE and all gene-subset IQ-TREES (Fig. S3). This set of relationships is identical to that found by Bet2017 (Fig. S7; although Scatophagidae was deigned "*incertae sedis*" in that study) and in Ghez2022 (Fig. S5). However, note that the position of Scatophagidae is unstable among the ASTRAL trees (Fig. S4). Rabosky et al. found a sister-group relationship between Scatophagidae and Priacanthidae[23].

### Order Chaetodontiformes (*sedis mutabilis*)

**Changes since Bet2017:** New circumscription of Chaetodontiformes

Contains: Chaetodontidae, Pomacanthidae (the placement of Leiognathidae is affected by systematic errors)

Bet2017 recognized Chaetodontiformes as containing Chaetodontidae and Leiognathidae (albeit with modest bootstrap support), a relationship also found by Ghez2022. In our analyses, Leiognathidae is a rogue taxon that never forms a relationship with Chaetodontidae, which may be caused by long branch attraction (see below). Pomacanthidae was considered "*incertae sedis*" by Bet2017.

A clade that includes Chaetodontidae and Pomacanthidae, although the exact circumscription varies, has precedence from morphology[4] and other molecular studies[3,45,58,65,66]. Our all-genes IQ-TREE resolves a sister-clade relationship between Chaetodontidae and Pomacanthidae with bootstrap support=100, which is also found in the majority of gene-subset IQ-TREE's (Fig. S3). The positions of both families are unstable across coalescent trees (Fig. S4). This group is a priority for future systematic study.

### Order Acanthuriformes

Equal to: Acanthuroidei (Ghez2022, NT2024)

Contains: Acanthuridae, Luvaridae, Zanclidae

The monophyly of these three families is supported by morphological evidence[67] and multi-locus molecular datasets[51]. The sister group relationship between Acanthuridae and Zanclidae, has persisted into the phylogenomic era without controversy[3,4]. We choose to retain the ordinal level status of this group following Bet2013/Bet2017 and originating with Wiley and Johnson[5].

### *Incertae sedis* within Lophiimorphae

Includes: Malacanthidae*, Callanthiidae*, Siganidae*, Lutjanidae, Haemulidae, Sciaenidae*, Emmelichthyidae*, Dinopercidae* (NS). Those marked with "*" were also considered *incertae sedis* in Bet2017. The positions of these families are unstable across our alternative analyses.

Ghez2022 found *Dinoperca* to be sister to *Haemulon* which suggests it can be minimally assigned to the superorder Lophiimorphae.

**Changes since Bet2017:** We do not find evidence for "Lutjaniformes" describing a sister-group relationship between Lutjanidae and Haemulidae.

### The new "new bush"

In 1989, Nelson[68] termed Perciformes the "bush at the top of the tree" in reference to the difficulty resolving relationships within this group. Twenty-eight years later, Bet2017 wrote: "although most family-level and ordinal groups within this series receive high nodal support, interrelationships among them are largely unresolved– Eupercaria constitutes the "new bush at the top."[2] In the phylogenomic era, this bush has been further resolved into five well-supported clades (considered orders by Ghez2022 and superorders herein). The "bush" has now been reduced to a set of taxa within Lophiimorphae, for which their interrelationships are highly unstable across molecular datasets (among our own gene subsets and comparing exons to UCEs[3]) and tree estimation strategies. This new "new bush" is denoted in Figs. S2–S5, and includes the orders Spariformes and Lobotiformes and the *incertae sedis* families Emmelichthyidae, Callanthiidae, Haemulidae, Lutjanidae, Malacanthidae, Sciaenidae, and Siganidae. We do not attempt to determine the sources of error creating the bush and leave this for future investigation. Interestingly, most nodes within the bush have a bootstrap support=100 in our all-genes concatenation-based tree, yet are resolved inconsistently among gene-subset trees (Fig. S3).

### Rogue taxa (*incertae sedis* at the superorder level)

The superorder-level placement of four families was inconsistent across our analyses. These were: Gerreidae, Leiognathidae, Creediidae and Champsodontidae (the latter two were usually sister families). We avoided assigning these families to a superorder pending future investigation.

Additionally, Monodactylidae (not sampled) as currently delimited[23,24] (*Monodactylus, Schuettea*) cannot be assigned to a superorder. Ghez2022 found *Schuettea* to be in their Acropomatiformes (our Pempherimorphae) and *Monodactylus* in their Acanthuriformes (our Lophiimorphae). Smith et al. [42] also considered *Schuettea* to be an acropomatiform.

### Gerreidae/Gerreiformes (Bet2017)

Gerreidae was sometimes the earliest diverging lineage within Lophiimorphae (Fig. S4; see also ref.[3]) and sometimes sister to Labrimorphae (Fig. S3; see also refs.[45,69,70]). Gerreidae has been assigned to Acanthuriformes (NT2024), but this assignment may be premature as its affiliation is highly unstable even with genome-wide data.

We continue to recognize the order Gerreiformes (containing Gerreidae only) following Bet2017. No matter its placement, it is apparent that Gerreidae is a phylogenetically unique lineage relative to other eupercarians.

### Leiognathidae

Leiognathidae has been previously assigned to Chaetodontiformes due to its presumed sister relationship with Chaetodontidae (Bet2017), a relationship also found by Ghez2022 using UCEs (see also ref. [71]). Our analyses did not find support for a close relationship between Leiognathidae and Chaetodontidae (see also ref.[45]). In our analyses, Leiognathidae was sometimes sister to Creediidae and Champsodontidae (Fig. S3), although this relationship is likely an artefact of long branch attraction, and sometimes is placed sister to Callanthiidae (Fig. S4).

### Creediidae and Champsodontidae

This clade was sometimes found in a clade with Siganidae, Leiognathidae, and Callanthiidae within Lophiimorphae (Fig. S3) and sometimes sister to Gerreidae and Lophiimorphae (Fig. S4). These families have been placed in Pempheriformes/Acropomatiformes (Bet2017, Ghez2022, NT2024) based on exons and UCE loci (see also ref.[42]). The families are subtended by very long branches in ref.[42], therefore it is likely that long branch attraction is causing the rogue behavior of these families.

**Data S2: Fossil calibrations for Eupercaria**

Museum abbreviations:

CMC = Museo della Famiglia Cerato, Bolca, Italy
CMM-V = Calvert Marine Museum Vertebrate Collection, Maryland, USA
GSP-UM = Geological Survey of Pakistan-University of Michigan collection, Quetta, Balochistan, Pakistan
IGM = Instituto Geológico de México, Universidad Nacional Autónoma de México, Ciudad de México, México
KIGAM = Korea Institute of Geoscience and Mineral Resources, Daejeon, South Korea
KMNH: Kitakyushu Museum of Natural History and Human History, Fukuoka, Japan
LACM = Los Angeles County Museum, California, USA
MCSNV/MCSN/MCSV = Museo Civico di Storia Naturale di Verona, Verona, Italy
MGPD/MGP-PD = Museo di Geologia e Paleontologia, Università di Padova, Padova, Italy
MGUP = Museo di Geologia e Paleontologia, University of Padua, Padua, Italy
MNHN = Muséum National d'Histoire Naturelle, Paris, France
NHMUK = Natural History Museum, London, United Kingdom
NHMW: Naturhistorisches Museum, Vienna, Austria
PIN = Paleontological Institute of the Russian Academy of Sciences, Moscow, Russia
SMNS = Staatliches Museum für Naturkunde, Stuttgart, Baden-Württemberg, Germany
UMMP = University of Michigan Museum of Paleontology, Michigan, USA
USNM = National Museum of Natural History, Smithsonian Institution, Washington, DC, USA
ZIN = Zoological Institute of the Russian Academy of Sciences, St. Petersburg, Russia
ZPALWr = Zoological Institute, Department of Paleozoology, Wroclaw University, Poland

**In total, we used 64 node calibrations:**

**** Calibrations 1-8 are fossils related to outgroups of Eupercaria ****
**** Calibrations 9-49 are fossils within Eupercaria ingroup ****
**** Calibrations 50-64 are based on the Isthmus of Panama ****

We used the Hedman algorithm[1] to estimate "soft" upper bounds on calibrations. This approach uses a list of fossil outgroup age records based on the oldest minima (below) to produce a probability distribution of the origin of a given clade. For the full dataset calibrated in BEAST v2.7.7[2] all calibration densities (except for the root; see below) were specified as lognormal distributions by setting the standard deviation to 1.0 for all priors and adjusting the mean in BEAUti to match the 95% credible interval estimated with the Hedman method for each calibration point.

The outgroup sequence to Acanthomorpha is based on refs. [3–5]

247.1 Ma, Holostei, †*Wasonulus eugnathoides*;
236.0 Ma, †*Prohalecites porroi*;

221.0 Ma, †Pholidophoridae, †*Knerichthys bronni*;
193.81 Ma, †*Dorsetichthys bechei*;
181.7 Ma, †*Leptolepis coryphaenoides*;
166.1 Ma, †Ichthyodectiformes, †*Occithrissops willsoni*;
151.2 Ma, Elopomorpha, †*Anaethalion zapporum*;
150.94 Ma, Otocephala, †*Tischlingerichthys viholi*;
150.94 Ma, non-eurypterygian Euteleostei †*Leptolepides haerteisi*;
125.0 Ma, Aulopiformes, †*Atolvorator longipectoralis*;
98.0 Ma, Lampridiformes, †*Aipichthys minor*

The Hedman method requires a maximum hard bound; for this bound we used †*Discoserra* (322.8 Ma) representing a probable stem neopterygian.

1. Root (Total Group Holocentridae)

   MRCA: *Myripristis berndti* & *Lobotes pacificus*

   Fossil: †*Stichocentrus liratus* NHMUK PV P.47835. Stem holocentrid[6]

   Notes: We follow Friedman et al. [5] and Hughes et al. [7] in using this fossil to calibrate the node representing the MRCA between Holocentridae and all remaining acanthomorphs. To avoid artifactual root age that BEAST estimates (i.e., excessively old ages), we applied a normal distribution to the trees estimated with the full dataset. We obtained the mean by averaging the ages obtained from independent, large-scale fish trees that applied multiple fossil percomorph calibrations, both within and outside Eupercaria. We therefore apply a mean of 133 Ma and a standard deviation of 5 for the root calibration, which encompasses the range of values reported in previous studies (Table S4) within three standard deviations of the mean.

   Min. age = 94.0 Ma, Hajula Lagerstätte, Lebanon[8]

   95% soft upper = 125 Ma

   Normal Distribution
   Mean = 133
   Standard Deviation = 5

   Outgroup age sequence = 247.1, 193.81, 193.81, 181.7, 164.9, 151.2, 150.89, 103.13, 98.0, 94.0

**Table S4**. Age estimates for the root (total group Holocentridae) from previous studies.

| Study | Mean age Root (Ma) | 95% HPD |
|---|---|---|
| Alfaro et al. (2018) [4] | 124 | (112-136) |
| Betancur-R et al. (2013) [9] | 141 | (125-158) |
| Betancur-R et al. (2017) [10] | 145 | |
| Chen et al. (2014) [11] | 125 | (114-136) |
| Ghezelayagh et al. (2022) [12] | 133 | |
| Hughes et al. (2018) [13] | 128 | (123-133) |
| Near et al. (2012) [14] | 126 | (118-136) |
| Near et al. (2013) [15] | 126 | (118-136) |
| Rabosky et al. (2018) [16] | 129 | |

2. Crown Syngnatharia

MRCA: *Aulostomus maculatus* & *Macroramphosus scolopax*

Fossil: †*Gasterorhamphosus zuppichinii* MCSNV Na T 877.

Orr [17] argued this is a stem lineage of a clade containing Macrorhamphosidae and Centriscidae (see also [14]).

Min. age = 83.6 Ma, "Calcari di Melissano", Porto Selvaggio, Lecce Province, Italy.

Notes: Previous studies[5] used a much more recent minimum age (~69 Ma). Santaquiteria et al. [18] reviewed new evidence that supports an older age of †*Gasterorhamphosus,* suggesting that past studies underestimated the age of Syngnatharia.

95% soft upper = 111 Ma

Lognormal Distribution
Mean = 1.64
Standard Deviation = 1.0

Outgroup age sequence = 247.1, 193.81, 193.81, 181.7, 164.9, 151.2, 150.89, 103.13, 98.0, 94.0, 83.6

3. Crown Scombridae

MRCA: *Rastrelliger brachysoma* & *Scomberomorus regalis*

Tree differences: This calibration was not used for the ASTRAL tree because Scombridae was not monophyletic.

Fossil: *Scomber †saadii*, MNHN EiP 14. Shares characteristics of *Scomber* and *Rastrelliger*[19]

Min. age = 33.9 Ma, Padbeh Formation of Istehbanât, Iran[5]

95% soft upper = 103 Ma

Lognormal Distribution
Mean = 2.59
Standard Deviation = 1.0

Outgroup age sequence = 247.1, 193.81, 193.81, 181.7, 164.9, 151.2, 150.89, 103.13, 98.0, 94.0, 33.9

4. Total group Stromateidae

MRCA: *Nomeus gronovii* & *Peprilus snyderi*

Fossil: †*Pinichthys pulcher*, PIN 1413/55. A stem stromateid[5]

Min. age = 32.02 Ma, Pshekha Horizon, †*Planorbella* Beds, Lower Maikop, Caucasus[5]

95% soft upper =103 Ma

Lognormal Distribution
Mean = 2.62
Standard Deviation = 1.0

Outgroup age sequence = 247.1, 193.81, 193.81, 181.7, 164.9, 151.2, 150.89, 103.13, 98.0, 94.0, 32.02

5. Total group Beloniformes

MRCA: *Hyporhamphus intermedius* & *Melanotaenia praecox*

Fossil: †*Rhamphexocoetus volans*, MGUP 8866, crown beloniform[15,20]

Notes: Given our limited outgroup sampling, the more conservative calibration position is Total group Beloniformes rather than Crown Beloniformes.

Min. age = 48.5 Ma. Bolca, Pesciara site, Italy[21]

95% soft upper = 105 Ma

Lognormal Distribution
Mean = 2.39
Standard Deviation = 1.0

Outgroup age sequence = 247.1, 193.81, 193.81, 181.7, 164.9, 151.2, 150.89, 103.13, 98.0, 94.0, 48.5

6. Total group Pomacentridae

MRCA: *Amphistichus rhodoterus* & *Acanthochromis polyacanthus*

Tree differences: The MRCA of *Calloplesiops altivelis* and *Acanthochromis polyacanthus* was used for the ASTRAL tree.

Fossil: †*Chaychanus gonzalezorum*, IGM 11383, placement in the family Pomacentridae is supported by the combination of the presence of pharyngognathy and two anal-fin spines in a supernumerary association[22]. This is the oldest fossil record of Pomacentridae.

Min. age = 61.5, Belisario Domínguez quarry, Mexico[22]

95% soft upper = 107 Ma

Mean = 2.18
Standard Deviation = 1.0

Outgroup age sequence = 247.1, 193.81, 193.81, 181.7, 164.9, 151.2, 150.89, 103.13, 98.0, 94.0, 61.5

7. Total group Channidae

MRCA: *Channa argus* & *Anabas testudineus*

Fossil: †*Eochanna chorlakkiensis,* GSP-UM 781, stem channid[23]. Fossil shares characteristics of extant Channidae and Synbranchidae (series Anabantaria sensu Betancur et al. [24]).

Min. age = 41.3 Ma, Kuldana Formation, Pakistan[25]

95% soft upper = 104 Ma

Mean = 2.5
Standard deviation = 1.0

Outgroup age sequence = 247.1, 193.81, 193.81, 181.7, 164.9, 151.2, 150.89, 103.13, 98.0, 94.0, 41.3

8. Total group Carangidae

MRCA: *Caranx ignobilis* & *Rachycentron canadum*

Fossil: †*Archaeus oblongus*, PIN 2179/94, a stem carangid[26,27]

Min. age = 55.3 Ma, Danatinsk Formation, Kopet Dag, Turkmenistan[28–30]

95% soft upper = 106 Ma

Mean = 2.28
Standard deviation = 1.0

Outgroup age sequence = 247.1, 193.81, 193.81, 181.7, 164.9, 151.2, 150.89, 103.13, 98.0, 94.0, 55.3

**Ingroup fossil calibrations**

**Perciformes:**

9. Total group Eleginopidae

MRCA: *Artedidraco skottsbergi* & *Eleginops maclovinus*

Fossil: †*Proeleginops grandeastmanorum*, USNM 433396. Balushkin [31] argued that this fossil was a stem eleginopsid based on the following combination of shared derived characters: V-shaped crests on the frontals, orbito-rostral part of the skull longer than the posterior part, supratemporal fossa well defined, seismosensory canals on the upper part of head closed, and similarity in shape of the mucus pit in the region of the coronal commissure, in the arrangement of large bony indentations in the posterior part of the frontals, and in the structure of the opercle. This identification has been accepted by subsequent authors[32,33].

Min. age = 38 Ma, Locality RV 8200 in Telm5, La Meseta formation, Seymour Island, Antarctica [34]

Notes: This is the oldest known fossil notothenoid[31,32].

95% soft upper = 104 Ma

Mean = 2.54
Standard deviation = 1.0

Outgroup age sequence = 247.1, 193.81, 193.81, 181.7, 164.9, 151.2, 150.89, 103.13, 98.0, 94.0, 38

10. Crown Trachinidae

MRCA: *Trachinus draco* & *Trachinus pellegrini*

Fossil: *Trachinus* cf. *†minutus* (SMNS 87457/293). This is a common species in Rupelian deposits of Europe[35,36]

Notes: This is the first time to our knowledge that this fossil is used to time-calibrate a molecular phylogeny of fishes.

Min. age: 31.2, Rauenberg Lagerstatte, Baden-Wurttemberg, Germany[37]

95% soft upper = 103 Ma

Mean = 2.63
Standard deviation = 1.0

Outgroup age sequence = 247.1, 193.81, 193.81, 181.7, 164.9, 151.2, 150.89, 103.13, 98.0, 94.0, 31.2

11. Crown Scorpaenoidei sensu Smith et al. [38]

MRCA: *Platycephalus indicus* & *Prionotus rubio*

Fossil: Scorpaenoidei family, genus, and species indet., MCSNV IG.VR.66957 [39]

First evidence of scorpaenoid fishes in the Bolca Lagerstätte and one of the oldest scorpaenoid fossils known. The cranium has a parietal spine overlaying a lateral-line canal, a feature diagnostic of the Scorpaenoidei sensu Smith et al. [38]. The authors presumed the fossil had affinities to the families Scorpaenidae or Synanceiidae, which would place the fossil in a nested position within crown Scorpaenoidei.

Notes: This is the first time to our knowledge that this fossil is used to time-calibrate a molecular phylogeny of fishes. Smith et al. [38] recognized Scorpaenoidei as containing the families Bembridae, Hoplichthyidae, Neosebastidae, Platycephalidae, Plectrogeniidae, Scorpaenidae, Synanceiidae, and Triglidae. Since their Scorpaenoidei is not monophyletic in our analyses (see Data S1), we conservatively chose the node representing the MRCA of those families.

Min. age = 48.96 Ma, Bolca, Monte Postale site, Italy[21]

95% soft upper = 105 Ma

Mean = 2.38
Standard deviation = 1.0

Outgroup age sequence = 247.1, 193.81, 193.81, 181.7, 164.9, 151.2, 150.89, 103.13, 98.0, 94.0, 48.96

12. Total group *Setarches/*Setarchinae

MRCA: *Setarches longimanus* & *Neomerinthe beanorum*

Fossil: *Setarches sp.,* ZIN 551p/552p[40]. Identification confirmed in-person by G. Carnevale.

Notes: This is the earliest record of Setarchinae to our knowledge, supplanting another fossil from the late Miocene[41]. This is the first time that this fossil is used to time-calibrate a molecular phylogeny of fishes. Note that the genus *Setarches* is not monophyletic in our analyses, so this fossil placement is conservative.

Min. age: 13.1 Ma, Kurasi Formation, Sakhalin, Russia[40,42]

95% soft upper = 83.5 Ma

Mean = 2.61
Standard deviation = 1.0

Outgroup age sequence = 247.1, 193.81, 193.81, 181.7, 164.9, 151.2, 150.89, 103.13, 98.0, 94.0, 48.96, 13.1

13. Total group *Zaprora*

MRCA: *Zaprora silenus* & *Pholis ornata*

Fossil: *Zaprora †koreana*, KIGAM 9A168, the earliest known fossil Zaproridae[43]. The authors placed this fossil in the family Zaproridae based on the following combination of shared derived characters: comparatively short-based anal fin with three anterior spines, well-pronounced caudal peduncle, very high neural and haemal spines, number of caudal vertebrae and anal fin rays, and deep caudal region with high, entirely spinous dorsal fin.

Notes: This is the first time to our knowledge that this fossil is used to time-calibrate a molecular phylogeny of fishes.

Min. age = 15 Ma, Duho Formation, Pohang, South Korea, representing an environment deeper than 500 meters[44]

95% soft upper = 84.0 Ma

Mean = 2.59
Standard deviation = 1.0

Outgroup age sequence = 247.1, 193.81, 193.81, 181.7, 164.9, 151.2, 150.89, 103.13, 98.0, 94.0, 48.96, 15

14. Total group *Myoxocephalus*

MRCA: *Psychrolutes phrictus* & *Myoxocephalus scorpius*

Fossil: *Myoxocephalus sp.*, ZIN 550p[40]. Identification confirmed in-person by G. Carnevale.

Notes: This is the first time that this fossil is used to time-calibrate a molecular phylogeny of fishes.

Min. age: 13.1 Ma, Kurasi Formation, Sakhalin, Russia[40]

95% soft upper = 83.5 Ma

Mean = 2.61
Standard deviation = 1.0

Outgroup age sequence = 247.1, 193.81, 193.81, 181.7, 164.9, 151.2, 150.89, 103.13, 98.0, 94.0, 48.96, 13.1

15. Total group Pholidae

MRCA: *Pholis ornata* & *Ptilichthys goodei*

Fossil: *†Agnevichthys gretchinae* (ZIN 150p) and *†Paleopholis laevis* (ZIN 198p), extinct genera within Pholidae[45]. Fossils are placed in the family Pholidae based on morphological synapomorphies including a low dorsal fin composed of thick spines, amphicoelus vertebrae, and the first pterygiophore of the anal fin significantly displaced anteriorly with several haemal spines inserted between the first and second pterygiophore. The fossils lack several autapomorphies of living Pholidae, justifying the stem placement of this calibration.

Min. age: 11.63 Ma (Serravallian–Tortonian), Agnevo Formation, Sakhalin, Russia[46]

95% soft upper = 60.5 Ma

Mean = 2.245
Standard deviation = 1.0

Outgroup age sequence = 247.1, 193.81, 193.81, 181.7, 164.9, 151.2, 150.89, 103.13, 98.0, 94.0, 48.96, 15, 11.63

16. Total group *Gasterosteus*

MRCA: *Gasterosteus aculeatus* & *Pungitius pungitius*

Fossil: *Gasterosteus cf. wheatlandi,* LACM 150177[20,47]. Species identification is debated, so we follow Hughes et al. [7] by treating this fossil conservatively as a stem *Gasterosteus*.

Min. age = 13.1 Ma, Alta Mira Shale, Monterey Formation, California[20]

95% soft upper = 83.5 Ma

Mean = 2.61
Standard deviation = 1.0

Outgroup age sequence = 247.1, 193.81, 193.81, 181.7, 164.9, 151.2, 150.89, 103.13, 98.0, 94.0, 48.96, 13.1

17. Crown *Parapercis* (Pinguipedidae)

MRCA: *Parapercis atlantica* & *Parapercis pacifica*

Fossil: *Parapercis †mesogea,* MNHN ORA26. Fossils assigned to *Parapercis* based on their overall physiognomy, meristics and a single strong opercular spine[48].

Min. age: 6 Ma, Raz-el-Ain, Oran, Algeria[49]

95% soft upper = 101 Ma

Mean = 2.01
Standard deviation = 1.0

Outgroup age sequence = 247.1, 193.81, 193.81, 181.7, 164.9, 151.2, 150.89, 103.13, 98.0, 94.0, 6

**Labriformes:**

18. Total group Hypsigenyini (Crown Labridae)

MRCA: *Coris pictoides* & *Lachnolaimus maximus*

Fossil: †*Phyllopharyngodon longipinnis*, MCSNV T 149 [50]. Fossil is the earliest known member of Hypsigenyini, which is the first tribe to split within Labridae and sister to all other tribes in our analyses and others[7]

Min. age = 48.5 Ma, Bolca, Pesciara site, Italy[21]

95% soft upper = 105 Ma

Mean = 2.39
Standard deviation = 1.0

Outgroup age sequence = 247.1, 193.81, 193.81, 181.7, 164.9, 151.2, 150.89, 103.13, 98.0, 94.0, 48.5

19. *Bodianus* & *Achoerodus*

MRCA: *Bodianus rufus* & *Achoerodus gouldii*

Fossil: *†Trigonodon jugleri,* NHMW 1997z0178/1970. The earliest record of Pseudodacinae[51]. Fossil is morphologically similar to the living genus *Pseudodax.*

Notes: We did not sample *Pseudodax,* so we placed this calibration at the MRCA of *Achoerodus* and *Bodianus* consistent with its placement in Hughes et al. [7].

Min. age = 15.97 Ma, Prambachkirchen and Aussertreffling, Austria[52]

95% soft upper = 83.8 Ma

Mean = 2.57
Standard deviation = 1.0

Outgroup age sequence = 247.1, 193.81, 193.81, 181.7, 164.9, 151.2, 150.89, 103.13, 98.0, 94.0, 48.5, 15.97

20. Crown Labrini

MRCA: *Symphodus melops* & *Sparisoma viride*

Fossil: *Tautoga sp.,* CMM-V-327. *Tautoga* is a living genus.

Notes: Since *Tautoga* was not sampled in our study, we follow Hughes et al. [7] by conservatively placing this calibration at crown Labrini.

Min. age = 15 Ma, Langhian, Calvert Formation, Maryland, USA[53]

95% soft upper = 83.7 Ma

Mean = 2.58
Standard deviation = 1.0

Outgroup age sequence = 247.1, 193.81, 193.81, 181.7, 164.9, 151.2, 150.89, 103.13, 98.0, 94.0, 48.5, 15

21. Total group *Symphodus*

MRCA: *Symphodus melops* & *Labrus bergylta*

Fossil: *Symphodus †westneati*, NHMW 2000z0135/0060a. Assigned to the extant genus *Symphodus* [54]

Min. age = 13.5 Ma, St. Margarethen, Burgenland, Austria[54]

95% soft upper = 60.9 Ma

Mean = 2.21
Standard deviation = 1.0

Outgroup age sequence = 247.1, 193.81, 193.81, 181.7, 164.9, 151.2, 150.89, 103.13, 98.0, 94.0, 48.5, 15, 13.5

22. Crown Julidini

MRCA: *Coris auricularis* & *Halichoeres poeyi*

Fossil: *Coris †sigismundi*, NHMW 1975/1691/0015 [54]

Notes: The living genus *Coris* is not monophyletic, so we follow Hughes et al. [7] in assigning this calibration to the earliest julidine node.

Min. age = 13.5 Ma, St. Margarethen, Burgenland, Austria [54]

95% soft upper = 83.5 Ma

Mean = 2.6
Standard deviation = 1.0

Outgroup age sequence = 247.1, 193.81, 193.81, 181.7, 164.9, 151.2, 150.89, 103.13, 98.0, 94.0, 48.5, 13.5

**Centrarchiformes:**

23. Total group *Coreoperca* (Crown Sinipercidae)

MRCA: *Coreoperca whiteheadi* & *Siniperca scherzeri*

Fossil: *Coreoperca †maruoi*, KMNH VP 100,261 [55]. Fossil was placed in living genus *Coreoperca* based on characteristics of the dorsal-fin spines, preopercle, neural spines, and scales.

Min. age = 15.3 Ma, Chojabaru Formation, Japan [55]

95% soft upper = 101Ma

Mean = 2.81
Standard deviation = 1.0

Outgroup age sequence = 247.1, 193.81, 193.81, 181.7, 164.9, 151.2, 150.89, 103.13, 98.0, 94.0, 15.3

24. *Pomoxis* & *Lepomis*

MRCA: *Pomoxis annularis* & *Lepomis humilis*

Fossil: †*Plioplarchus whitei,* USNM V 4695, extinct genus within crown Centrarchidae and one of the oldest centrarchid fossils [56]

Notes: Near and Kim [56] performed phylogenetic analyses and placed this fossil in the clade containing *Acantharchus*, *Ambloplites*, *Archoplites*, *Centrarchus*, and *Pomoxis*. This clade is sister to a polytomy containing *Enneacanthus* (not sampled in our analyses) and *Lepomis*.

Min. age = 34.2 Ma, Sentinel Butte, North Dakota, USA [57,58]

95% soft upper = 103Ma

Mean = 2.54
Standard deviation = 1.0

Outgroup age sequence = 247.1, 193.81, 193.81, 181.7, 164.9, 151.2, 150.89, 103.13, 98.0, 94.0, 34.2

25. *Ambloplites* & *Pomoxis*

MRCA: *Ambloplites rupestris* & *Pomoxis annularis*

Fossil: *Archoplites †clarki,* UMMP V 74202 (Smith and Miller 1985). Phylogenetic analyses by Near and Kim [56] placed *Archoplites* (living and extinct) as sister to *Ambloplites.*

Min. age = 15.5 Ma, Clarkia Lake Beds, Idaho, USA [59,60]

95% soft upper = 81.3 Ma

Mean = 2.54
Standard deviation = 1.0

Outgroup age sequence = 247.1, 193.81, 193.81, 181.7, 164.9, 151.2, 150.89, 103.13, 98.0, 94.0, 34.2, 15.5

**Pempheriformes:**

26. Crown *Acropoma*

MRCA: *Acropoma japonicum* & *Doederleinia berycoides*

Fossil: *Acropoma †lepidotus* MNHN Bol 415/416. Bannikov [61] re-identified this fossil as *Acropoma* based on (e.g.) the predorsal and fin formulas, small mouth, large and blunt teeth, and three anal fin spines.

Min. age = 48.5, Bolca, Pesciaria site, Italy [21]

95% soft upper = 105 Ma

Mean = 2.39

Standard deviation = 1.0

Outgroup age sequence = 247.1, 193.81, 193.81, 181.7, 164.9, 151.2, 150.89, 103.13, 98.0, 94.0, 48.5

27. Total group Champsodontidae

MRCA: *Champsodon atridorsalis* and *Crystallodytes cookie*

Fossil: †*Eochampsodon elongatus* PIN 4425–46 [62]. A stem member of Champsodontidae; differs from living genera by its vertebral counts, more anterior position of the rostral extremity of the anal fin, and reduced fimbriation of the opercular series.

Notes: This is the earliest fossil of Champsodontidae.

Min. age = 38.4 Ma, Gornyi Luch, Kuma Horizon, Caucasus, Russia [63]

95% soft upper = 104 Ma

Mean = 2.54
Standard deviation = 1.0

Outgroup age sequence = 247.1, 193.81, 193.81, 181.7, 164.9, 151.2, 150.89, 103.13, 98.0, 94.0, 38.4

**Lophiimorphae:**

28. Total group Ephippidae

MRCA: *Drepane punctata* & *Platax teira*

Fossil: †*Eoplatax papilio,* MGP-PD 26285, an ephippid with unclear phylogenetic relationships to living genera[64]

Min. age = 48.5 Ma, Bolca, Pesciara site, Italy [21]

95% soft upper = 105 Ma

Mean = 2.39
Standard deviation = 1.0

Outgroup age sequence = 247.1, 193.81, 193.81, 181.7, 164.9, 151.2, 150.89, 103.13, 98.0, 94.0, 48.5

29. Crown Lutjanidae

MRCA: *Etelis radiosus* & *Lutjanus purpureus*

Fossil: †*Hypsocephalus atlanticus,* LACM VP 27859, presumably a stem *Hoplopagrus* [65]

Notes: We follow Rincon-Sandoval et al. [66] who used this fossil to calibrate crown Lutjanidae, the most conservative position given that the fossil shares characters with several living genera of lutjanids.

Min. age = 33.9 Ma. *Operculinoides-Asterocyclina* Zone in the Crystal River Formation in north Florida [67]

95% soft upper = 103 Ma

Mean = 2.59
Standard deviation = 1.0

Outgroup age sequence = 247.1, 193.81, 193.81, 181.7, 164.9, 151.2, 150.89, 103.13, 98.0, 94.0, 33.9

30. Total group *Dentex* (Crown Sparidae)

MRCA: *Dentex carpenteri* & *Acanthopagrus latus*

Fossil: †*Sciaenurus bowerbanki*, NHMUK P3975.

Notes: Phylogenetic analyses by Day et al. [68] place this fossil as sister to *Dentex.* Since *Dentex* is polyphyletic in our analyses, our chosen placement includes the stem of the MRCA of all living *Dentex* species.

Min. age = 52.1 Ma, London Clay Formation, Great Britain [69]

95% soft upper = 105 Ma

Mean = 2.32
Standard deviation = 1.0

Outgroup age sequence = 247.1, 193.81, 193.81, 181.7, 164.9, 151.2, 150.89, 103.13, 98.0, 94.0, 52.1

31. Crown Malacanthidae

MRCA: *Branchiostegus australiensis* & *Malacanthus plumieri*

Fossil: †*Hoplolatilus visendus,* PIN RAN 4425–30 [70]. Placed in the living genus *Hoplolatilus* based on the a lack of a strongly expressed skull bone spination, vertebral formula, and meristic characters.

Min. age = 38.4 Ma, Gornyi Luch, Kuma Horizon, Caucasus, Russia [63]

Notes: This is the earliest plausible malacanthid fossil[70]. While we did not sample *Hoplolatilus* in our analyses, the genus is putatively well nested in the phylogeny of Malacanthidae, justifying the crown placement of the fossil calibration. *Hoplolatilus* and *Malacanthus* are in the subfamily Malacanthinae, which is presumably sister to the subfamily Latilinae on the basis of morphological systematics[71].

95% soft upper = 104 Ma

Mean = 2.54
Standard deviation = 1.0

Outgroup age sequence = 247.1, 193.81, 193.81, 181.7, 164.9, 151.2, 150.89, 103.13, 98.0, 94.0, 38.4

32. Total group *Chaetodon*

MRCA: *Chaetodon capistratus* & *Prognathodes aculeatus*

Fossil: *†Chaetodon wattsi*, MGP-PD R664. This species does not belong to any living subgenus of *Chaetodon* [72].

Min. age = 28 Ma, Calcareniti di Castelgomberto Formation, Perarolo, Italy [72]

95% soft upper = 103 Ma

Mean = 2.67
Standard deviation = 1.0

Outgroup age sequence = 247.1, 193.81, 193.81, 181.7, 164.9, 151.2, 150.89, 103.13, 98.0, 94.0, 28

33. Total group Luvaridae (Crown Acanthuriformes sensu Betancur et al. [24])

MRCA: *Luvarus imperialis* & *Acanthurus lineatus*

Fossil: †*Kushlukia permira,* PIN 2179/64, †Kushlukiidae. Phylogenetic analyses by Bannikov and Tyler [73] placed Kushlukiidae as sister to Luvaridae.

Min. age = 55.3 Ma, Danatinsk Formation, Kopet Dag, Turkmenistan[28–30]

95% soft upper = 106

Mean = 2.26
Standard deviation = 1.0

Outgroup age sequence = 247.1, 193.81, 193.81, 181.7, 164.9, 151.2, 150.89, 103.13, 98.0, 94.0, 55.3

34. Total group Nasinae (Crown Acanthuridae)

MRCA: *Naso minor* & *Prionurus laticlavius*

Fossils: †*Sorbinithurus sorbinii* MCSNV 524, stem nasine[74]

Min. age = 48.5 Ma, Bolca, Pesciara site, Italy [21]

95% soft upper = 89.4 Ma

Mean = 2.07
Standard deviation = 1.0

Outgroup age sequence = 247.1, 193.81, 193.81, 181.7, 164.9, 151.2, 150.89, 103.13, 98.0, 94.0, 55.3, 48.5

35. Total group *Pristigenys* (Crown Priacanthidae)

MRCA: *Pristigenys niphonia* & *Priacanthus arenatus*

Fossil: †*Pristigenys substriata,* MNHN F.Bol529. Phylogenetic analyses by Carnevale et al. [75] placed *Pristigenys* as sister to *Pseudopriacanthus.* This clade was sister to a clade containing all other living priacanthid genera (*Cookeolus, Heteropriacanthus, Priacanthus*).

Min. age = 48.5 Ma, Bolca, Pesciara site, Italy [21]

95% soft upper = 105 Ma

Mean = 2.39
Standard deviation = 1.0

Outgroup age sequence = 247.1, 193.81, 193.81, 181.7, 164.9, 151.2, 150.89, 103.13, 98.0, 94.0, 48.5

36. ***** For all trees, we applied two alternative calibration schemes: with and without #36.**

Total group Tetraodontiformes

MRCA: *Mola mola* & *Antennarius commerson*

Fossil: †*Plectocretacicus clarae,* MCSV S.L.1/2

Notes: According to phylogenetic analyses by Arcila and Tyler [76], the extinct †Plectocretacicoidea is either sister to Lophiiformes+Tetraodontiformes or sister to Tetraodontiformes. Their position as stem Tetraodontiformes is supported by 14 synapomorphies. However, this position has been questioned by other authors, who instead believe it to be *incertae sedis* in Acanthomorpha[4]. This fossil potentially has large consequences on the overall age of Lophiimorphae/Eupercaria, given its age and relationship to its sister group Tetraodontiformes [77].

Min. age = 94 Ma, Sannine Limestone, Hakel, Lebanon [77–79]

95% soft upper = 113 Ma

Mean = 1.28
Standard deviation = 1.0

Outgroup age sequence = 247.1, 193.81, 193.81, 181.7, 164.9, 151.2, 150.89, 103.13, 98.0, 94.0, 94.0

37. Total group Diodontidae+Tetraodontidae (Tetraodontoidei sensu [76])

MRCA: *Mola mola* & *Arothron hispidus*

Fossil: †*Balkaria histiopterygia*, PIN 5314/2. †Balkariidae

Notes: Phylogenetic analyses by Arcila and Tyler [76] placed this fossil as sister along the stem of Diodontidae. However, Bannikov et al. [80] suggested it is sister to (Diodontidae, Tetraodontidae). We follow Hughes et al. [7] by using the more conservative placement suggested by Bannikov et al. [80].

Min. age = 55.8 Ma, Kheu River Formation, Kabardino-Balkaria, Russia [76,80]

With †*Plectocretacicus:*

95% soft upper = 98.3 Ma

Mean = 2.1
Standard deviation = 1.0

Outgroup age sequence = 247.1, 193.81, 193.81, 181.7, 164.9, 151.2, 150.89, 103.13, 98.0, 94.0, 94.0, 55.8

Without †*Plectocretacicus:*

95% soft upper = 106 Ma

Mean = 2.25
Standard deviation = 1.0

Outgroup age sequence = 247.1, 193.81, 193.81, 181.7, 164.9, 151.2, 150.89, 103.13, 98.0, 94.0, 55.8

38. Total group Tetraodontidae

MRCA: *Chilomycterus antennatus* & *Arothron hispidus*

Fossil: †*Eotetraodon tavernei*, MCSNV IGVR 81994. Phylogenetic analyses by Arcila and Tyler [76] placed this fossil along the stem of Tetraodontidae.

Min. age = 48.5 Ma, Bolca, Pesciara site, Italy [21]

With †*Plectocretacicus:*

95% soft upper = 85.8 Ma

Mean = 1.97
Standard deviation 1.0

Outgroup age sequence = 247.1, 193.81, 193.81, 181.7, 164.9, 151.2, 150.89, 103.13, 98.0, 94.0, 94.0, 55.8, 48.5

Without †*Plectocretacicus:*

95% soft upper = 89.4 Ma

Mean = 2.04
Standard deviation = 1.0

Outgroup age sequence = 247.1, 193.81, 193.81, 181.7, 164.9, 151.2, 150.89, 103.13, 98.0, 94.0, 55.8, 48.5

39. Total group Balistidae

MRCA: *Odonus niger* & *Aluterus heudelotii*

Fossil: †*Gornylistes prodigiosus*, PIN 4425/95

Notes: This is the earliest record of Balistidae [81]. Phylogenetic analyses by Arcila and Tyler [76] confirmed this is a stem balistid.

Min age = 38.4 Ma, Gornyi Luch, Kuma Horizon, Caucasus, Russia [63]

With †*Plectocretacicus:*

95% soft upper = 96.8 Ma

Mean = 2.42
Standard deviation = 1.0

Outgroup age sequence = 247.1, 193.81, 193.81, 181.7, 164.9, 151.2, 150.89, 103.13, 98.0, 94.0, 94.0, 38.4

Without †*Plectocretacicus:*

95% soft upper = 104 Ma

Mean = 2.54
Standard deviation = 1.0

Outgroup age sequence = 247.1, 193.81, 193.81, 181.7, 164.9, 151.2, 150.89, 103.13, 98.0, 94.0, 38.4

40. Crown Ostraciidae

MRCA: *Lactophrys triqueter* & *Ostracion cubicus*

Fossil: †*Eolactoria sorbinii*, MCSN T6–T7. Phylogenetic analyses by Arcila et al. [79] and Arcila and Tyler [76] placed †*Eolactoria* as sister to *Ostracion.* The (*†Eolactoria,Ostracion*) clade is sister to (*†Oligolactoria,*(Lactoria,Tetrosomus)). This is presumably the earliest record of Ostraciidae.

Min. age = 48.5 Ma, Bolca, Pesciara site, Italy [21]

With †*Plectocretacicus:*

95% soft upper = 97.6 Ma

Mean = 2.25
Standard deviation = 1.0

Outgroup age sequence = 247.1, 193.81, 193.81, 181.7, 164.9, 151.2, 150.89, 103.13, 98.0, 94.0, 94.0, 48.5

Without †*Plectocretacicus:*

95% soft upper = 105 Ma

Mean = 2.39
Standard deviation = 1.0

Outgroup age sequence = 247.1, 193.81, 193.81, 181.7, 164.9, 151.2, 150.89, 103.13, 98.0, 94.0, 48.5

41. Total group *Triacanthodes*

MRCA*: Paratriacanthodes retrospinis* & *Triacanthodes_sp1*

Fossil: †*Carpathospinosus propheticus,* ZPALWr.A/3000–3009 [82].

Phylogenetic analyses by Arcila et al. [79] placed this fossil as sister to *Triacanthodes,* while Arcila and Tyler [76] found it was nested within *Triacanthodes.* We use the more conservative placement for the fossil here.

Min. age = 28 Ma, IPM 4 of Menilite Beds, Przysietnica, Poland (discussion in Tyler et al. [82], age used by refs. [76,79])

With †*Plectocretacicus:*

95% soft upper = 95.9 Ma

Mean = 2.57
Standard deviation = 1.0

Outgroup age sequence = 247.1, 193.81, 193.81, 181.7, 164.9, 151.2, 150.89, 103.13, 98.0, 94.0, 94.0, 28

Without †*Plectocretacicus:*

95% soft upper = 102 Ma

Mean = 2.66
Standard deviation = 1.0

Outgroup age sequence = 247.1, 193.81, 193.81, 181.7, 164.9, 151.2, 150.89, 103.13, 98.0, 94.0, 28

42. Total group Lophiidae (Crown Lophiiformes)

MRCA: *Sladenia sp.* and *Antennatus coccineus*

Fossil: †*Sharfia mirabilis*, MNHN Bol 38-39 [83]

Notes: Diagnostic characters unambiguously place †*Sharfia* in Lophiidae. It is distinct from all other lophiids by its triangular opercle and non-fimbriate subopercle (a state also found in some frogfishes, batfishes, and chaunacids). Phylogenetic analyses by Carnevale and Pietsch [84] using a matrix of 38 characters show that †*Sharfia* is sister to all other known lophiids based on one synapomorphy (opercle strongly bifurcate). †*Caruso* and †*Sharfia* are together the oldest known fossil representatives of Lophiidae. Since Lophiidae is sister to all other Lophiiformes, this fossil is effectively a calibration on Crown Lophiiformes.

Min. age = 48.5 Ma, Bolca, Pesciara site, Italy [21]

With †*Plectocretacicus:*

95% soft upper = 97.6 Ma

Mean = 2.25
Standard deviation = 1.0

Outgroup age sequence = 247.1, 193.81, 193.81, 181.7, 164.9, 151.2, 150.89, 103.13, 98.0, 94.0, 94.0, 48.5

Without †*Plectocretacicus:*

95% soft upper = 105 Ma

Mean = 2.39
Standard deviation = 1.0

Outgroup age sequence = 247.1, 193.81, 193.81, 181.7, 164.9, 151.2, 150.89, 103.13, 98.0, 94.0, 48.5

43. Total group *Sladenia* (Crown Lophiidae)

MRCA: *Sladenia sp.* and *Lophiodes spilurus*

Fossil: †*Caruso brachysomus* (articulated skeleton) MNHN Bol 42/43 [84]

Notes: Phylogenetic analyses by Carnevale and Pietsch [84] using a matrix of 38 characters show that the fossil †*Caruso* is sister to extant *Sladenia* based on two synapomorphies. This places it within Crown Lophiidae. †*Caruso* and †*Sharfia* are together the oldest known fossil representatives of Lophiidae.

Min. age = 48.5 Ma, Bolca, Pesciara site, Italy [21]

With †*Plectocretacicus:*

95% soft upper = 84.7 Ma

Mean = 1.94
Standard deviation = 1.0

Outgroup age sequence = 247.1, 193.81, 193.81, 181.7, 164.9, 151.2, 150.89, 103.13, 98.0, 94.0, 94.0, 48.5, 48.5

Without †*Plectocretacicus:*

95% soft upper = 88.6 Ma

Mean = 2.05
Standard deviation = 1.0

Outgroup age sequence = 247.1, 193.81, 193.81, 181.7, 164.9, 151.2, 150.89, 103.13, 98.0, 94.0, 48.5, 48.5

44. *Lophius* & *Lophiodes*

MRCA: *Lophius litulon* & *Lophiodes spilurus*

Fossil: †*Eosladenia caucasica*, PIN 4425–72 [85].

Notes: Phylogenetic analyses by Carnevale and Pietsch [84] using a matrix of 38 characters show that the clade (†*Eosladenia*,(*Lophiomus*,*Lophius*)) is sister to *Lophiodes.* The sister group relationship between †*Eosladenia* and *Lophiomus* + *Lophius* is supported by one synapomorphy (caudal centrum depressed, bearing lateral transverse processes). This places the fossil well within crown Lophiidae.

Min. age = 38.4 Ma [63], Bartonian Stage, Gornyi Luch, Kuma Horizon, Caucasus, Russia [85]

With †*Plectocretacicus:*

95% soft upper = 69.4 Ma

Mean = 1.79
Standard deviation = 1.0

Outgroup age sequence = 247.1, 193.81, 193.81, 181.7, 164.9, 151.2, 150.89, 103.13, 98.0, 94.0, 94.0, 48.5, 48.5, 38.4

Without †*Plectocretacicus:*

95% soft upper = 71.7 Ma

Mean = 1.85
Standard deviation = 1.0

Outgroup age sequence = 247.1, 193.81, 193.81, 181.7, 164.9, 151.2, 150.89, 103.13, 98.0, 94.0, 48.5, 48.5, 38.4

45. Total group Ogcocephalidae

MRCA: *Antennarius commerson* & *Coelophrys micropa*

Fossil: †*Tarkus squirei*, MCSNV T158/T159 [86]. Taxon is based on five skeletons including one complete articulated skeleton.

Notes: The fossil is diagnosed as an ogcocephalid based on the shape of the illicium, tubercles on the head and body, depressed body shape, horizontal gape of the mouth, and several skeletal and meristic characters [86]. Its relationship to living ogcocephalid genera is unclear, and is morphologically unique from all other ogcocephalids. This fossil is the earliest known record, as well as the first articulated skeletal record of Ogcocephalidae. Placing this calibration at the node corresponding to stem Ogcocephalidae is conservative.

Min. age = 48.5 Ma, Bolca, Pesciara site, Italy [21]

With †*Plectocretacicus:*

95% soft upper = 84.7 Ma

Mean = 1.95
Standard deviation = 1.0

Outgroup age sequence = 247.1, 193.81, 193.81, 181.7, 164.9, 151.2, 150.89, 103.13, 98.0, 94.0, 94.0, 48.5, 48.5

Without †*Plectocretacicus:*

95% soft upper = 88.6 Ma

Mean = 2.05
Standard deviation = 1.0

Outgroup age sequence = 247.1, 193.81, 193.81, 181.7, 164.9, 151.2, 150.89, 103.13, 98.0, 94.0, 48.5, 48.5

46. Total group Antennariidae *sensu stricto* (Hart et al. [87])

MRCA: *Antennatus coccineus* (Antennariidae) & *Kuiterichthys furcipilis* (Rhycheridae)

Fossil: †*Eophryne barbutii,* MCSNV B.6513 [88]

Notes: Identification as an antennariid on the basis of: general physiognomy of the body, globose shape, mouth large and oblique, simple pectoral fin, sigmoid vertebral column, meristic values of dorsal and anal fins.

Hart et al. [87] broke the former Antennariidae into four families, where Antennariidae is now restricted to the former subfamily "Antennariinae". Carnevale and Pietsch [88] noted similarity of †*Eophryne* to the living genera *Antennarius* (which was not considered distinct from *Fowlerichthys* at the time of their study), *Histrio*, and *Nudiantennarius*. All three of these genera remain in the restricted circumscription of Antennariidae *sensu* Hart et al. [87]. †*Eophryne* has an epural present, which is a diagnostic character of "Antennariinae", though it lacks the other diagnostic character of an endopterygoid being present [89]. This provides justification that †*Eophryne* is at least a crown antennarioid, and (conservatively) a stem antennariid. Past molecular studies instead used †*Eophryne* to calibrate Total group Antennarioidei, whether as a deliberately conservative choice [12] or erroneously [16].

Min. age = 48.5 Ma, Ypresian of Monte Bolca, Pesciara [21]

With †*Plectocretacicus:*

95% soft upper = 71.6 Ma

Mean = 1.49
Standard deviation = 1.0

Outgroup age sequence = 247.1, 193.81, 193.81, 181.7, 164.9, 151.2, 150.89, 103.13, 98.0, 94.0, 94.0, 48.5, 48.5, 48.5

Without †*Plectocretacicus:*

95% soft upper = 73.8 Ma

Mean = 1.59
Standard deviation = 1.0

Outgroup age sequence = 247.1, 193.81, 193.81, 181.7, 164.9, 151.2, 150.89, 103.13, 98.0, 94.0, 48.5, 48.5, 48.5

47. Total group *Fowlerichthys,* or Crown Antennariidae *sensu* Hart et al. [87]

MRCA: *Fowlerichthys avalonis* (Antennariidae) & *Antennatus coccineus* (Antennariidae)

Fossil: †*Neilpeartia ceratoi,* CMC 3, a nearly complete and well-preserved articulated skeleton [89]

Notes: While Histiophryninae had not yet been separated from Antennariidae at the time this fossil was published [87], Carnevale et al. [89] pointed out that †*Neilpeartia* has diagnostic features of "Antennariinae" (now Antennariidae *sensu stricto*): both endopterygoid and epural present. Note that the epural is also present in †*Eophryne*. A sister group relationship with the extant genus *Fowlerichthys* is supported by the five distally branched pelvic fin rays. This would place †*Neilpeartia* in crown "Antennariinae". *Fowlerichthys* is the sister to all other "antennariines" and is separated from them by a long internal branch [90]. Therefore, this calibration is justifiable for crown "Antennariinae" or Antennariidae *sensu stricto* [87]. This represents the "oldest known unquestionable evidence of crown antennariids in the fossil record" [89]. An alternative, more conservative placement of this calibration would be at Total group "Antennariinae", which would make it redundant with †*Eophryne*.

Min. age = 48.5 Ma, Bolca, Pesciara site, Italy[21]

With †*Plectocretacicus:*

95% soft upper = 61.9 Ma

Mean = 0.95
Standard deviation = 1.0

Outgroup age sequence = 247.1, 193.81, 193.81, 181.7, 164.9, 151.2, 150.89, 103.13, 98.0, 94.0, 94.0, 48.5, 48.5, 48.5, 48.5

Without †*Plectocretacicus:*

95% soft upper = 63.3 Ma

Mean = 1.04
Standard deviation = 1.0

Outgroup age sequence = 247.1, 193.81, 193.81, 181.7, 164.9, 151.2, 150.89, 103.13, 98.0, 94.0, 48.5, 48.5, 48.5, 48.5

48. Total group Brachionichthyidae

MRCA: *Brachionichthys australis* & *Kuiterichthys furcipilis*

Fossil: †*Histionotophrous bassani,* MGPD 68487, and †*Orrichthys longimanus,* MCSNV T.160/161. †*Histionotophrous* is known from at least a dozen specimens including complete skeletons. †*Orrichthys* is based on two nearly complete skeletons.

Notes: Pietsch [91] suggested the fossil †*Histionotophrous* was a brachionichthyid instead of an antennariid, and even supposed it could be synonymous with the living genus *Brachionichthys.* Phylogenetic analysis by Carnevale and Pietsch [92] using a matrix of 36 characters found †*Histionotophrous* and †*Orrichthys* to be sister genera. This clade is sister to the living brachionichthyid genera *Brachionichthys* and *Sympterichthys.* This analysis confirms that both fossil genera can be classified as stem brachionichthyids.

Min. age = 48.5 Ma, Bolca, Pesciara site, Italy [21]

With †*Plectocretacicus:*

95% soft upper = 61.9 Ma

Mean = 0.95
Standard deviation = 1.0

Outgroup age sequence = 247.1, 193.81, 193.81, 181.7, 164.9, 151.2, 150.89, 103.13, 98.0, 94.0, 94.0, 48.5, 48.5, 48.5, 48.5

Without †*Plectocretacicus:*

95% soft upper = 63.3 Ma

Mean = 1.04

Standard deviation = 1.0

Outgroup age sequence = 247.1, 193.81, 193.81, 181.7, 164.9, 151.2, 150.89, 103.13, 98.0, 94.0, 48.5, 48.5, 48.5, 48.5

49. Total group *Oneirodes*

MRCA: *Oneirodes kreffti* & *Chaenophryne draco*

Fossil: *Oneiroides sp.,* ZIN 461p, a nearly complete, partly disarticulated skeleton [93].

Notes: The species identity is unclear, but fossil is assignable to *Oneirodes* on the basis of its anteriorly bifurcated frontal bones, the absence of pelvic fins and scales, presence of sphenotic spines, and meristics. This is the oldest record of *Oneirodes* in the fossil record, supplanting the 7.6 million-year-old fossil from the Puente Formation [94] used as a calibration in past molecular studies [16,95].

Min. age = 13.1 Ma, Kurasi Formation, Sakhalin, Russia (see discussion and references in [40,42])

With †*Plectocretacicus:*

95% soft upper = 79.8 Ma

Mean = 2.55
Standard deviation = 1.0

Outgroup age sequence = 247.1, 193.81, 193.81, 181.7, 164.9, 151.2, 150.89, 103.13, 98.0, 94.0, 94.0, 48.5, 13.1

Without †*Plectocretacicus:*

95% soft upper = 83.5 Ma

Mean = 2.61
Standard deviation = 1.0

Outgroup age sequence = 247.1, 193.81, 193.81, 181.7, 164.9, 151.2, 150.89, 103.13, 98.0, 94.0, 48.5, 13.1

**Geologic calibrations:**

Calibrations 50–64 are geologic calibrations placed on the age of sister-species pairs separated by the Isthmus of Panama. Following Rincon-Sandoval et al. [66], we use 2.8 Ma as the minimum age for these calibrations reflecting the undisputed minimum geologic age for the closure of the Isthmus of Panama.

All geologic calibrations were set as an exponential distribution with an offset of 2.8 and a mean of 1.0.

**Table S5: List of geologic calibrations.**

| Cal. No. | Family | East Pacific species | West Atlantic species |
|---|---|---|---|
| 50 | Scorpaenidae | *Scorpaena mystes* | *Scorpaena plumieri* |
| 51 | Epinephelidae | *Paranthias colonus* | *Paranthias furcifer* |
| 52 | Epinephelidae | *Rypticus bicolor* | *Rypticus saponaceus* |
| 53 | Gerreidae | *Eucinostomus currani* | *Eucinostomus melanopterus* |
| 54 | Sciaenidae | *Bairdiella armata* | *Bairdiella ronchus* |
| 55 | Sciaenidae | *Umbrina xanti* | *Umbrina coroides* |
| 56 | Haemulidae | *Anisotremus interruptus* | *Anisotremus surinamensis* |
| 57 | Haemulidae | *Rhonciscus bayanus* | *Rhonciscus crocro* |
| 58 | Lutjanidae | *Lutjanus argentiventris* | *Lutjanus purpureus*/*L. griseus* (ASTRAL gene-subset 4 only) |
| 59 | Lutjanidae | *Lutjanus guttatus* | *Lutjanus synagris*/*L. mahogoni* (ASTRAL gene-subsets 2 and 11 only) |
| 60 | Lutjanidae | *Lutjanus novemfasciatus* | *Lutjanus cyanopterus* |
| 61 | Pomacanthidae | *Holacanthus passer* | *Holacanthus ciliaris* |
| 62 | Tetraodontidae | *Canthigaster punctatissima* | *Canthigaster rostrata* |
| 63 | Tetraodontidae | *Sphoeroides annulatus* | *Sphoeroides testudineus* |
| 64 | Tetraodontidae | *Sphoeroides lobatus* | *Sphoeroides maculatus* |

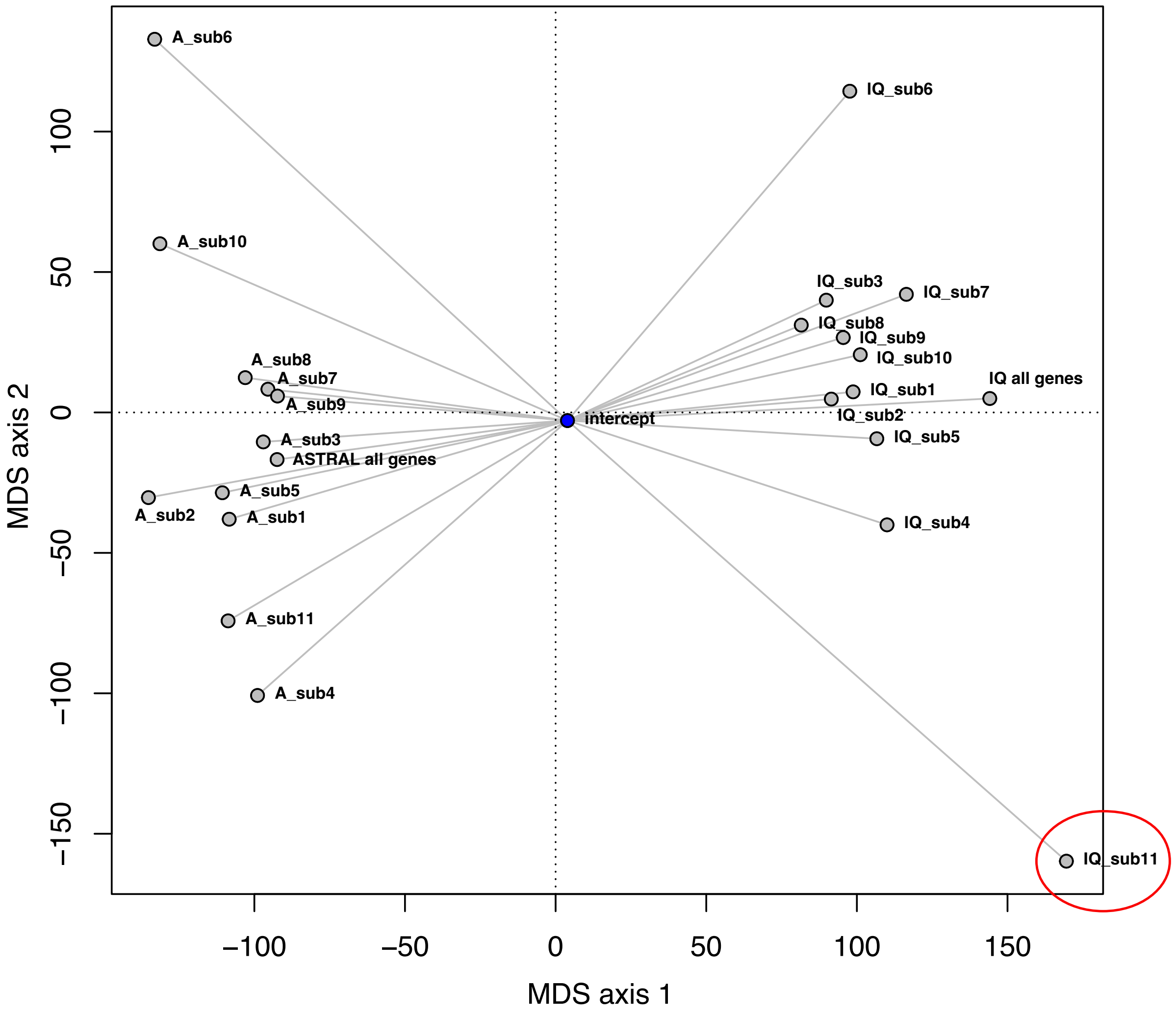


**Figure S1. Tree space plot showing variation among ASTRAL and IQ-TREE "all genes" and "gene subset" trees.** The average (centroid) tree is represented by a blue point ("intercept"). The IQ-TREE made from gene subset 11 (red oval) was considered an outlier and removed from the study.

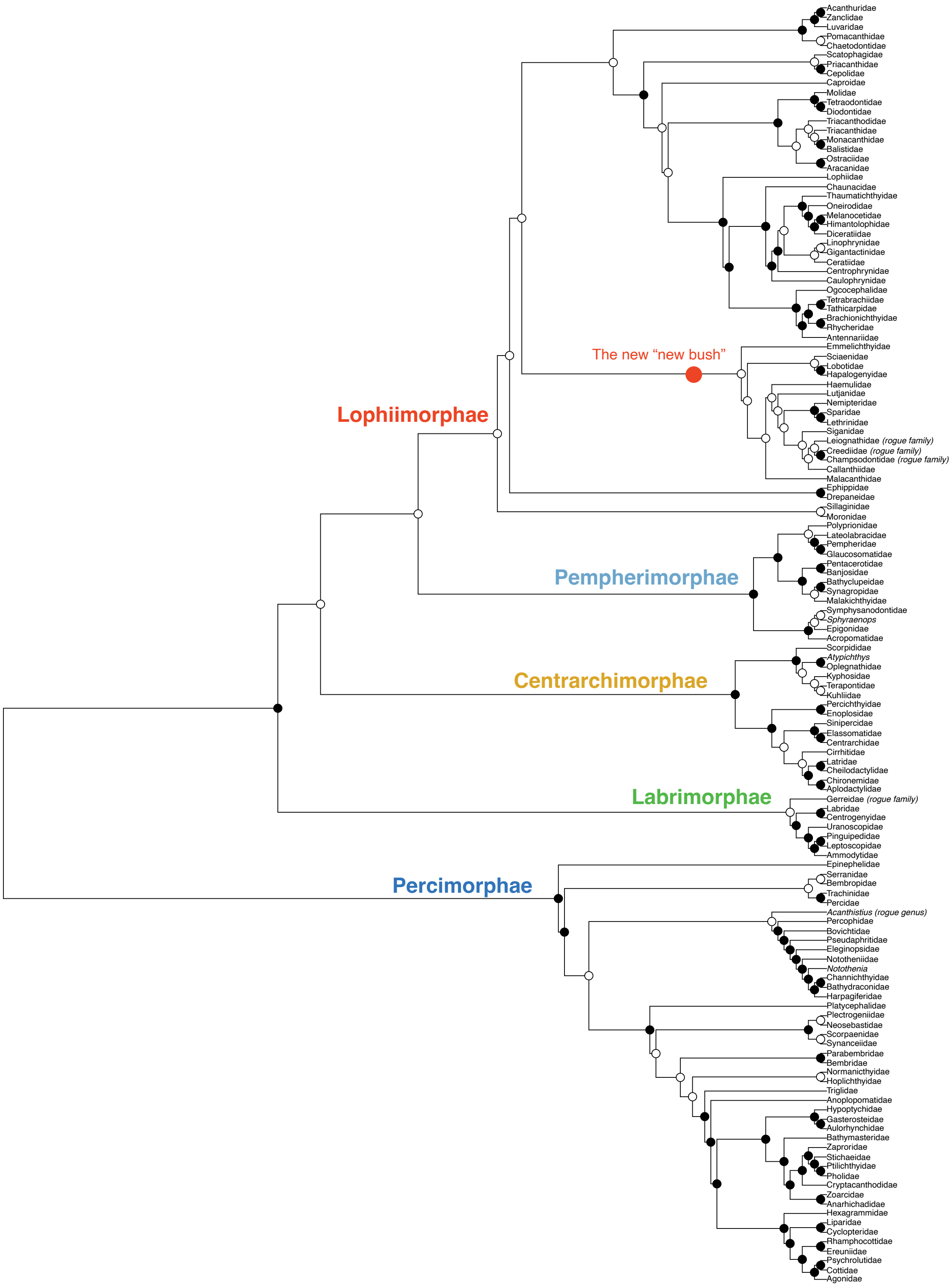


**Figure S2. Comparing the family-level topologies of the two "all genes" trees.** The IQ-TREE cladogram trimmed to one tip per family is shown. Circles on nodes indicate the match to the ASTRAL tree, where black indicates that the same families form a clade in both trees. White indicates that the families that form that clade in the IQ-TREE do not form a clade in the ASTRAL tree (one or more families found outside that node).

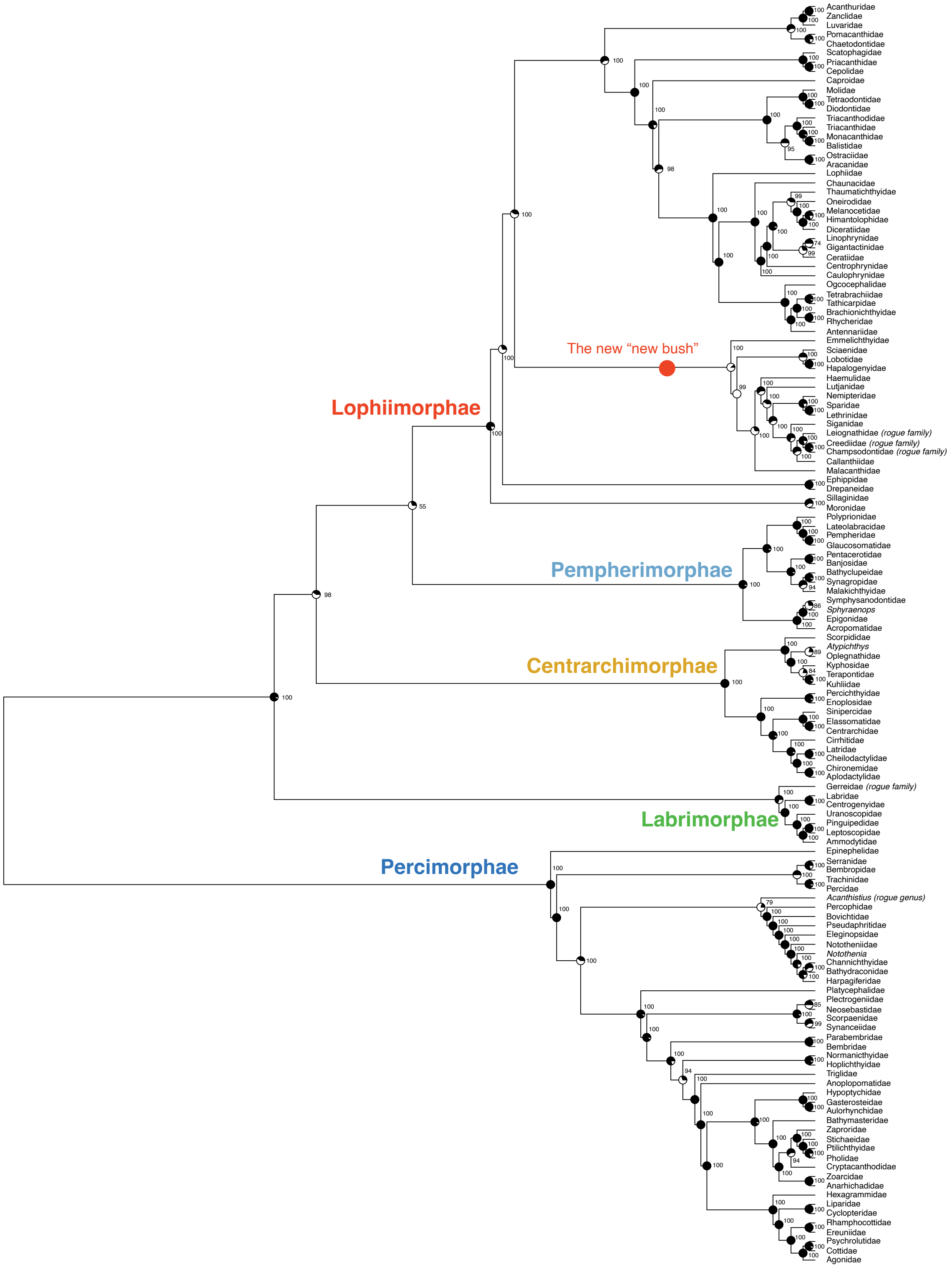


**Figure S3. Comparing the family-level topologies of all concatenated trees.** The IQ-TREE “all genes” cladogram trimmed to one tip per family is shown; support values are given based on 1000 UFboot simulations. Pies on nodes indicate the degree of matching to the ten IQ-TREE gene subset topologies, where more black indicates that a greater number of gene subset trees match the topology of the “all genes” tree.

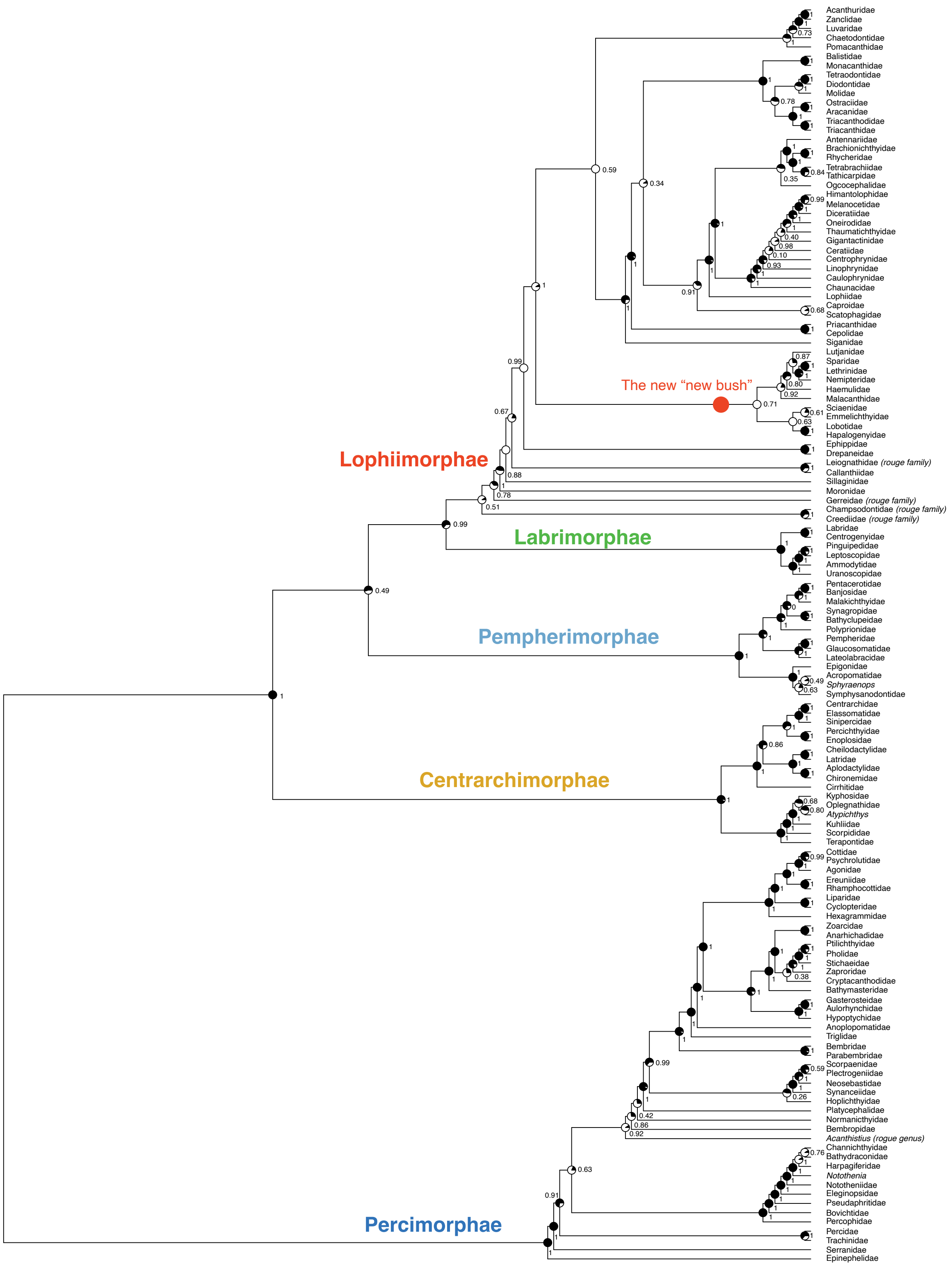


**Figure S4. Comparing the family-level topologies of all coalescent species trees.** The ASTRAL “all genes” cladogram trimmed to one tip per family is shown; support values are given based on local posterior probabilities. Pies on nodes indicate the degree of matching to the eleven ASTRAL gene subset topologies, where more black indicates that a greater number of gene subset trees match the topology of the “all genes” tree.

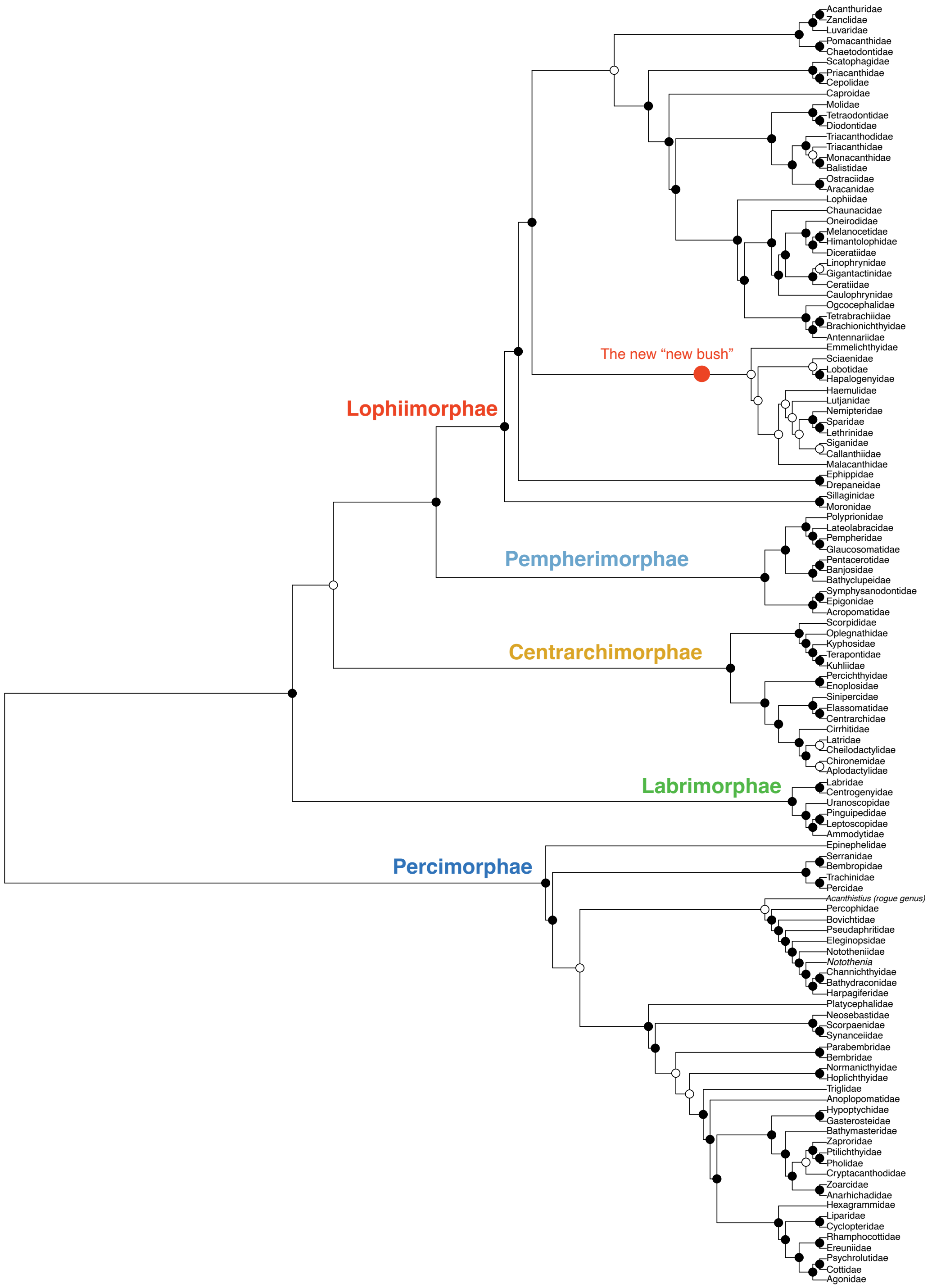


**Figure S5. Comparing the family-level topologies of our IQ-TREE “all genes” tree versus Ghezelayagh et al.** [1]. The tree used for this comparison is the “1100taxa_NoDups_FINAL.tre” file in the Dryad package of that study. The IQ-TREE “all genes” cladogram trimmed to one tip per family is shown. Circles on nodes indicate the match to the comparison tree, where black indicates that the same families form a clade in both trees. White indicates the clade was found in our study but not the comparison tree (one or more families found outside that node). Rogue families (Gerreidae, Leiognathidae, Creediidae, Champsodontidae) were removed for this comparison.

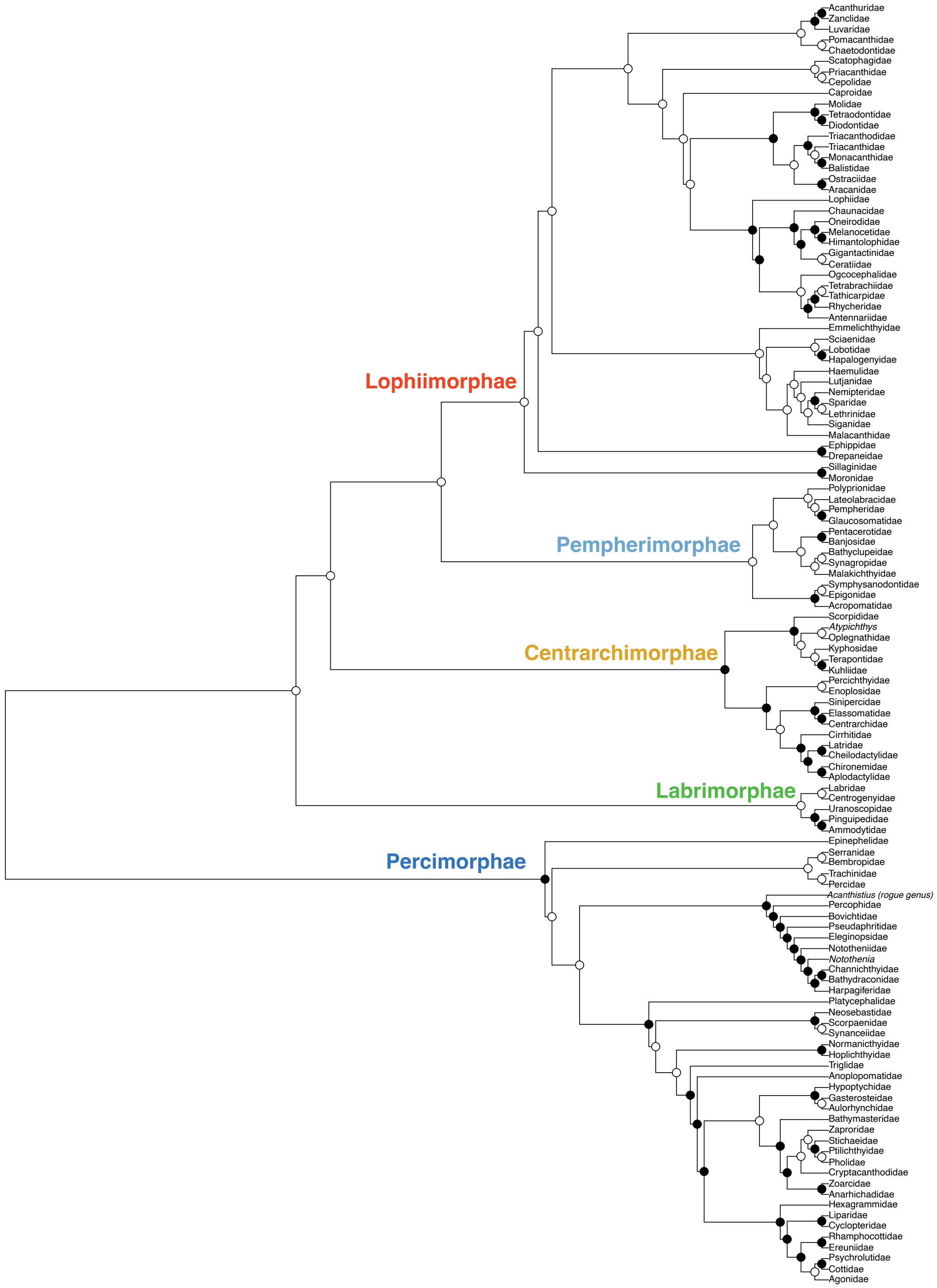


**Figure S6. Comparing the family-level topologies of our IQ-TREE "all genes" tree versus Rabosky et al.** [2]. The tree used for this comparison is the "actinopt_12k_treePL.tre" file in the Dryad package of that study (tree including species with genetic data only). The IQ-TREE "all genes" cladogram trimmed to one tip per family is shown. Circles on nodes indicate the match to the comparison tree, where black indicates that the same families form a clade in both trees. White indicates the clade was found in our study but not the comparison tree (one or more families found outside that node). Rogue families (Gerreidae, Leiognathidae, Creediidae, Champsodontidae) were removed for this comparison.

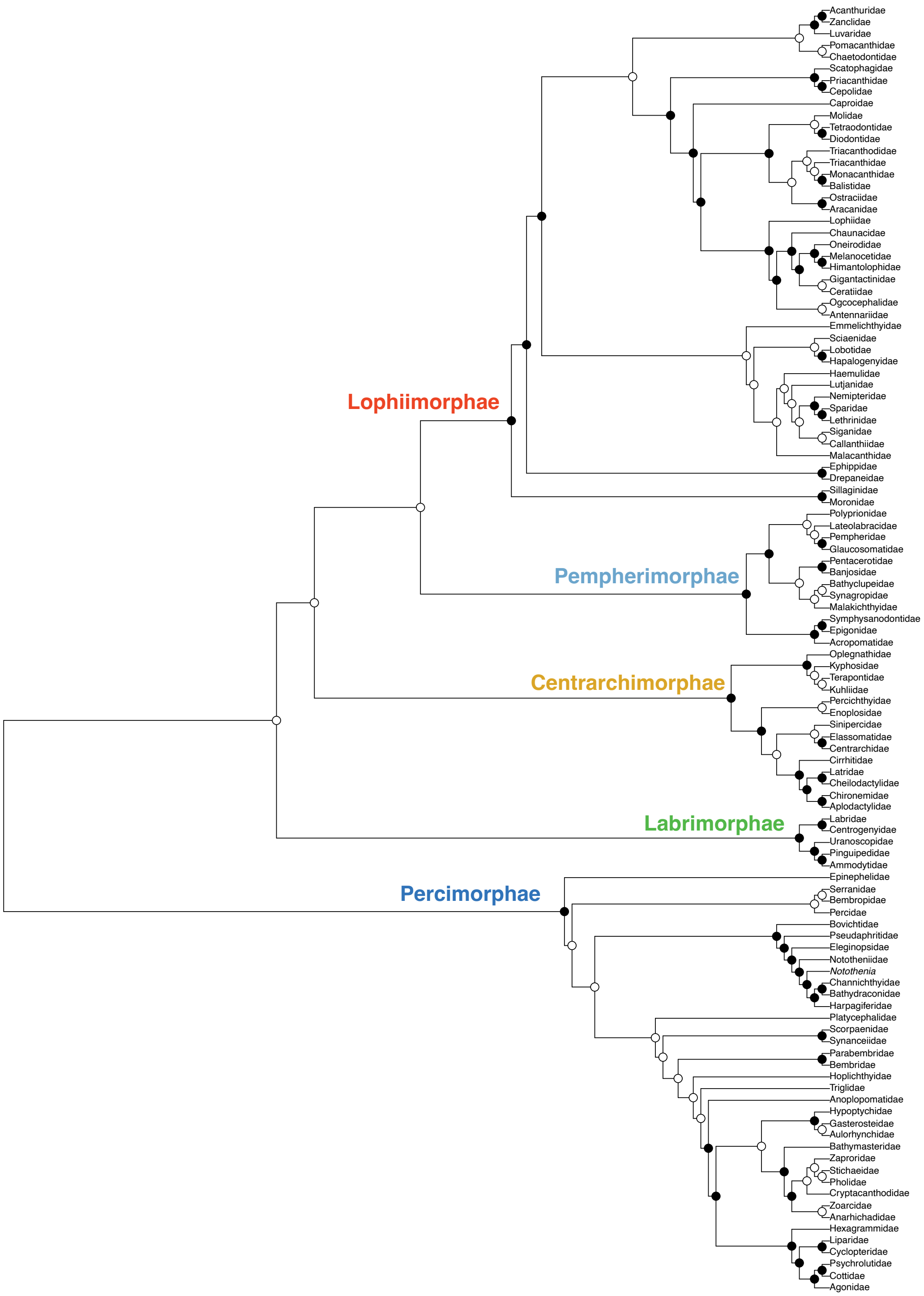


**Figure S7. Comparing the family-level topologies of our IQ-TREE "all genes" tree versus Betancur-R et al.**[3]. The tree used for this comparison is shown in Fig. 2 of that study. The IQ-TREE "all genes" cladogram trimmed to one tip per family is shown. Circles on nodes indicate the match to the comparison tree, where black indicates that the same families form a clade in both trees. White indicates the clade was found in our study but not the comparison tree (one or more families found outside that node). Rogue families (Gerreidae, Leiognathidae, Creediidae, Champsodontidae) were removed for this comparison.

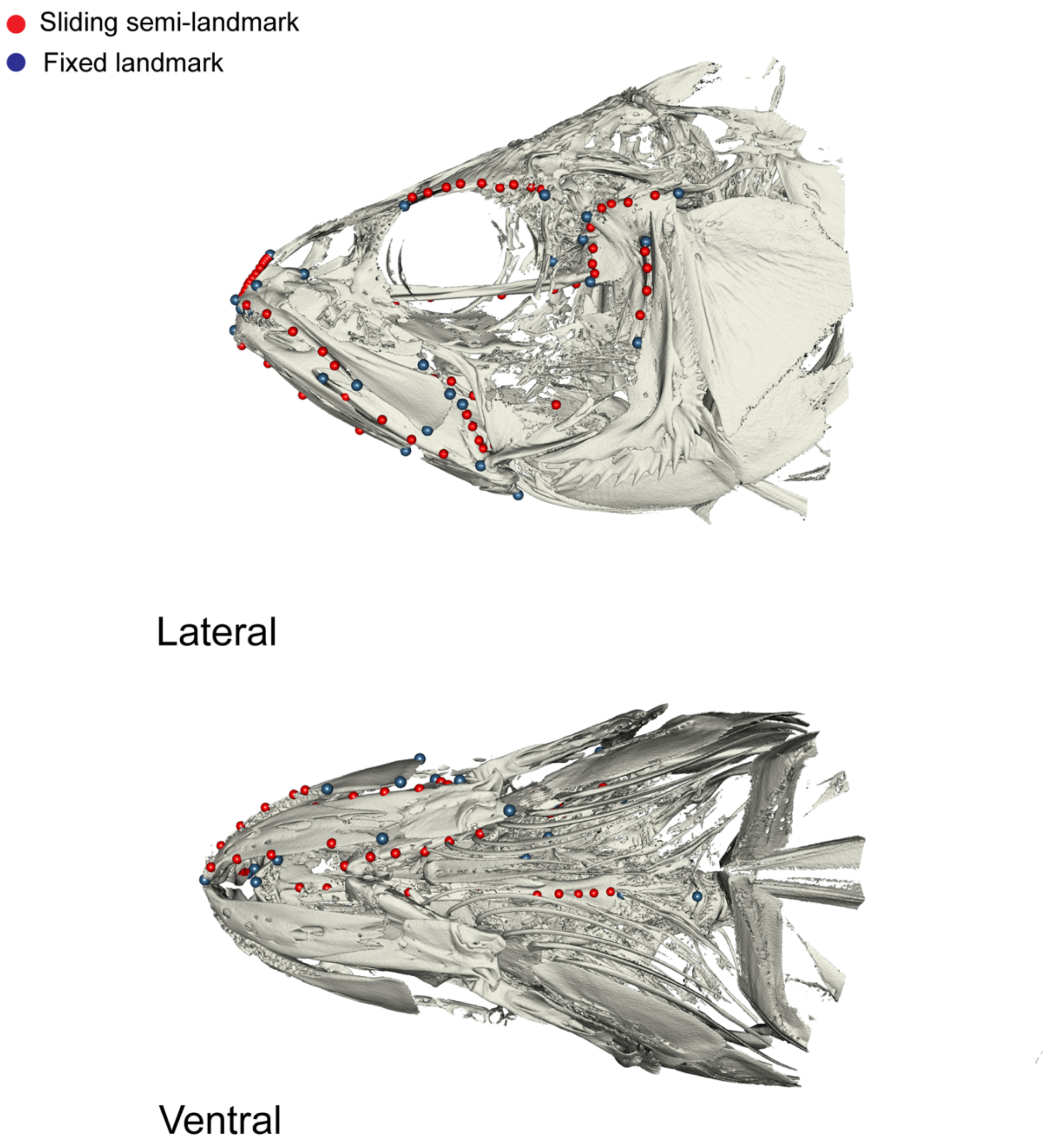


**Figure S8. Landmarks applied to CT scans of fish skulls.** Red points indicate curves.

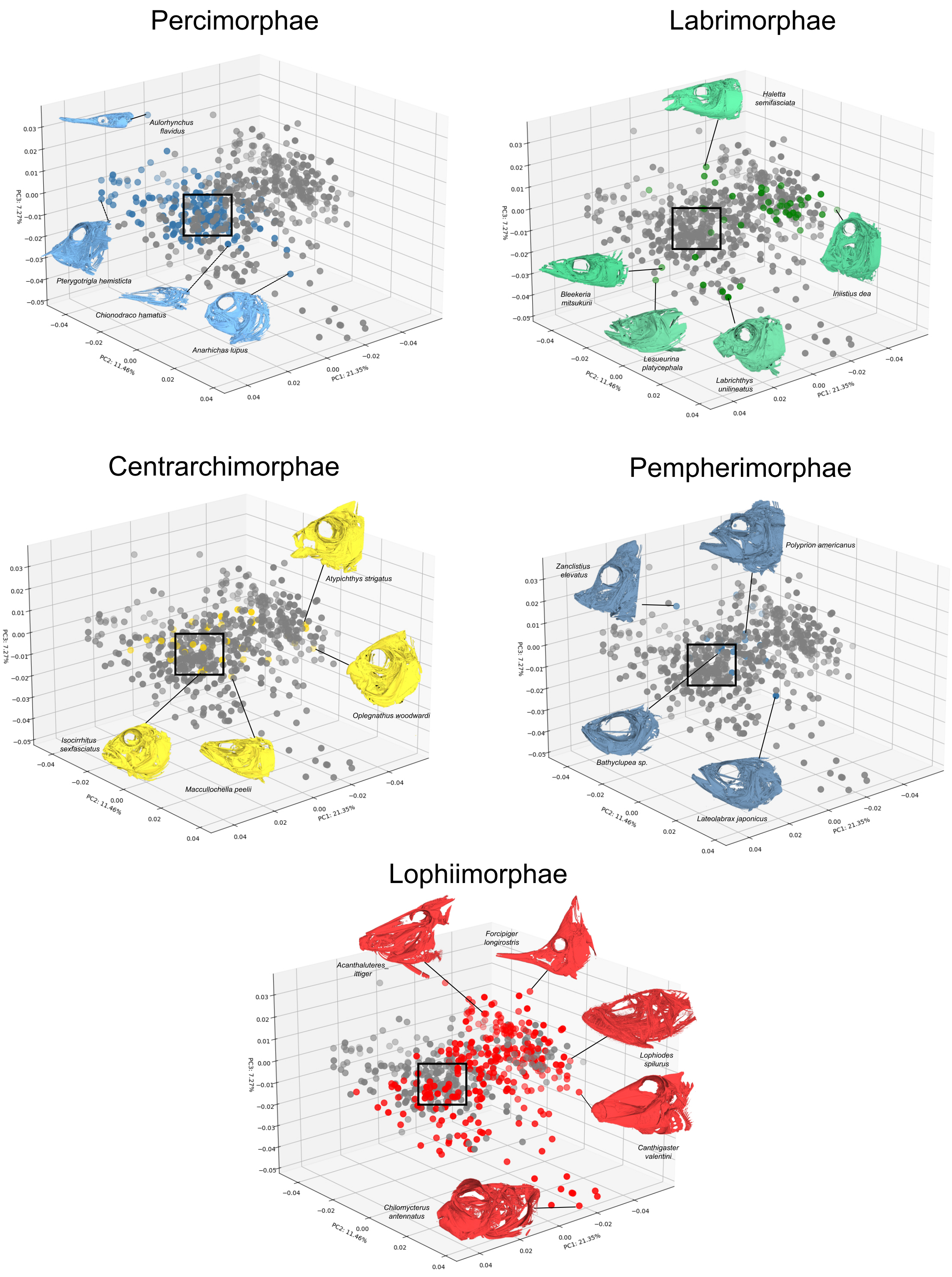


**Figure S9: Skull shape diversity by superorder**. Morphospaces based on 571 species of eupercarian fishes with the primary three axes of skull shape variation shown. Insets depict representative skull shapes for different regions of shape space. Square indicates the “Percomorph Pile” of Figure 2. See Data S1 and Table S3 for details of the five eupercarian superorders.

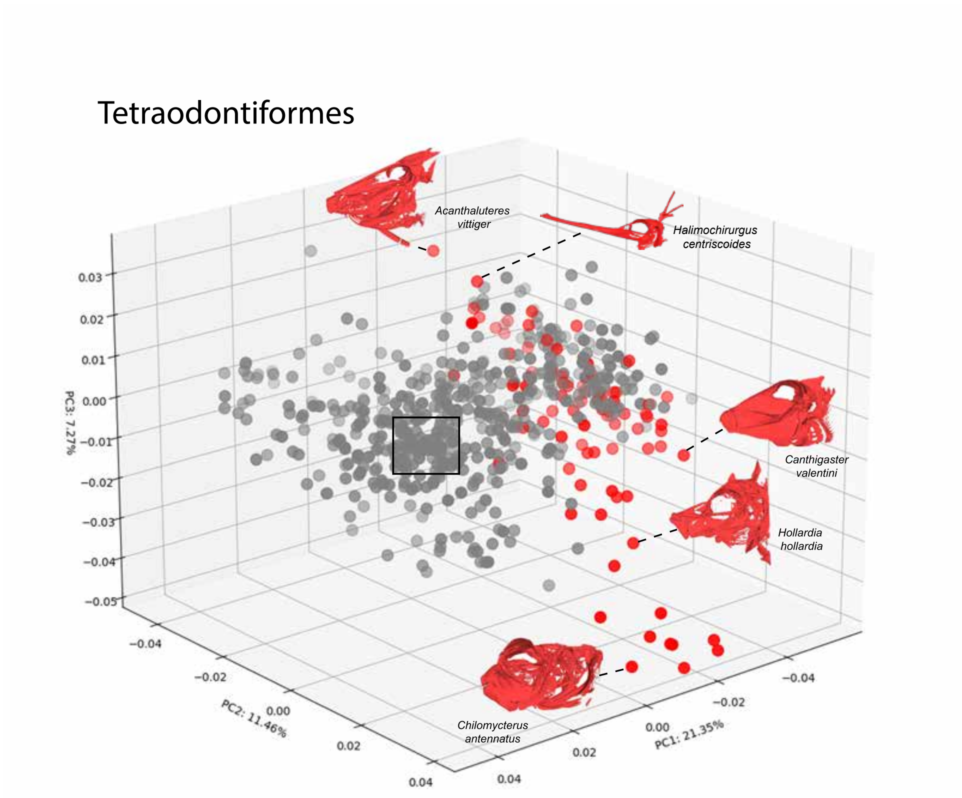


**Figure S10. Skull shape diversity among Tetraodontiformes.** Grey points indicate non-tetraodontiform eupercarians. Square indicates the “Percomorph Pile” of Figure 2.

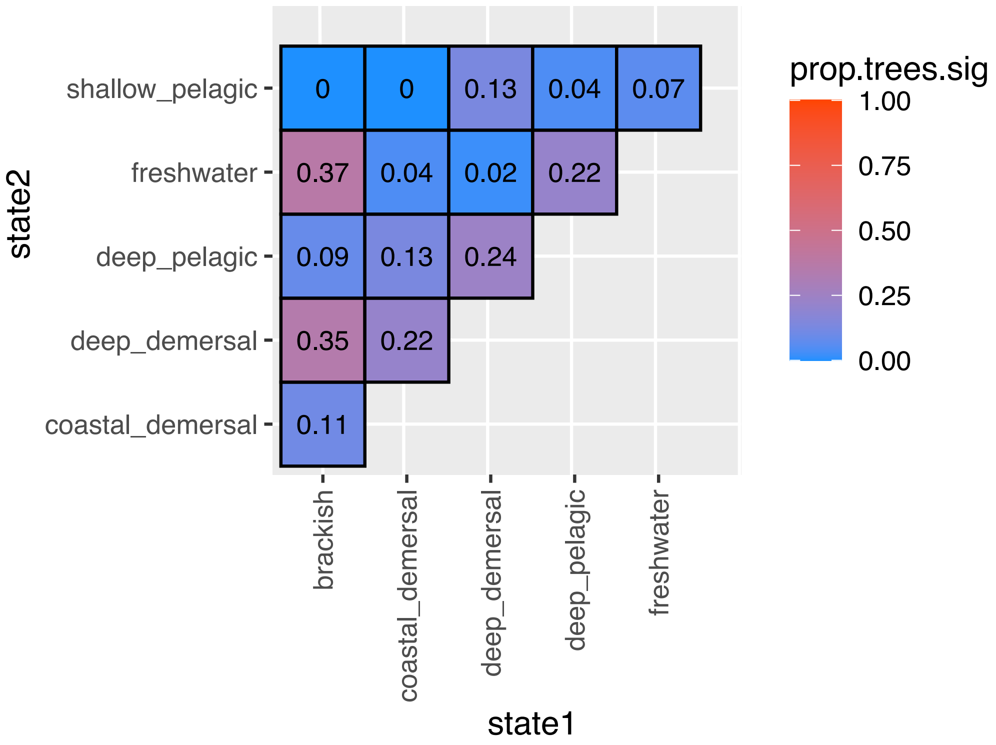


**Figure S11.** Results of PGLS post-hoc comparisons. Pairwise comparisons show the proportion of trees (out of 46) in which skull shapes were significantly different between habitats.

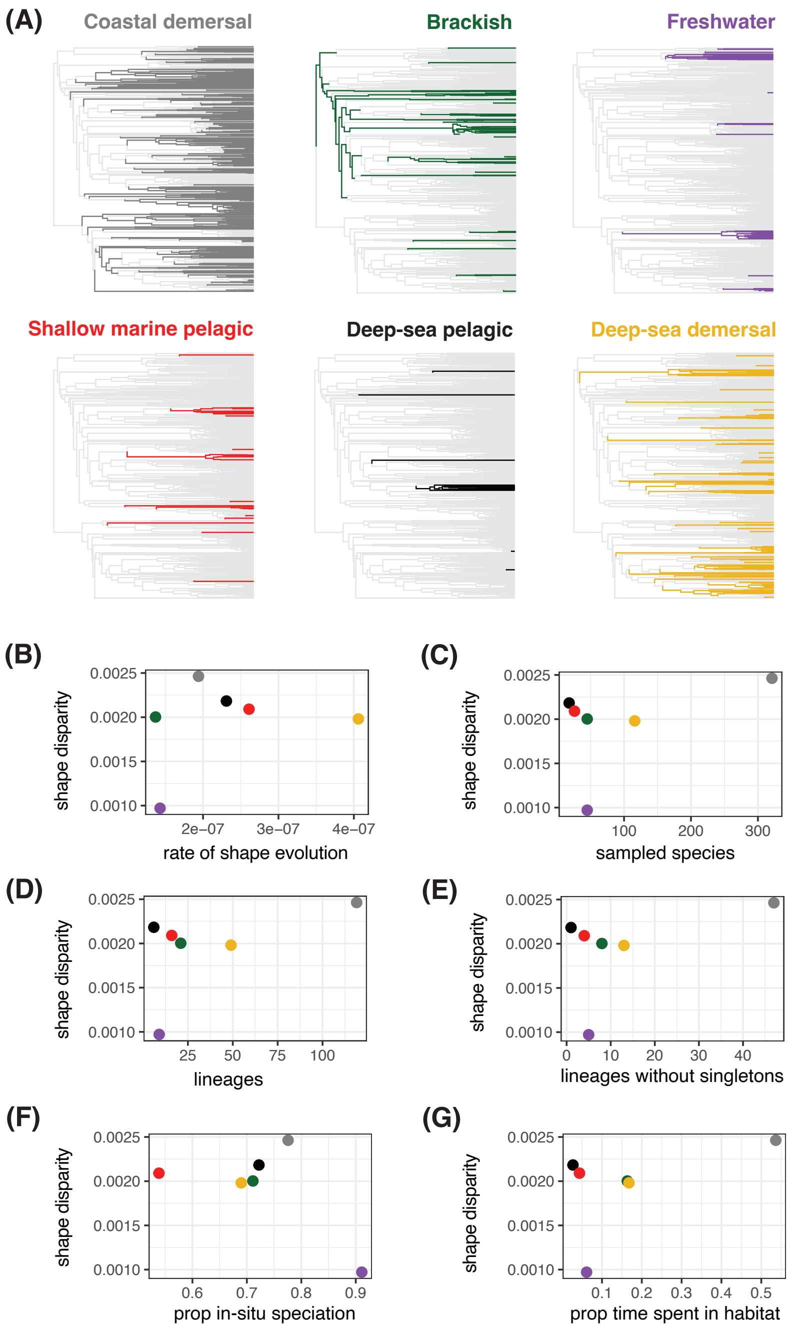


**Figure S12. Routes to accumulating phenotypic disparity, based on ASTRAL -Plecto tree.** (A) Ancestral habitat reconstructions, with each habitat shown individually for visual clarity. (B–G) Disparity in skull shape by habitat, related to six factors: (B) rate of skull shape evolution (inferred using geomorph R package), (C) number of species in each habitat sampled in CT scan dataset, (D) number of phylogenetically independent lineages in each habitat, as inferred from ancestral state reconstructions, (E) count of phylogenetically independent lineages without singletons (habitat transition associated with one terminal), (F) the proportion of species in each habitat derived from in-situ speciation, as opposed to recent colonization (transitions associated with terminal branches), (G) the proportion of total evolutionary time spent in the habitat as inferred by SIMMAP.

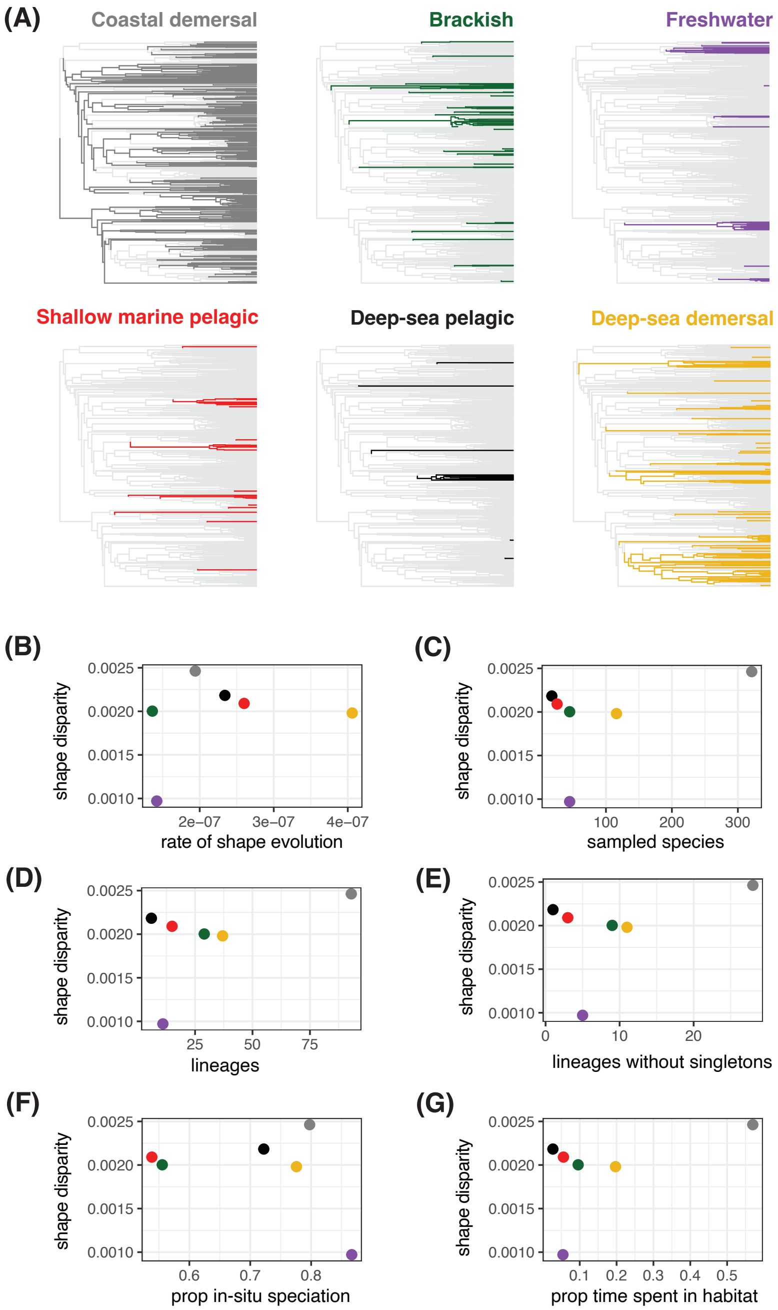


**Figure S13. Routes to accumulating phenotypic disparity, based on ASTRAL +Plecto tree.** (A) Ancestral habitat reconstructions, with each habitat shown individually for visual clarity. (B–G) Disparity in skull shape by habitat, related to six factors: (B) rate of skull shape evolution (inferred using geomorph R package), (C) number of species in each habitat sampled in CT scan dataset, (D) number of phylogenetically independent lineages in each habitat, as inferred from ancestral state reconstructions, (E) count of phylogenetically independent lineages without singletons (habitat transition associated with one terminal), (F) the proportion of species in each habitat derived from in-situ speciation, as opposed to recent colonization (transitions associated with terminal branches), (G) the proportion of total evolutionary time spent in the habitat as inferred by SIMMAP.

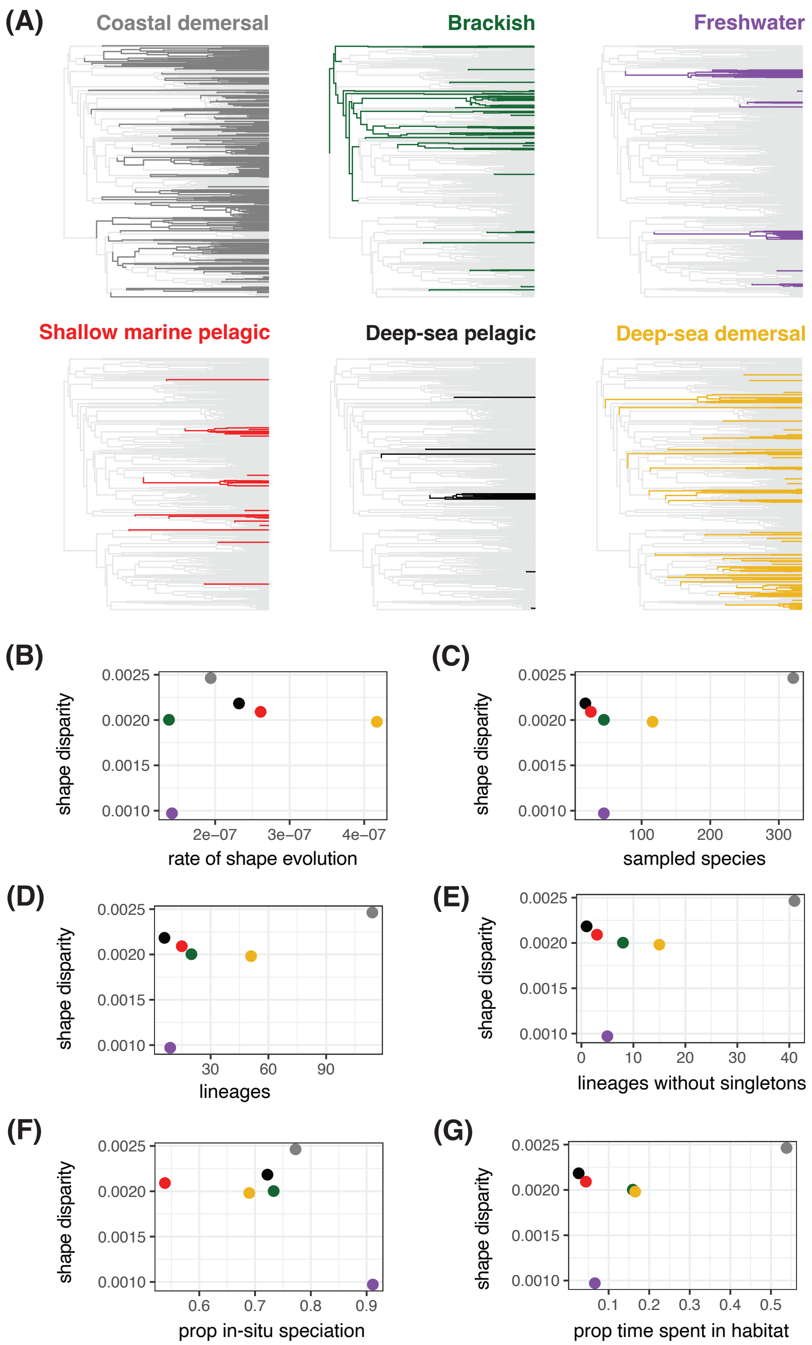


**Figure S14. Routes to accumulating phenotypic disparity, based on IQ-TREE +Plecto tree.** (A) Ancestral habitat reconstructions, with each habitat shown individually for visual clarity. (B–G) Disparity in skull shape by habitat, related to six factors: (B) rate of skull shape evolution (inferred using geomorph R package), (C) number of species in each habitat sampled in CT scan dataset, (D) number of phylogenetically independent lineages in each habitat, as inferred from ancestral state reconstructions, (E) count of phylogenetically independent lineages without singletons (habitat transition associated with one terminal), (F) the proportion of species in each habitat derived from in-situ speciation, as opposed to recent colonization (transitions associated with terminal branches), (G) the proportion of total evolutionary time spent in the habitat as inferred by SIMMAP

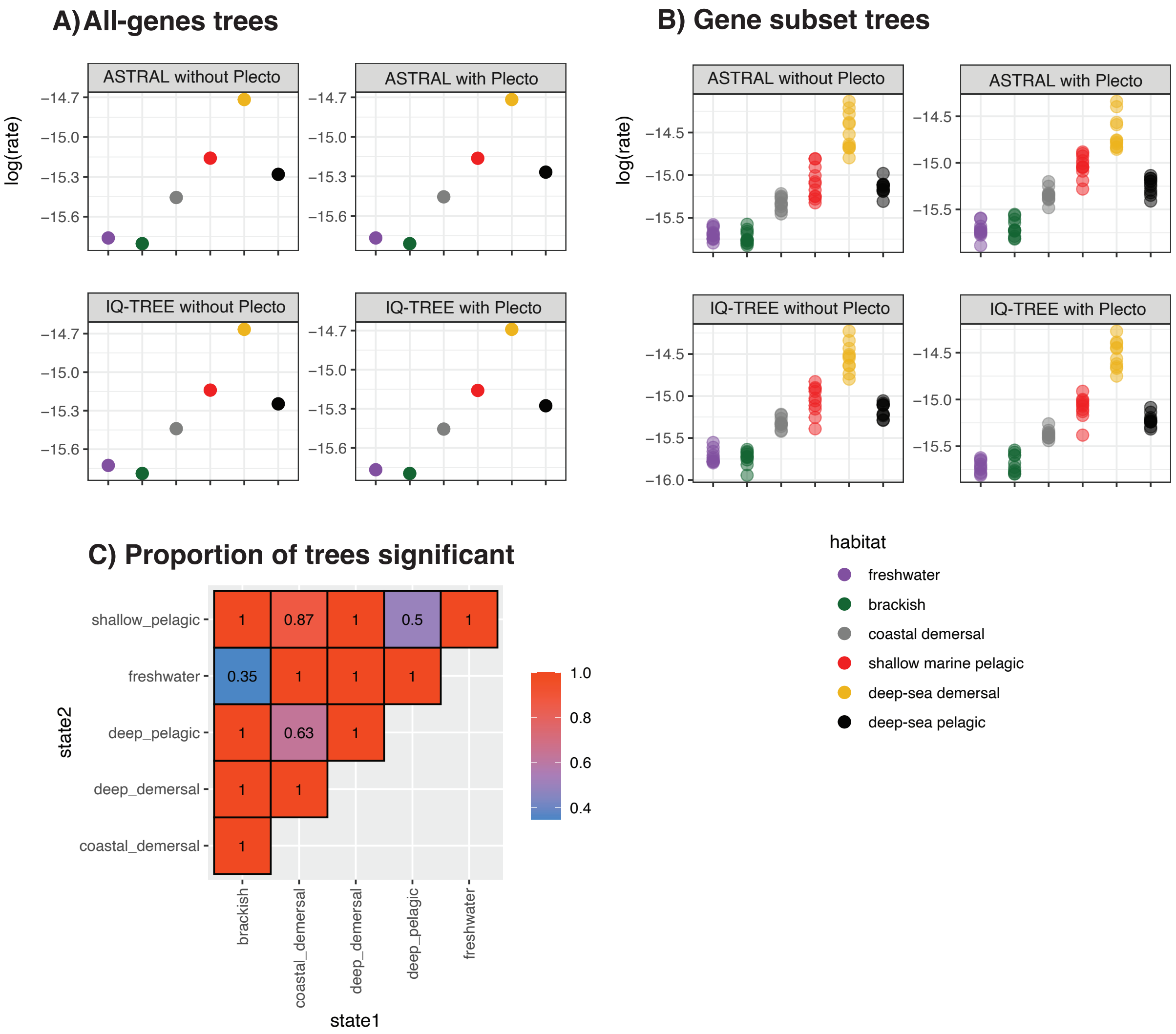


**Figure S15. Rates of skull shape evolution by habitat.** Variation in rates estimated using geomorph[4,5] are shown for the four "all-genes" trees (A) and the 42 gene-subset tree variants (B). Panel (C) shows the proportion of input trees (out of 46) for which each pairwise comparison was significant (i.e. $p<0.05$).

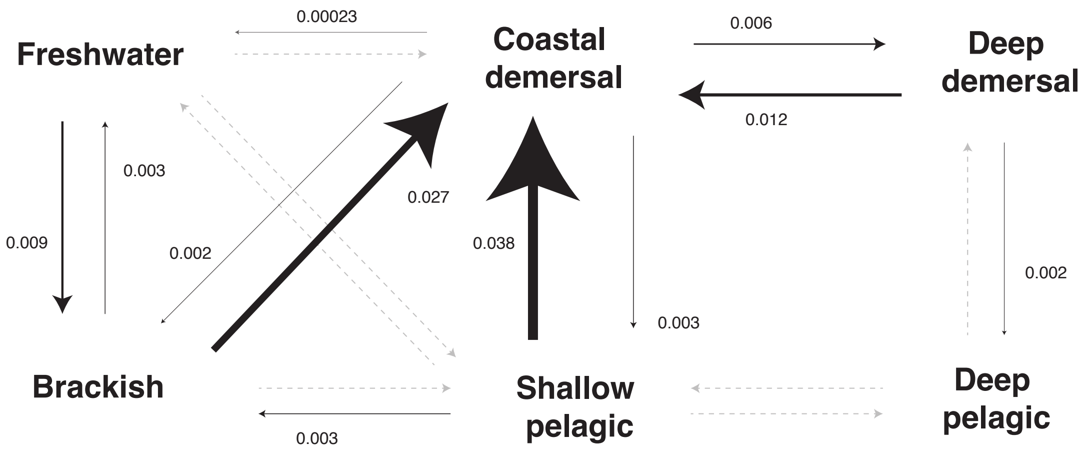


**Figure S16. Transition rates among habitats inferred using SIMMAP based on ASTRAL -Plecto tree.** Arrows are scaled by the magnitude of the transition rate. Arrows in light grey represent transitions that were allowed by the evolutionary model but not observed. See Table S9 for transitions not allowed by the evolutionary model used in SIMMAP.

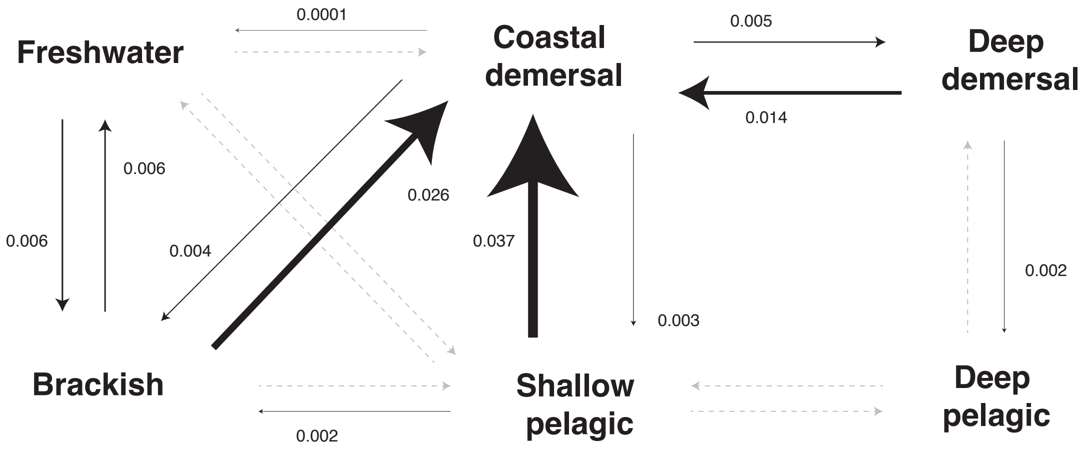


**Figure S17. Transition rates among habitats inferred using SIMMAP based on ASTRAL +Plecto tree.** Arrows are scaled by the magnitude of the transition rate. Arrows in light grey represent transitions that were allowed by the evolutionary model but not observed. See Table S9 for transitions not allowed by the evolutionary model used in SIMMAP.

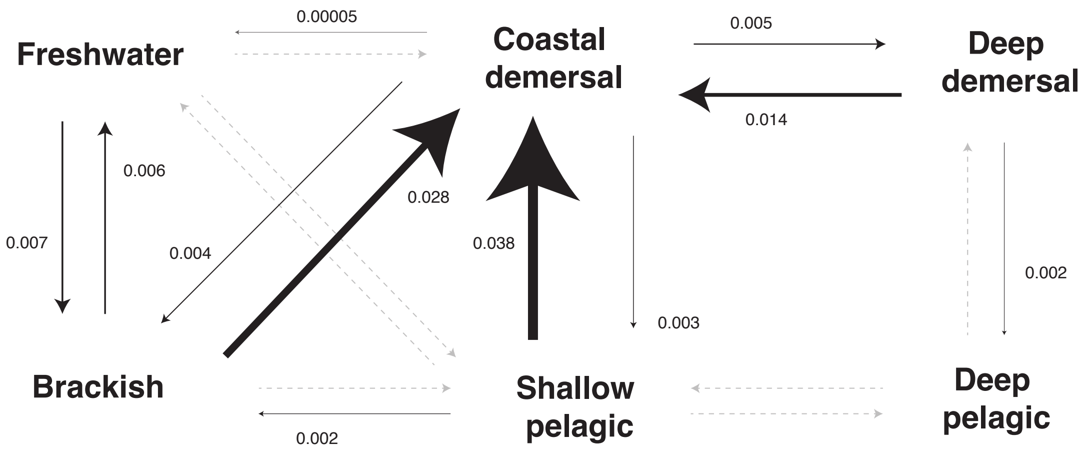


**Figure S18. Transition rates among habitats inferred using SIMMAP based on IQ-TREE -Plecto tree.** Arrows are scaled by the magnitude of the transition rate. Arrows in light grey represent transitions that were allowed by the evolutionary model but not observed. See Table S9 for transitions not allowed by the evolutionary model used in SIMMAP.

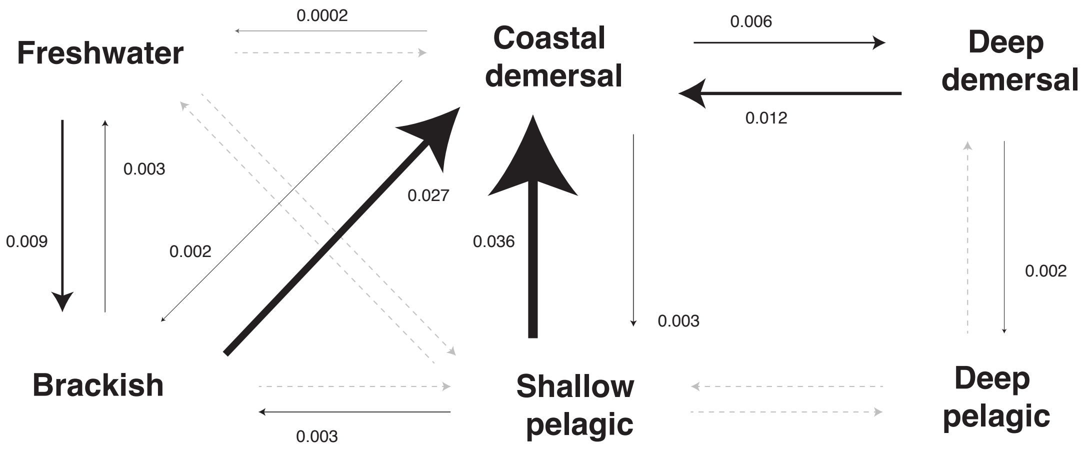


**Figure S19. Transition rates among habitats inferred using SIMMAP based on IQ-TREE +Plecto tree.** Arrows are scaled by the magnitude of the transition rate. Arrows in light grey represent transitions that were allowed by the evolutionary model but not observed. See Table S9 for transitions not allowed by the evolutionary model used in SIMMAP.

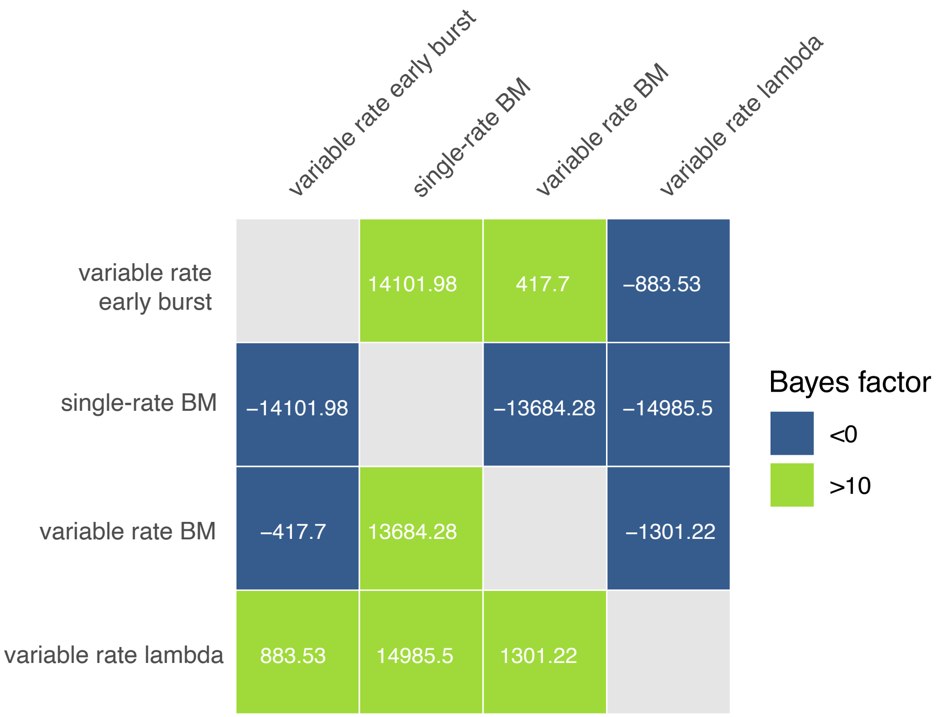


**Figure S20. BayesTraits model comparison using Bayes factors.** Models are fit using transformations of the input tree consistent with predictions of different evolutionary scenarios. BM=Brownian motion.

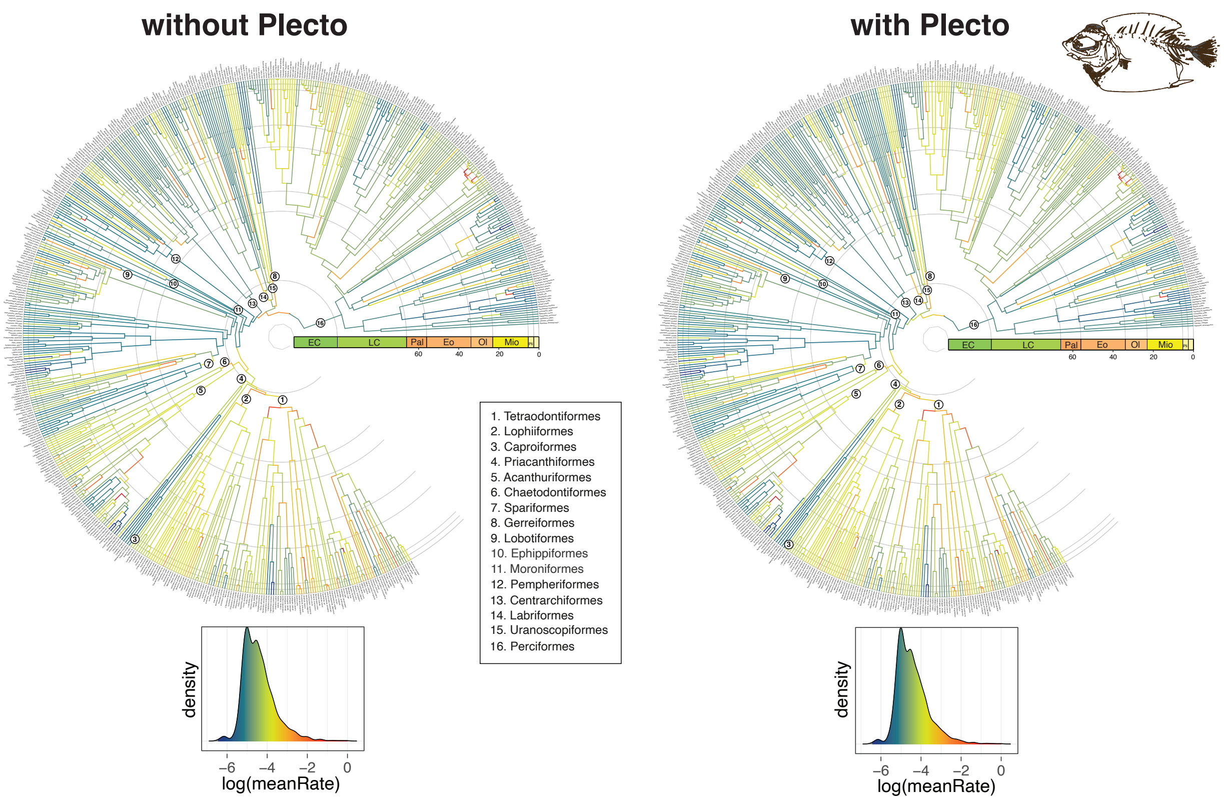


**Figure S21.** Comparing BayesTraits results between +Plecto and -Plecto IQ-TREE variants. Taxonomic orders are labeled according to our new classification (Data S1).

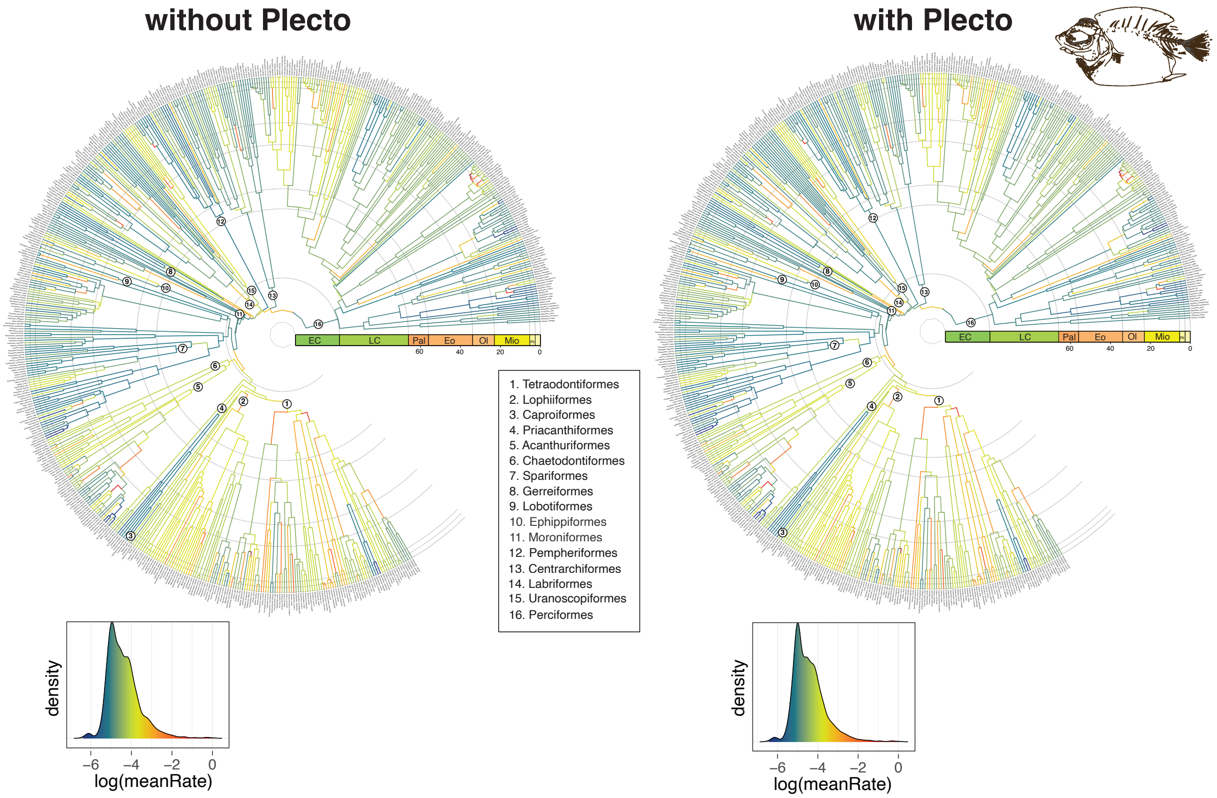


**Figure S22.** Comparing BayesTraits results between +Plecto and -Plecto ASTRAL variants. Taxonomic orders are labeled according to our new classification (Data S1).

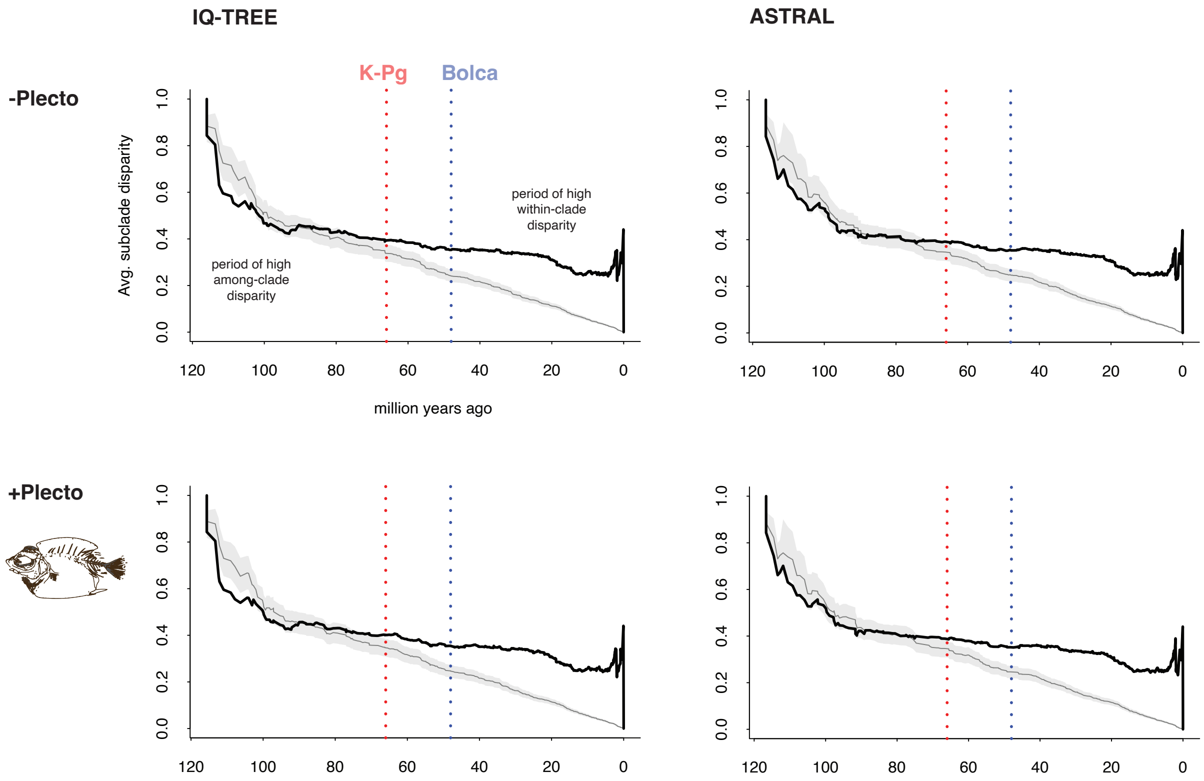


**Figure S23. Biphasic evolution of skull shape disparity through time, based on four input tree variants.** Solid black line shows the observed pattern of disparity through time relative to the Brownian motion null expectation generated using 10,000 simulations (grey shading)[6,7]. Two significant events are marked with a red line (K-Pg mass extinction, 66 million years ago) and blue line (Monte Bolca Lagerstätte, 48.5 million years ago[8]).

**Table S1.** Sampling of Eupercaria in this study versus two recent molecular phylogenies, using the reference list available at www.fishtreeoflife.org.

| Study | Source of data | Percent of species sampled | Percent of genera sampled | Percent of families sampled |
|---|---|---|---|---|
| This study | 1,095 exons | 15.1 | 39.4 | 82.6 |
| Ghezelayagh et al. [1] | 989 UCE loci | 5.6 | 23.9 | 87.6 |
| Rabosky et al. [2] | 27 genes | 46.2 | 66.0 | 86.3 |

***Table S2.** List of genetic material and quality control. Given the substantial file size, this supplementary table is archived on Zenodo.

***Table S3.** Comparing previous classifications of Eupercaria. Given the substantial file size, this supplementary table is archived on Zenodo.

**Table S4**. Age estimates for the root (total group Holocentridae) from previous studies (Also available in Data S2).

| Study | Mean age Root (Ma) | 95% HPD |
|---|---|---|
| Alfaro et al. (2018) [4] | 124 | (112-136) |
| Betancur-R et al. (2013) [9] | 141 | (125-158) |
| Betancur-R et al. (2017) [10] | 145 | |
| Chen et al. (2014) [11] | 125 | (114-136) |
| Ghezelayagh et al. (2022) [12] | 133 | |
| Hughes et al. (2018) [13] | 128 | (123-133) |
| Near et al. (2012) [14] | 126 | (118-136) |
| Near et al. (2013) [15] | 126 | (118-136) |
| Rabosky et al. (2018) [16] | 129 | |

**Table S5: List of geologic calibrations** (Also available in Data S2).

| Cal. No. | Family | East Pacific species | West Atlantic species |
|---|---|---|---|
| 50 | Scorpaenidae | *Scorpaena mystes* | *Scorpaena plumieri* |
| 51 | Epinephelidae | *Paranthias colonus* | *Paranthias furcifer* |
| 52 | Epinephelidae | *Rypticus bicolor* | *Rypticus saponaceus* |
| 53 | Gerreidae | *Eucinostomus currani* | *Eucinostomus melanopterus* |
| 54 | Sciaenidae | *Bairdiella armata* | *Bairdiella ronchus* |
| 55 | Sciaenidae | *Umbrina xanti* | *Umbrina coroides* |
| 56 | Haemulidae | *Anisotremus interruptus* | *Anisotremus surinamensis* |
| 57 | Haemulidae | *Rhonciscus bayanus* | *Rhonciscus crocro* |
| 58 | Lutjanidae | *Lutjanus argentiventris* | *Lutjanus purpureus*/*L. griseus* (ASTRAL gene-subset 4 only) |
| 59 | Lutjanidae | *Lutjanus guttatus* | *Lutjanus synagris*/*L. mahogoni* (ASTRAL gene-subsets 2 and 11 only) |
| 60 | Lutjanidae | *Lutjanus novemfasciatus* | *Lutjanus cyanopterus* |
| 61 | Pomacanthidae | *Holacanthus passer* | *Holacanthus ciliaris* |
| 62 | Tetraodontidae | *Canthigaster punctatissima* | *Canthigaster rostrata* |
| 63 | Tetraodontidae | *Sphoeroides annulatus* | *Sphoeroides testudineus* |
| 64 | Tetraodontidae | *Sphoeroides lobatus* | *Sphoeroides maculatus* |

***Table S6.** List of voucher material used for CT scanning and habitat codes. Given the substantial file size, this supplementary table is archived on Zenodo.

**Table S7.** Results of PGLS regressions of habitat against three-dimensional cranial shape (*n*=571 species) across 46 alternative input phylogenies.

| Molecular dataset | Tree estimation | Fossil calibration scheme | P-value | $R^2$ |
|---|---|---|---|---|
| All genes | ASTRAL | +Plecto | 0.0016 | 0.027 |
| All genes | ASTRAL | - Plecto | 0.0016 | 0.027 |
| All genes | IQ-TREE | +Plecto | 0.0002 | 0.035 |
| All genes | IQ-TREE | - Plecto | 0.0001 | 0.035 |
| Gene subset 1 | ASTRAL | +Plecto | 0.0001 | 0.084 |
| Gene subset 2 | ASTRAL | +Plecto | 0.0001 | 0.048 |
| Gene subset 3 | ASTRAL | +Plecto | 0.0001 | 0.068 |
| Gene subset 4 | ASTRAL | +Plecto | 0.0001 | 0.033 |
| Gene subset 5 | ASTRAL | +Plecto | 0.0001 | 0.057 |
| Gene subset 6 | ASTRAL | +Plecto | 0.0001 | 0.055 |
| Gene subset 7 | ASTRAL | +Plecto | 0.0002 | 0.044 |
| Gene subset 8 | ASTRAL | +Plecto | 0.0001 | 0.039 |
| Gene subset 9 | ASTRAL | +Plecto | 0.0012 | 0.043 |
| Gene subset 10 | ASTRAL | +Plecto | 0.0001 | 0.137 |
| Gene subset 11 | ASTRAL | +Plecto | 0.0001 | 0.140 |
| Gene subset 1 | ASTRAL | - Plecto | 0.0027 | 0.055 |
| Gene subset 2 | ASTRAL | - Plecto | 0.0001 | 0.145 |
| Gene subset 3 | ASTRAL | - Plecto | 0.0001 | 0.099 |
| Gene subset 4 | ASTRAL | - Plecto | 0.0001 | 0.107 |
| Gene subset 5 | ASTRAL | - Plecto | 0.0001 | 0.190 |
| Gene subset 6 | ASTRAL | - Plecto | 0.0032 | 0.060 |
| Gene subset 7 | ASTRAL | - Plecto | 0.0001 | 0.046 |
| Gene subset 8 | ASTRAL | - Plecto | 0.0001 | 0.054 |
| Gene subset 9 | ASTRAL | - Plecto | 0.0001 | 0.051 |
| Gene subset 10 | ASTRAL | - Plecto | 0.0001 | 0.064 |
| Gene subset 11 | ASTRAL | - Plecto | 0.0001 | 0.140 |
| Gene subset 1 | IQ-TREE | +Plecto | 0.0001 | 0.060 |
| Gene subset 2 | IQ-TREE | +Plecto | 0.0008 | 0.045 |
| Gene subset 3 | IQ-TREE | +Plecto | 0.0014 | 0.040 |
| Gene subset 4 | IQ-TREE | +Plecto | 0.0002 | 0.048 |
| Gene subset 5 | IQ-TREE | +Plecto | 0.0012 | 0.048 |
| Gene subset 6 | IQ-TREE | +Plecto | 0.0001 | 0.133 |
| Gene subset 7 | IQ-TREE | +Plecto | 0.0005 | 0.036 |
| Gene subset 8 | IQ-TREE | +Plecto | 0.0001 | 0.102 |
| Gene subset 9 | IQ-TREE | +Plecto | 0.0004 | 0.053 |
| Gene subset 10 | IQ-TREE | +Plecto | 0.0026 | 0.022 |
| Gene subset 1 | IQ-TREE | - Plecto | 0.0001 | 0.080 |
| Gene subset 2 | IQ-TREE | - Plecto | 0.0032 | 0.086 |
| Gene subset 3 | IQ-TREE | - Plecto | 0.0001 | 0.069 |
| Gene subset 4 | IQ-TREE | - Plecto | 0.0001 | 0.045 |
| Gene subset 5 | IQ-TREE | - Plecto | 0.0005 | 0.045 |
| Gene subset 6 | IQ-TREE | - Plecto | 0.0001 | 0.086 |
| Gene subset 7 | IQ-TREE | - Plecto | 0.0001 | 0.052 |
| Gene subset 8 | IQ-TREE | - Plecto | 0.0001 | 0.059 |
| Gene subset 9 | IQ-TREE | - Plecto | 0.0001 | 0.046 |

| | | | | |
|---|---|---|---|---|
| Gene subset 10 | IQ-TREE | - Plecto | 0.0001 | 0.056 |

**Table S8.** Pairwise comparison of shape disparity among habitats.

**Table S8A.** Procrustes variances for defined groups

| | Brackish | Coastal demersal | Deep-sea demersal | Deep-sea pelagic | Freshwater | Shallow marine pelagic |
|---|---|---|---|---|---|---|
| Procrustes variance | 0.0020 | 0.0025 | 0.0020 | 0.0022 | 0.0010 | 0.0021 |

**Table S8B.** Pairwise absolute differences between variances.

| | Brackish | Coastal demersal | Deep-sea demersal | Deep-sea pelagic | Freshwater | Shallow marine pelagic |
|---|---|---|---|---|---|---|
| Brackish | NA | | | | | |
| Coastal demersal | 0.00046 | NA | | | | |
| Deep-sea demersal | 0.00002 | 0.00048 | NA | | | |
| Deep-sea pelagic | 0.00018 | 0.00028 | 0.00020 | NA | | |
| Freshwater | 0.00103 | 0.00149 | 0.00101 | 0.00121 | NA | |
| Shallow marine pelagic | 0.00009 | 0.00037 | 0.00011 | 0.00009 | 0.00112 | NA |

**Table S8C.** Pairwise p-values based on 10,000 permutations. Significant comparisons are in bold.

| | Brackish | Coastal demersal | Deep-sea demersal | Deep-sea pelagic | Freshwater | Shallow marine pelagic |
|---|---|---|---|---|---|---|
| Brackish | NA | | | | | |
| Coastal demersal | **0.025** | NA | | | | |
| Deep-sea demersal | 0.9285 | **0.0007** | NA | | | |
| Deep-sea pelagic | 0.6305 | 0.3907 | 0.5484 | NA | | |
| Freshwater | **0.0003** | **0.0001** | **0.0001** | **0.0014** | NA | |
| Shallow marine pelagic | 0.7878 | 0.1665 | 0.7023 | 0.8229 | **0.0004** | NA |

**Table S9.** Ordered habitat transition model for ancestral state reconstructions in SIMMAP. Cells with a "0" indicate the transition was not allowed. Transitions numbered 1–18 were allowed to take on independent rates.

| | Brackish | Coastal demersal | Deep-sea demersal | Deep-sea pelagic | Freshwater | Shallow marine pelagic |
|---|---|---|---|---|---|---|
| Brackish | NA | 1 | 0 | 0 | 2 | 3 |
| Coastal demersal | 4 | NA | 5 | 0 | 6 | 7 |
| Deep-sea demersal | 0 | 8 | NA | 9 | 0 | 0 |
| Deep-sea pelagic | 0 | 0 | 10 | NA | 0 | 11 |
| Freshwater | 12 | 13 | 0 | 0 | NA | 14 |
| Shallow marine pelagic | 15 | 16 | 0 | 17 | 18 | NA |

**Table S10.** Results of evolutionary convergence analyses for habitat (using the *convevol* package) using the four "all genes" trees as alternative inputs.

| Tree estimation | Fossil calibration scheme | Habitat | Number of independent groups | Metric type | Metric value | P-value |
|---|---|---|---|---|---|---|
| ASTRAL | - Plecto | coastal demersal | 92 | Ct1 | 0.053 | NA |
| | | | | Ct2 | 1.828 | NA |
| | | | | Ct3 | 0.033 | NA |
| | | | | Ct4 | 0 | NA |
| ASTRAL | + Plecto | coastal demersal | 63 | Ct1 | 0.049 | NA |
| | | | | Ct2 | 1.989 | NA |
| | | | | Ct3 | 0.036 | NA |
| | | | | Ct4 | 0 | NA |
| IQ-TREE | - Plecto | coastal demersal | 59 | Ct1 | 0.071 | NA |
| | | | | Ct2 | 2.254 | NA |
| | | | | Ct3 | 0.04 | NA |
| | | | | Ct4 | 0 | NA |
| IQ-TREE | + Plecto | coastal demersal | 90 | Ct1 | 0.055 | NA |
| | | | | Ct2 | 1.764 | NA |
| | | | | Ct3 | 0.031 | NA |
| | | | | Ct4 | 0 | NA |
| ASTRAL | - Plecto | shallow pelagic | 15 | Ct1 | 0.102 | 0.18 |
| | | | | Ct2 | 2.423 | 0.18 |
| | | | | Ct3 | 0.058 | 0.08 |
| | | | | Ct4 | 0.001 | 0.21 |
| ASTRAL | + Plecto | shallow pelagic | 14 | Ct1 | 0.104 | 0.13 |
| | | | | Ct2 | 2.472 | 0.21 |
| | | | | Ct3 | 0.059 | 0.05 |
| | | | | Ct4 | 0.002 | 0.1 |
| IQ-TREE | - Plecto | shallow pelagic | 14 | Ct1 | 0.099 | 0.14 |
| | | | | Ct2 | 2.335 | 0.15 |
| | | | | Ct3 | 0.055 | 0.05 |
| | | | | Ct4 | 0.002 | 0.03 |
| IQ-TREE | + Plecto | shallow pelagic | 14 | Ct1 | 0.101 | 0.15 |
| | | | | Ct2 | 2.344 | 0.2 |
| | | | | Ct3 | 0.055 | 0.08 |
| | | | | Ct4 | 0.002 | 0.08 |
| ASTRAL | - Plecto | deep pelagic | 4 | Ct1 | 0.378 | 0 |
| | | | | Ct2 | 8.481 | 0.01 |

| | | | | | | |
|---|---|---|---|---|---|---|
| | | | | Ct3 | 0.125 | 0 |
| | | | | Ct4 | 0.003 | 0.06 |
| ASTRAL | + Plecto | deep pelagic | 4 | Ct1 | 0.37 | 0.01 |
| | | | | Ct2 | 7.803 | 0.05 |
| | | | | Ct3 | 0.116 | 0.05 |
| | | | | Ct4 | 0.002 | 0.11 |
| IQ-TREE | - Plecto | deep pelagic | 4 | Ct1 | 0.337 | 0 |
| | | | | Ct2 | 6.161 | 0 |
| | | | | Ct3 | 0.088 | 0.04 |
| | | | | Ct4 | 0.001 | 0.13 |
| IQ-TREE | + Plecto | deep pelagic | 4 | Ct1 | 0.344 | 0.04 |
| | | | | Ct2 | 6.672 | 0.06 |
| | | | | Ct3 | 0.094 | 0.08 |
| | | | | Ct4 | 0.001 | 0.12 |
| ASTRAL | - Plecto | deep demersal | 25 | Ct1 | 0.075 | 0.31 |
| | | | | Ct2 | 2.444 | 0.15 |
| | | | | Ct3 | 0.056 | 0.01 |
| | | | | Ct4 | -0.002 | 0.77 |
| ASTRAL | + Plecto | deep demersal | 19 | Ct1 | 0.156 | 0.03 |
| | | | | Ct2 | 2.87 | 0.12 |
| | | | | Ct3 | 0.071 | 0.02 |
| | | | | Ct4 | 0.001 | 0.27 |
| IQ-TREE | - Plecto | deep demersal | 20 | Ct1 | 0.147 | 0.07 |
| | | | | Ct2 | 2.827 | 0.18 |
| | | | | Ct3 | 0.07 | 0.01 |
| | | | | Ct4 | 0.001 | 0.19 |
| IQ-TREE | + Plecto | deep demersal | 27 | Ct1 | 0.058 | 0.3 |
| | | | | Ct2 | 2.336 | 0.16 |
| | | | | Ct3 | 0.052 | 0.05 |
| | | | | Ct4 | -0.001 | 0.52 |
| ASTRAL | - Plecto | brackish | 18 | Ct1 | 0.026 | 0.17 |
| | | | | Ct2 | 0.386 | 0.36 |
| | | | | Ct3 | 0.006 | 0.43 |
| | | | | Ct4 | -0.001 | 0.52 |
| ASTRAL | + Plecto | brackish | 28 | Ct1 | -0.017 | 0.3 |
| | | | | Ct2 | 0.206 | 0.36 |
| | | | | Ct3 | 0 | 0.44 |
| | | | | Ct4 | -0.002 | 0.22 |
| IQ-TREE | - Plecto | brackish | 28 | Ct1 | -0.033 | 0.37 |
| | | | | Ct2 | -0.085 | 0.39 |
| | | | | Ct3 | -0.004 | 0.46 |
| | | | | Ct4 | -0.001 | 0.25 |
| IQ-TREE | + Plecto | brackish | 18 | Ct1 | 0.005 | 0.27 |

| | | | | | | |
|---|---|---|---|---|---|---|
| | | | | Ct2 | -0.031 | 0.41 |
| | | | | Ct3 | -0.002 | 0.46 |
| | | | | Ct4 | -0.001 | 0.4 |
| ASTRAL | - Plecto | freshwater | 9 | Ct1 | 0.039 | 0.43 |
| | | | | Ct2 | 1.641 | 0.41 |
| | | | | Ct3 | 0.039 | 0.24 |
| | | | | Ct4 | 0.001 | 0.16 |
| ASTRAL | + Plecto | freshwater | 10 | Ct1 | -0.003 | 0.54 |
| | | | | Ct2 | 1.329 | 0.37 |
| | | | | Ct3 | 0.032 | 0.2 |
| | | | | Ct4 | -0.003 | 0.25 |
| IQ-TREE | - Plecto | freshwater | 11 | Ct1 | -0.063 | 0.47 |
| | | | | Ct2 | 1.064 | 0.37 |
| | | | | Ct3 | 0.025 | 0.2 |
| | | | | Ct4 | -0.003 | 0.1 |
| IQ-TREE | + Plecto | freshwater | 9 | Ct1 | 0.029 | 0.5 |
| | | | | Ct2 | 1.298 | 0.48 |
| | | | | Ct3 | 0.032 | 0.31 |
| | | | | Ct4 | -0.001 | 0.26 |

**Table S11.** Results of evolutionary convergence analyses for diet (using the *convevol* package) using the four "all genes" trees as alternative inputs.

| Tree estimation | Fossil calibration scheme | Diet | Number of independent groups | Metric type | Metric value | P-value |
|---|---|---|---|---|---|---|
| ASTRAL | - Plecto | piscivore | 51 | Ct1 | 0.048 | 0.01 |
| | | | | Ct2 | 1.62 | 0.05 |
| | | | | Ct3 | 0.029 | 0 |
| | | | | Ct4 | -0.004 | 0.39 |
| ASTRAL | + Plecto | piscivore | 51 | Ct1 | 0.039 | 0.01 |
| | | | | Ct2 | 1.41 | 0.08 |
| | | | | Ct3 | 0.026 | 0.03 |
| | | | | Ct4 | -0.004 | 0.32 |
| IQ-TREE | - Plecto | piscivore | 52 | Ct1 | 0.023 | 0.05 |
| | | | | Ct2 | 0.898 | 0.17 |
| | | | | Ct3 | 0.017 | 0.09 |
| | | | | Ct4 | -0.003 | 0.31 |
| IQ-TREE | + Plecto | piscivore | 53 | Ct1 | 0.029 | 0 |
| | | | | Ct2 | 1.089 | 0.07 |
| | | | | Ct3 | 0.02 | 0.07 |
| | | | | Ct4 | -0.003 | 0.17 |
| ASTRAL | - Plecto | benthivore / invertivore | 19 | Ct1 | 0.087 | 0.01 |
| | | | | Ct2 | 2.304 | 0.06 |
| | | | | Ct3 | 0.047 | 0.01 |
| | | | | Ct4 | 0 | 0.03 |
| ASTRAL | + Plecto | benthivore / invertivore | 19 | Ct1 | 0.091 | NA |
| | | | | Ct2 | 2.433 | NA |
| | | | | Ct3 | 0.046 | NA |
| | | | | Ct4 | 0.001 | NA |
| IQ-TREE | - Plecto | benthivore / invertivore | 13 | Ct1 | 0.092 | 0 |
| | | | | Ct2 | 2.304 | 0.07 |
| | | | | Ct3 | 0.044 | 0.02 |
| | | | | Ct4 | 0 | 0.03 |
| IQ-TREE | + Plecto | benthivore / invertivore | 13 | Ct1 | 0.075 | 0.05 |
| | | | | Ct2 | 1.899 | 0.13 |
| | | | | Ct3 | 0.04 | 0.05 |
| | | | | Ct4 | 0.001 | 0.11 |

| | | | | | | |
|---|---|---|---|---|---|---|
| ASTRAL | - Plecto | durophage | 28 | Ct1<br>Ct2<br>Ct3<br>Ct4 | 0.056<br>2.145<br>0.038<br>0 | NA<br>NA<br>NA<br>NA |
| ASTRAL | + Plecto | durophage | 29 | Ct1<br>Ct2<br>Ct3<br>Ct4 | 0.067<br>2.18<br>0.039<br>0 | NA<br>NA<br>NA<br>NA |
| IQ-TREE | - Plecto | durophage | 28 | Ct1<br>Ct2<br>Ct3<br>Ct4 | 0.044<br>2.163<br>0.037<br>0 | 0.5<br>0.15<br>0.12<br>0.28 |
| IQ-TREE | + Plecto | durophage | 29 | Ct1<br>Ct2<br>Ct3<br>Ct4 | 0.043<br>2.132<br>0.037<br>0 | 0.51<br>0.2<br>0.14<br>0.3 |
| ASTRAL | - Plecto | planktivore | 8 | Ct1<br>Ct2<br>Ct3<br>Ct4 | 0.142<br>3.7<br>0.094<br>0.004 | 0.26<br>0.17<br>0.08<br>0.05 |
| ASTRAL | + Plecto | planktivore | 8 | Ct1<br>Ct2<br>Ct3<br>Ct4 | 0.141<br>3.663<br>0.093<br>0.004 | 0.25<br>0.24<br>0.09<br>0.03 |
| IQ-TREE | - Plecto | planktivore | 8 | Ct1<br>Ct2<br>Ct3<br>Ct4 | 0.152<br>3.84<br>0.092<br>0.004 | 0.22<br>0.18<br>0.07<br>0.07 |
| IQ-TREE | + Plecto | planktivore | 8 | Ct1<br>Ct2<br>Ct3<br>Ct4 | 0.152<br>3.793<br>0.091<br>0.004 | 0.21<br>0.17<br>0.06<br>0.09 |
| ASTRAL | - Plecto | herbivore / detritivore | 8 | Ct1<br>Ct2<br>Ct3<br>Ct4 | 0.241<br>4.274<br>0.066<br>0.003 | 0.06<br>0.1<br>0.16<br>0.05 |
| ASTRAL | + Plecto | herbivore / detritivore | 8 | Ct1<br>Ct2<br>Ct3<br>Ct4 | 0.245<br>4.354<br>0.068<br>0.003 | 0.07<br>0.13<br>0.17<br>0.1 |

| | | | | | | |
|---|---|---|---|---|---|---|
| IQ-TREE | - Plecto | herbivore / detritivore | 8 | Ct1<br>Ct2<br>Ct3<br>Ct4 | 0.258<br>4.639<br>0.07<br>0.003 | 0.06<br>0.09<br>0.15<br>0.05 |
| IQ-TREE | + Plecto | herbivore / detritivore | 8 | Ct1<br>Ct2<br>Ct3<br>Ct4 | 0.256<br>4.522<br>0.068<br>0.003 | 0.01<br>0.12<br>0.14<br>0.04 |
